# Illuminating the mystery of thylacine extinction: a role for relaxed selection and gene loss

**DOI:** 10.1101/2025.03.27.645633

**Authors:** Buddhabhushan Girish Salve, Nagarjun Vijay

## Abstract

Gene loss shapes lineage-specific traits but is often overlooked in species survival. In this study, we investigate the role of ancestral gene loss using the extinction icon─thylacine (*Thylacinus cynocephalus*). While studies of neutral genetic variation indicate a population decline before extinction, the impact of thylacine-specific ancestral gene losses remains unexplored. The availability of a chromosomal-level genome of the extinct thylacine offers a unique opportunity for such comparative studies. Here, we leverage palaeogenomic data to compare gene presence/absence patterns between the Tasmanian devil and thylacine. We discovered ancestral (between 13-1 Ma) loss of *SAMD9L*, *HSD17B13*, *CUZD1* and *VWA7* due to multiple gene-inactivating mutations, corroborated by short-read sequencing. The loss of *SAMD9* correlates with a carnivorous diet, with ancestral gene loss timing mirroring the thylacine’s shift to hypercarnivory. Our genome-wide analysis reveals olfactory receptor loss and relaxed selection, aligning with reduced olfactory lobes in the thylacine, indicating olfaction is not its primary hunting sense. By integrating palaeogenomic data with comparative genomics, our study reveals ancestral gene losses and their impact on species fitness while providing a foundation for selecting candidate species and genes for thylacine de-extinction. Our approach can be extended to other extinct/endangered species, highlighting genetic factors for de-extinction/conservation efforts.

## 1) Introduction

Traditional genetics-based approaches to species conservation have focused mainly on measures of genetic diversity, the prevalence of mutational load and recent demographic histories [1–4]. However, there has been little focus on ancestral gene loss and its potential role in species survival and extinction. This aspect remains underexplored in conservation genetics despite its potential impact on long-term species survival and resilience to rapid environmental changes due to anthropogenic activities. Several studies have shown that gene loss plays a key role in shaping phenotypic evolution [5], such as adaptations for blood-feeding in bats [6], modifications of the epidermis for aquatic environments in cetaceans [7], dietary variation and retention/pseudogenisation of *ADH7* [4] and loss of the *PLAAT1* gene in vertebrates for adaptation to low-light environments [8]. Nonetheless, ancestral gene losses have demonstrated implications for species survival. For instance, (i) the ancient loss of the *PON1* gene in marine mammals eliminates their primary defence against neurotoxicity from anthropogenic organophosphorus compounds, posing potential risks in modern environments [9], (ii) in odontocetes, the loss of *MX1* and *MX2* genes leads to decreased resistance to viral infections and long-term survival [10], and (iii) the loss of the *CNR2* gene makes parrots more susceptible to neuroinflammation [11], highlighting the consequences of gene loss in these species in terms of long-term survival and disease susceptibility. In previous studies, we highlighted that the loss of ciliary and immune cell genes contributes to differential susceptibility to influenza in bird species [12,13]. These findings emphasise that species-specific ancestral gene losses, which may have conferred advantages in adapting to unique environments in the distant past, can now have adverse effects in the present day because of anthropogenic activities [7] and underscore a critical link between gene loss and increased extinction risk.

As a promising approach, knowledge of gene losses in extinct species could provide valuable insights into the causes of species extinction, refine de-extinction strategies, and offer clues about evolutionary processes, such as trait evolvability. For example, the loss of the *FOXQ1* gene among extinct woolly mammoths has been linked to a reduced ability to adapt to cold climates, ultimately leading to lower fitness [14]. The advent of machine learning-based comparative genomics tools such as TOGA [15] and the increase in publicly available genomic data from both extant and extinct species make it possible to conduct genome-wide screening for ancestral gene loss in extinct species [15–17]. Gene loss can result from genetic drift, positive or relaxed selection, leading to exon deletions or the accumulation of mutations that disrupt the reading frame, such as premature stop codons, frameshifts, dysfunctional splice sites, or repeat insertions or chromosomal rearrangements [6,8,12,13]. The scarcity of multiple samples from extinct species hampers the robust estimation of genetic diversity and the prevalence of mutational load, making it challenging to fully understand their genetic dynamics and evolutionary history [18]. However, single-sample genomic data are sufficient for ancestral gene loss inference. Genome-wide screening for ancestral gene loss becomes vital when the only surviving species within a clade becomes extinct. For example, the thylacine (*Thylacinus cynocephalus*; also known as the Tasmanian tiger or Tasmanian wolf) was the largest modern-day carnivorous marsupial and the sole surviving member of the genus *Thylacinus* and the family *Thylacinidae* until modern times [19]. The *Thylacinidae* were small-bodied, generalist faunivores until the middle Miocene climatic transition (MMCT; ∼15–13 Ma). Following this period, they increased in size and specialised in a hypercarnivorous diet [20]. This raises a critical question in conservation genetics: which genetic changes are unique to a species, and what are their functional impacts, particularly in terms of the species’ long-term survivability, resilience to environmental changes, and vulnerability to novel diseases?

Thylacines were once widespread across continental Australia and New Guinea, but their disappearance from the mainland during the late Holocene was likely driven by climate change, human intensification, and competition with dingoes [21–23]. In Tasmania, however, thylacine numbers had already declined significantly by the early 1800s, following European settlement, the introduction of farming, a government hunting bounty, and disease, leading to eventual extinction, with the last recorded thylacine dying in 1936 [16,17,24]. However, mitochondrial and genomic DNA sequences suggest that limited genetic diversity in thylacine precedes climatic transition-mediated extinction [17,25]. Previous research has focused primarily on thylacine’s canid-like craniofacial convergence, estimation of extinction date, phylogenetic relationships, demographic history, annotation of immune genes, genetic diversity, biology and anatomy [16,17,19,22,25–34].

In this study, we leverage paleogenomic data of thylacine to examine gene presence and absence patterns compared with the Tasmanian devil (*Sarcophilus harrisii*). We began with a genome-wide screen for gene losses in the thylacine, corroborating these loss events with short-read Illumina sequencing data. Additionally, we reconstructed the gene loss history using genomes from other marsupial species and assessed relaxed selection signatures. Our results offer novel insights into the ancestral loss of key genes in the thylacine─ *SAMD9L*, *HSD17B13*, *CUZD1* and *VWA7* potentially compromising its health by affecting antiviral defence, metabolic processes, lactation, pancreatitis and tumour susceptibility, which may have influenced its extinction and environmental adaptations, providing a basis for refining genomics-informed genetic-engineering based de-extinction strategies. Although the present study focuses on the thylacine, our approach can easily be extended to other species (extinct and extant) and improve our understanding of species-specific biology, thereby refining de-extinction strategies and extending conservation efforts to include genetic-engineering-based enhancements.

## 2) Methods

### (a) Genome-wide screen for thylacine-specific gene loss

We identified gene loss in the thylacine (*Thylacinus cynocephalus*) by comparing its genome to the Tasmanian devil (*Sarcophilus harrisii*). Firstly, to compute pairwise genome alignments between the Tasmanian devil (GCF_902635505.1; chromosome-wise; as target) and the thylacine (GCA_007646695.3; whole genome; as query), we used the make_lastz_chains v2.0.8 pipeline (https://github.com/hillerlab/make_lastz_chains), which employs LASTZ v1.04.15 with the following parameters: K = 2400, L = 3000, Y = 9400, H = 2000. Next, we applied the machine learning-based tool TOGA v1.1.7 [15] to determine the status of genes in the thylacine genome, using chain alignment and NCBI RefSeq annotation of the Tasmanian devil (52,692 transcripts) (see electronic supplementary material, figure S1).

### (b) Reconstruction of gene loss history and its validation

We assessed the gene orthology of the gene loss candidates on the basis of their syntenic position within the genomes of human (*Homo sapiens*), opossum (*Monodelphis domestica*), yellow-footed antechinus (*Antechinus flavipes*) and Tasmanian devil (*Sarcophilus harrisii*) (see electronic supplementary material, table S1-S2). The gene-inactivating events inferred by TOGA for “clearly lost” genes were corroborated with the CDS of Tasmanian devil as query using the BLASTn v2.13.0 (parameters: -evalue 0.01 -max_target_seqs 5000 -outfmt ‘17 SQ’) search in ∼223 Gb short-read data of thylacine (SRR5055303, SRR5055304, SRR5055305, and SRR5055306) [35]. The alignments were then used to generate visualizations with IGV-reports v1.12.0, and the results were inspected to confirm the presence of in-frame stop codons and frame-disrupting events. Additionally, we used bam-readcount v1.0.1 [36] to determine the number of reads supporting the reference versus the alternative (disruptive) base or indel at the positions of predicted events by TOGA v1.1.7, providing further support for the inferred gene loss. Additionally, cross-species mapping of thylacine short-read data to Tasmanian devil was done using BWA-mem v0.7.17 [37]. We reassessed the whole-genome sequencing short-read data for the presence of in-frame stop codons using HybPiper v2.3.1 and Patchwork v0.5.2. To assess the integrity of the *HSD17B13* locus in the chromosomal-level assembly of the opossum (*Monodelphis domestica*), we first aligned long-read data of PacBio HiFi to the opossum reference genome using BWA-MEM v0.7.17 [37] (parameters: bwa mem -x pacbio). The resulting BAM file was then used as the input for the scan_alignment and alignment_plot subprograms in the klumpy v1.0.11 [38]. However, similar analyses could not be performed for the numbat (*Myrmecobius fasciatus*), coppery ringtail possum (*Pseudochirops cupreus*), plush-coated ringtail possum (*Pseudochirops corinnae*), and thylacine due to the unavailability of long-read sequencing data for these species.

To reconstruct the history of gene loss, we downloaded available chromosomal-level marsupial genomes from NCBI. In addition, we included scaffold-level assemblies of phylogenetically closely related species, numbat (*Myrmecobius fasciatus*) (n=21; see electronic supplementary material, table S3). We started with a BLASTn v2.13.0 search of marsupial genomes using the Tasmanian devil gene sequences as queries. The resulting hits were processed using mergeBed -d 10000 to merge close intervals and slopBed -b 30000 to expand the regions on both sides. These regions were then extracted using getFastaFromBed of BEDTools v2.27.1 [39]. The resulting FASTA sequences were used to generate pairwise alignments between Tasmanian devil (as target genomes) and twenty marsupial genomes (as query) using make_lastz_chains pipeline, as mentioned in the previous section. We ran TOGA with parameters --o2o to process only the genes with a single orthologous chain, using NCBI RefSeq annotation of the Tasmanian devil.

### (c) Transcriptional status of lost genes

We investigated the RNA expression of the genes lost in thylacine using the publicly available Tasmanian devil and dunnart transcriptome sequencing data from 27 tissues (allantois, amnion, axillary nerve, bone marrow, brachial plexus nerve, brain stem, cerebellum, cerebrum, distal yolk sac, endometrium, eye, heart, liver, lung, optic nerve, ovary, oviduct, pancreas, prostate gland, proximal yolk sac, sciatic nerve, skin, spleen, testis, thyroid, trigeminal nerve and uterus) of both males and females. (BioProject ID: PRJEB34650 [40] and PRJNA1028148 [41]; see electronic supplementary material, table S4). We obtained transcripts per million (TPM) values by performing pseudo-alignments of the reads to the longest isoforms of the Tasmanian devil and dunnart transcripts using kallisto v0.51.0 [42].

We screened the expression of the lost gene in the thylacine miRNA-seq short-read database using BLASTn v2.13.0. To validate the presence of protein-coding gene expression in this dataset, we also queried (as a positive control) the sequences of *LOC100913894* (*ACTA*) and *LOC100925998* (*MYH7*), based on a previous report [43]. Additionally, we assessed the transcriptional status of the *VWA7* gene using RNA-Seq data from the tongue, lung, and liver of the numbat (*Myrmecobius fasciatus*) [44] and *SAMD9*/*9L* using transcriptome data from testes, spleen, lung, liver, kidney and heart of tammar wallaby (*Macropus eugenii*).

### (d) Evaluating the strength of selection, limitations with GC-content and estimation of gene loss timing

To test the relaxation of selection in gene loss candidates, we obtained pairwise codon alignments between the Tasmanian devil and other marsupials, including the thylacine from TOGA [15]. Codons affected by frameshifting insertions or deletions and premature stop codons were replaced by “NNN” to maintain a reading frame. A multiple sequence alignment (MSA) of the protein-coding sequence was done using PRANK v.170427 in GUIDANCE2 v2.01. The sequence saturation in MSA was checked in the DAMBE v7.3.32 program as proposed by Xia et al. [45]. We considered MSA for molecular evolutionary analysis when Iss < Iss.c (Iss.c is the value at which the sequences will begin to fail to recover the true tree) and p-value < 0.05. Furthermore, the alignments and phylogenetic tree obtained from the Timetree website (https://timetree.org/) were used to investigate whether the gene evolves under relaxed selection specifying one branch as the foreground and all other branches as background at a time in the RELAX model of the HyPhy v2.5.48 program [46] and codeml of PAML v4.9f program [47]. We also implemented aBSREL and BUSTED, which are branch and gene-wide tests, respectively, to test for signatures of selection on branches of the marsupial phylogeny for the lost genes. Additionally, site-specific models such as FEL and MEME were used to identify sites evolving under positive and negative selection.

The gene loss timing was estimated using the method proposed by Meredith et al. [49], which is based on two assumptions: (i) dS*f* = dS*p* (the synonymous substitution rate in the mixed branch (dS*f*) does not change after the gene is pseudogenised (dS*p*), i.e. the 1ds method), and (ii) dS*f* = 0.7 dS*p* (the synonymous substitution rate of a functional copy of the gene (dS*f*) is 0.7 times the substitution rate after pseudogenisation (dS*p*), i.e. the 2ds method). We used 38.5 Ma as the divergence time for the thylacine from its most recent common ancestor (MRCA) and dN/dS (ω) estimated for functional (ω*f*), mixed (ω*m*), and pseudogene (ω*p*) using a branch-free model in the codeml program from PAML using unrooted trees.

### (e) *SAMD9* and *SAMD9L* gene phylogeny

To confirm that the *SAMD9* and *SAMD9L* sequences used are 1-to-1 orthologs, we evaluate the maximum likelihood tree generated from marsupial species’ *SAMD9* and *SAMD9L* orthologs. The MSA of *SAMD9*/*9L* was used to generate a gene tree using IQ-TREE v2.3.6 (parameters: *-m MFP --alrt 1000 -B 1000 --boot-trees*) [50]. We also calculated a concordance factor for each branch of the gene tree, comparing the gene tree computed from the complete MSA against the fraction of gene trees of the 1 kb window sequence.

### (f) Identifying *SAMD9* loss and phylogenetic logistic regression

We investigated the status of *SAMD9* and *SAMD9L* genes across mammals using TOGA. To examine whether *SAMD9* gene loss is associated with the percentage of endothermic vertebrates in the diet, we performed phylogenetic logistic regression analyses using the R package “phylolm” [51]. Specifically, we tested whether the proportion of endothermic vertebrates in the diet of each species correlated with the intactness (coded as 1) or loss (coded as 0) of *SAMD9*. We were able to collate the proportion of endothermic vertebrates for a total of 72 mammal species whose *SAMD9* gene status is also available from Jiao, Hengwu et al. [52] and Kapsetaki, Stefania E et al. [53] (see electronic supplementary material, table S5).

For detailed methods and parameter settings, see the electronic supplementary data.

## 3) Results

In brief, our comparative genomic analysis identified ancestral gene losses in the extinct thylacine, attributed to frame-disrupting changes and/or in-frame stop codons in synteny-confirmed orthologs of *SAMD9L*, *HSD17B13*, *CUZD1*, and *VWA7* (figure 1). Signatures of relaxed selection were detected in the thylacine lineage for *SAMD9*/*9L*, *CUZD1*, and *VWA7* genes using RELAX and/or codeml (see electronic supplementary material, table S6). The estimated timing of gene loss for these genes ranges from approximately 13-6 Ma, and the loss of the *SAMD9* gene is negatively correlated with the percentage of endothermic vertebrates in the diet (figure 2; see electronic supplementary material, table S5-S7). Additionally, we observed rampant gene losses in olfactory receptor genes and other genes prone to pseudogenisation accompanied by signatures of relaxed selection (figure 3; see electronic supplementary material, table S1 and S6). The GC content of the four lost genes ranges from 37.65% to 51.71% and is not susceptible to lower coverage due to GC bias. The intermediate levels of GC among these genes suggest these loss events are not a consequence of intragenomic mutational heterogeneity [54].

**Figure 1.**
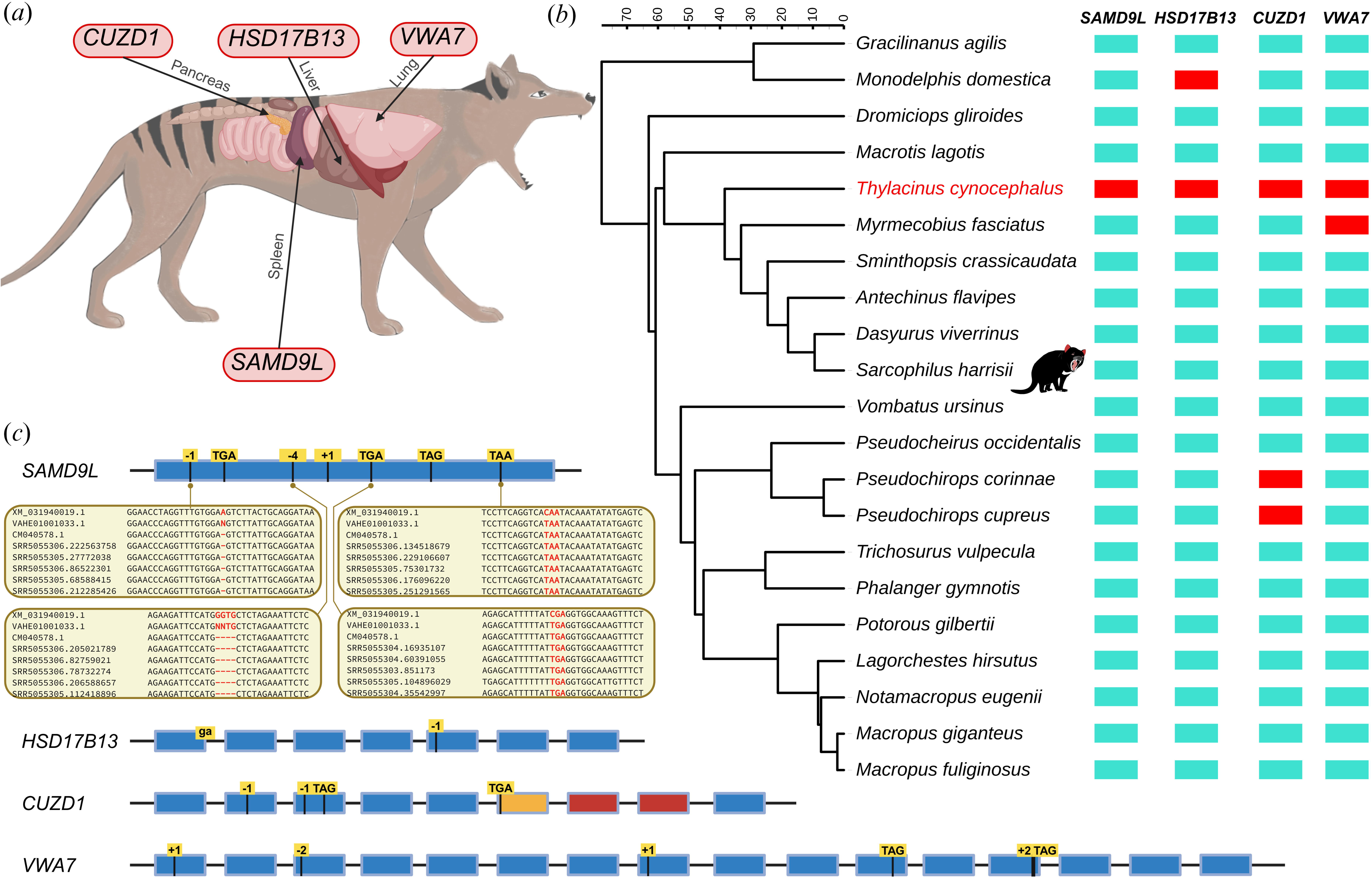
Gene losses in the thylacine (*Thylacinus cynocephalus*). (a) Organs where the lost genes play an important role in the thylacine. We used dog anatomy to illustrate the corresponding organs in the thylacine. (b) Status of the lost genes across 21 marsupial species. A red box indicates gene loss in a given species, while a turquoise box represents an intact gene. The phylogenetic tree used for the analysis was obtained from the TimeTree website (https://timetree.org/) and annotated using iTOL (https://itol.embl.de/). The timescale is in Ma. (c) Frame-disrupting events leading to gene loss. The blue box represents intact exons, while the events of exons affected by frame-disrupting changes are shown with yellow highlights. Exons with no BLASTn hits are shown in red, and those with partial BLASTn hits are depicted in orange. Short-read support is shown for four gene-inactivating events of the *SAMD9L* gene (for all events, see electronic supplementary material, figure S6).

**Figure 2.**
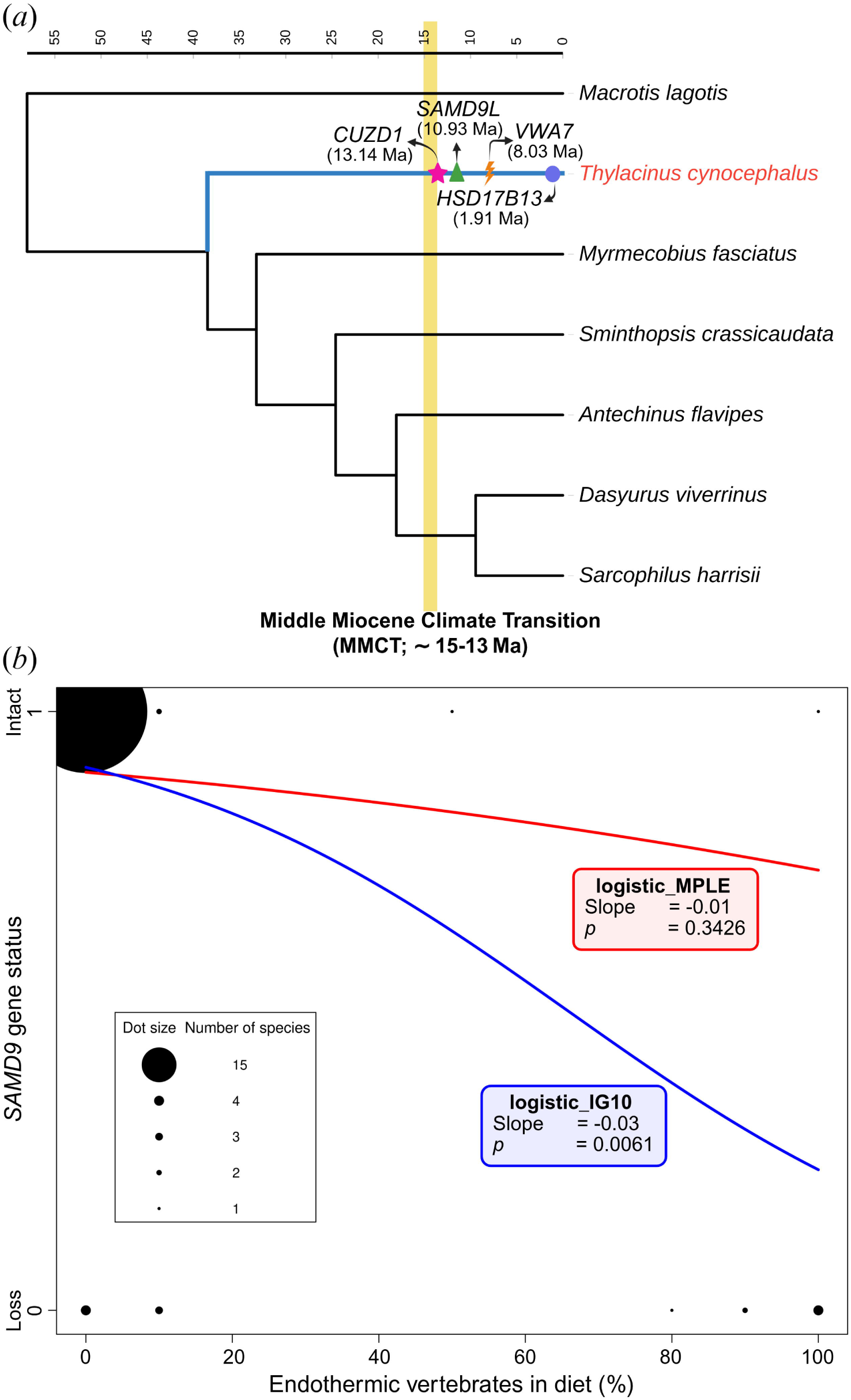
Timing of gene loss in the thylacine mirrors the shift to hypercarnivory. (a) The species tree illustrates the gene loss timing in the thylacine lineage, inferred using the 1ds method under the F1×4 codon frequency model in PAML (see electronic supplementary material, table S7). The mixed branch is coloured in blue, while the functional branches are in black. The yellow-filled rectangle highlights the Middle Miocene Climatic Transition (approximately 15–13 Ma). The timescale is in Ma. (b) The phylogenetic logistic regression finds a negative correlation between the proportion of endothermic vertebrates in the diet and *SAMD9* gene loss in mammals (n = 72). We ran two logistic regression models, the logistic_MPLE (red) and the logistic_IG10 (blue). Diet and genotypes of the species included are shown in black, with the size of the dot indicating sample size.

**Figure 3.**
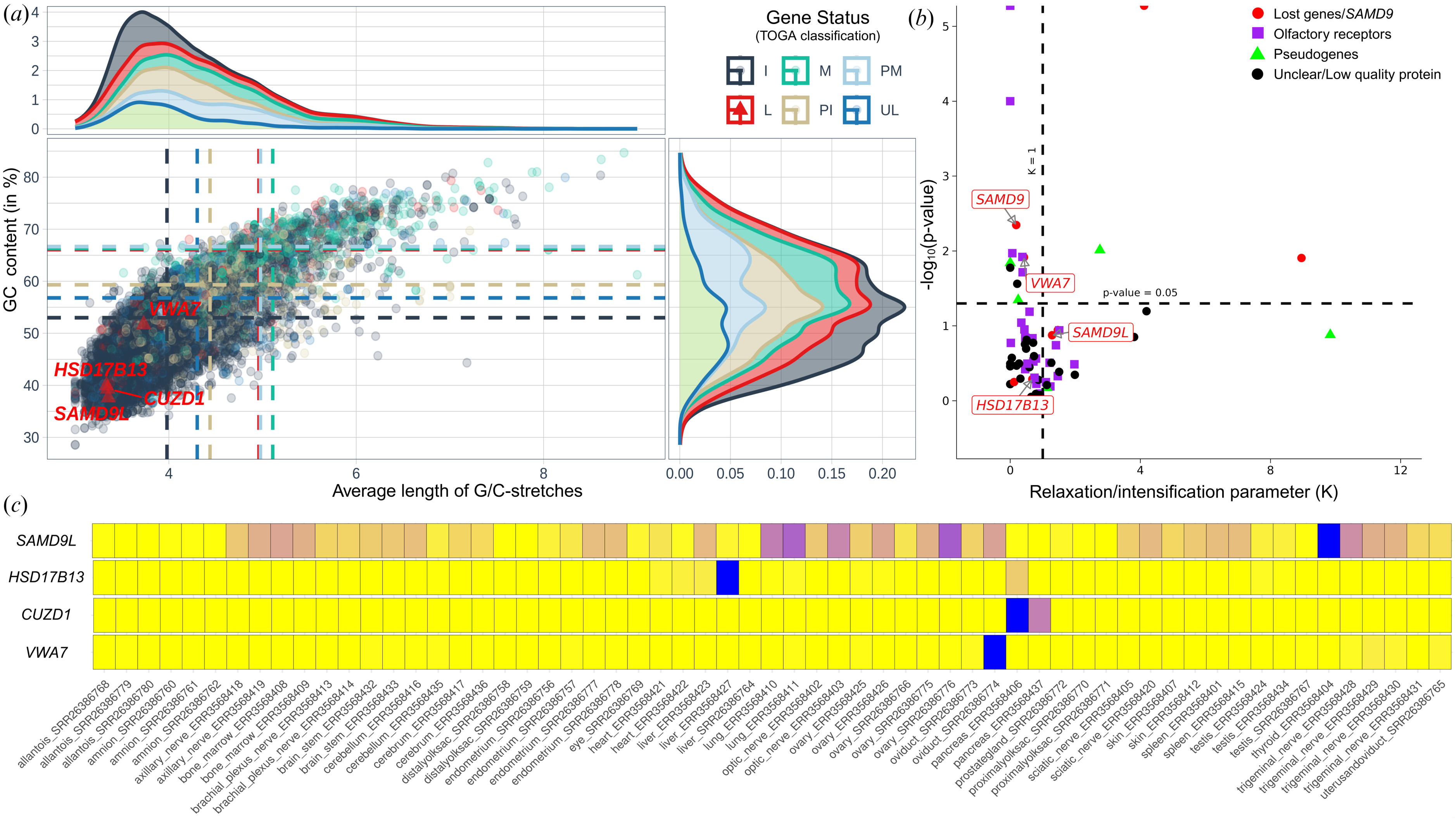
The GC content, relaxed selection signatures, and expression analysis of lost genes. (a) Genes classified by TOGA as missing (M), partially missing (PM), partially intact (PI), unclear loss (UL), and clearly lost (L) exhibit high GC content with the prevalence of long stretches of G/C in the thylacine. Vertical and horizontal dotted lines indicate the third quartile values for a particular classification. The lost genes are marked with red-filled triangles and labelled accordingly. (b) Relaxed selection signatures in the thylacine. The x-axis represents the k-value, and the y-axis represents the -log10(p-value) obtained using the RELAX model of HYPHY (see electronic supplementary material, table S6). (c) Heatmap depicting transcript per million (TPM) values of lost genes across tissues. The tissue names and their corresponding SRR IDs are shown on the x-axis. The colour gradient is from yellow (lowest) to blue (highest). All genes were expressed in at least one tissue above a significant threshold (>1 TPM; see electronic supplementary material, table S4).

### (a) Candidate Gene Losses: Identification and Validation

The thylacine genome was assembled using Illumina sequencing (HiSeq and NextSeq), which exhibits reduced coverage in regions with high GC content [16,17,55]. This limitation can lead to false positives in gene loss identification [56]. In our analysis, many genes categorised by TOGA as missing, partially intact, partially missing, uncertain loss, or clearly lost were associated with high GC content, low or no coverage, missing exons, or deleted exons (figure 3a; see electronic supplementary material, table S1 and figure S1-S2). We filtered out gene loss candidates with high GC content, prevalence of G/C stretches, and insufficient coverage to minimise false positives, ensuring robust identification of true gene losses. Similarly, paralogs can affect the assembly, ultimately leading to ambiguity in validating gene-disrupting events for specific genes (see electronic supplementary material, figure S2-S5 and data). Additionally, genes annotated as “Low-quality protein” in the Tasmanian devil genome were excluded, as they are potentially lost in the Tasmanian devil itself.

Furthermore, we observed widespread gene loss and signatures of relaxed selection in olfactory genes (figure 3b; see electronic supplementary material, table S1 and S6). These loci experience reduced purifying selection and evolve rapidly in many mammals, exhibiting high rates of gene gain and loss [57]. Therefore, we did not pursue these findings further in our study. Also, we observe the loss of genes prone to pseudogenisation (as per NCBI annotation), such as *CASPASE-16*, *PTCHD3*, and *PRSS45* (see electronic supplementary material, table S1 for supporting references).

### (b) Loss of *SAMD9L* and signatures of relaxed selection in *SAMD9*/*9L*

Our genome-wide analysis identified that *SAMD9L* is lost in the thylacine due to multiple gene-inactivating mutations. These mutations include the deletion of one base (A) at codon-287, an in-frame stop codon (CGA→TGA) at codon-511, a deletion of four bases (GGTG) between 1932-1937^th^ position, an insertion of one base (C) at codon-667, and in-frame stop codons at codon-680 (CGA→TGA), codon-1207 (CAG→TAG), and codon-1463 (CAA→TAA) (figure 1). All seven events are supported by short-read sequencing data (see electronic supplementary material, figure S6-S8 and data). Further, HybPiper found non-terminal stop codons for *SAMD9L*, ruling out potential paralog-to-contig assignment errors. Among the screened marsupial genomes, *SAMD9L* loss was unique to the thylacine (figure 1), with an estimated timing of gene loss at approximately 11–8 Ma (figure 2a; see electronic supplementary material, table S7). While other marsupial species such as the opossum, yellow-footed antechinus, and Tasmanian devil have intact *SAMD9L* genes with *TMPPE*, *CRTAP*, and *SUSD5* on the left flank, and *SAMD9*, *CDK6*, *FAM133B*, and *PEX1* on the right flank. In human *CALCR*, *VPS50* and *HEPACAM2* genes are on the left flank of *SAMD9L*, and *SAMD9*, *CDK6*, *FAM133B,* and *PEX1* genes are on the right flank (see electronic supplementary material, table S2). In addition to the synteny-based identification of *SAMD9L* orthologs, gene sequence-based phylogenetic trees also delineated the true orthologs with good bootstrap support (see electronic supplementary material, figure S9-S10). In the Tasmanian devil, *SAMD9L* is annotated as *LOC100924681* (XM_031940019.1), with a GC content of 37.65%. The *SAMD9L* is expressed in the Tasmanian devil and dunnart lung, ovary, and thyroid transcriptomes (figure 3c; see electronic supplementary material, table S4). Notably, we found signatures of relaxed selection using RELAX in *SAMD9* of the thylacine (p-value = 0.004; k-value = 0.16; figure 3b); in contrast, relaxed selection signatures are not found in closely related species of the *Dasyuridae* family. Additionally, we observed relaxed selection signatures for *SAMD9L* using codeml in PAML (see electronic supplementary material, table S6) and episodic diversifying positive selection using aBSREL (p-value = 0.00019) and BUSTED (p-value = 0.0061) models of the Hyphy program in thylacine. Some marsupial species of the *Macropodidae* family also exhibit signatures of relaxed selection (see electronic supplementary material, figure S11 and table S6). However, we found evidence of robust expression in the tammar wallaby (*Macropus eugenii*) (see electronic supplementary material, figure S12).

The signatures of relaxed selection in *SAMD9*/*9L* and *SAMD9L* gene loss due to multiple gene-inactivating mutations indicate that thylacine may have become vulnerable to the critical function provided by *SAMD9* and *SAMD9L* genes. Interestingly, we found that the *SAMD9* gene is annotated as “Low-quality protein” in 22 out of 130 annotated species on NCBI, of which 7/7 are carnivore species. Although most mammals retain an intact *SAMD9*, widespread losses of *SAMD9* were detected in the carnivore lineage. We detected inactivating mutations leading to loss of function, including frameshifting indels and inframe stop codons, in mammalian orders such as Carnivora and Pholidota (see electronic supplementary material, figure S13 and table S5). However, shared loss of *SAMD9* is rare in the Perissodactyla and Artiodactyla orders. Widespread loss of *SAMD9* in carnivores, in contrast to conservation in other vertebrates, suggested a tentative link between diet and dispensability of this gene. We found a negative correlation between the percentage of endothermic vertebrates in the diet and the *SAMD9* gene loss (slopeD= -0.01, p-valueD=D0.3426 for logistic_MPLE method (Insignificant); slopeD=D-0.03, p-valueD=D0.0061 for logistic_IG10 method (Significant), (figure 2b, see electronic supplementary material, table S5)) strengthening the relationship between *SAMD9* dispensability and carnivorous diet.

### (c) Loss of *HSD17B13*

In the thylacine, *HSD17B13* was inactivated through a splice-site mutation (gt →ga) at Exon1, and the deletion of one base (G) in Exon5 (figure 1c; see electronic supplementary material, figure S6). The estimated timing of this gene loss is approximately 2–1 Ma (figure 2a; see electronic supplementary material, table S7). The Tasmanian devil’s *HSD17B13* (XM_003774508.4) has a GC content of 39.98% and is expressed in the liver and pancreas (figure 3c; see electronic supplementary material, table S4). The micro-synteny of the human *HSD17B13* gene is flanked by *IBSP*, *DMP1*, *DSPP*, *SPARCL1*, *SCPPPQ1*, *NUDT9*, and *HSD17B11* on the left, while the right flank consists of *KLHL8*, *AFF1*, *C6H4orf36*, and *SLC10A6* genes. In the yellow-footed antechinus and Tasmanian devil, the left flank includes *IBSP*, *DMP1*, *DSPP*, *NUDT9*, and *HSD17B11*, while the right flank remains consistent with humans, containing *KLHL8*, *AFF1*, *C6H4orf36*, and *SLC10A6,* confirming 1-to-1 orthology (see electronic supplementary material, table S2).

Notably, *HSD17B13* annotation is missing from the annotated genome of the opossum in the NCBI Genome Data Viewer (see electronic supplementary material, figure S14). We found that *HSD17B13* is independently lost in opossum due to an in-frame stop codon (TGG → TGA) in Exon1 (BLASTn search of short-read data revealed polymorphic hits for this event) and deletion of Exons 3-to-7 (figure 1b, see electronic supplementary material, figure S14). We identified an assembly gap at the syntenic locus (see electronic supplementary material, figure S15); however, a BLASTn search in the long-read database returned spurious hits for Exons 3-to-7. Additionally, signatures of relaxed selection were found in the opossum using RELAX (p-value = 0.0008; k-value = 0.00) and PAML’s codeml (see electronic supplementary material, table S6). At the same time, episodic diversifying positive selection was detected in the opossum’s *HSD17B13* using aBSREL (p-value = 0.00) and BUSTED (p-value = 0.00) models (see electronic supplementary material, table S6).

### (d) Loss of *CUZD1*

We identified multiple gene-inactivating mutations in *CUZD1* that contributed to its loss in thylacine (figure 1c). These include the deletion of one base (G) at codons-67 and 95, an in-frame stop codon (TAC → TAG) at codon-119, and another premature stop codon (TGC → TGA) at codon-283 (see electronic supplementary material, figure S6). Furthermore, BLASTn analysis recovered only partial sequences for Exon6, whereas no BLASTn hits were obtained for Exons 7-8, indicating its likely deletion. The estimated timing of gene loss is approximately 13-9 Ma (figure 2a). The *CUZD1* gene, annotated as *LOC100927202* (XM_003755291.4) in the Tasmanian devil, has a GC content of 39.35% and is expressed in the pancreas (figure 3c; see electronic supplementary material, table S4). Notably, we found signatures of relaxed selection by the PAML codeml. At the same time, we found evidence for intensification by RELAX model (p-value = 0.00; k-value = 3.72) and signatures of episodic diversifying positive selection are detected in thylacine *CUZD1* by aBSREL (p-value = 0.00) and BUSTED (p-value = 0.00) models of Hyphy (see electronic supplementary material, table S6). Additionally, *CUZD1* independently lost in the golden ringtail possum (*Pseudochirops corinnae*) and the coppery ringtail possum *(Pseudochirops cupreus*); both species belong to the marsupial family *Pseudocheiridae*. In golden ringtail possum, no BLASTn hits were found for Exons 1-to-4 in the genomic short-read database, while Exon5 exhibited two in-frame stop codons (CAA → TAA and TCA → TGA) and one base (A) deletion. In coppery ringtail possum, an in-frame stop codon (CAA → TAA) and one base (A) deletion led to the inactivation of *CUZD1* (figure 1b; see electronic supplementary material, figure S16). Two gene-inactivating mutations in Exon5 of these two species are shared. At the same time, this gene remains intact in the western ringtail possum (*Pseudocheirus occidentalis*), suggesting that *CUZD1* was lost independently within the *Pseudochirops* genus at approximately 25-20 Ma (figure 1b).

Our micro-syntenic analysis found that in humans, *KZF5*, *PSTK*, *LOC124902519*, *C10orf88B*, and *FAM24B* genes occur on the left flank of *CUZD1* and *LOC124900290*, *DMBT1L1*, and *SPADHI* on the right flank. However, among marsupials (opossum, yellow-footed antechinus and Tasmanian devil), the right flank has *C2H10orf88*, *PSTK,* and *IKZF5,* whereas, on the left flank, *HTRA1* and *DMBT1* were annotated in the NCBI genome data viewer and located approximately 350 kb away supporting 1-to-1 orthology (see electronic supplementary material, table S2).

### (e) Loss of *G7c* (*VWA7*)

This gene has acquired multiple gene-inactivation mutations in thylacine (figure 1c). Insertion of one base (C) in codon-37, two base deletion of CA in codon-171, insertion of one base (T) at codon-403, TGG → TAG premature stop codon at codon-575 and insertion of two bases (TA) leading to an in-frame stop codon (GCC → TAG) at codon-706 (see electronic supplementary material, figure S6). The estimated timing of gene loss is approximately 8-6 Ma (figure 2a). The GC content of Tasmanian devil *VWA7* (XM_003768930.3) is 51.71 % and expressed in the dunnart oviduct (figure 3; see electronic supplementary material, table S4). The micro-synteny of *VWA7* in humans includes *HSPA1A*, *HSPA1L*, *LSM2*, and *VARS1* on the left flank, whereas *SAPCD1*, *MSH5*, *CLIC1*, and *DAH2* are on the right flank. A similar orthology relationship was found in marsupials such as the opossum, yellow-footed antechinus, and Tasmanian devil (see electronic supplementary material, table S2). We found that *VWA7* is lost in numbat due to an in-frame stop codon in Exon5 and Exon14 and the deletion of one base in Exon14 (figure 1, also see electronic supplementary material, figure S17). We found signatures of relaxed selection by RELAX (p-value = 0.01; k-value = 0.40) as well as by codeml in thylacine, and lack of expression at its syntenic locus, suggesting *VWA7* gene loss in numbat (see electronic supplementary material, table S6, and figure S18).

## 4) Discussion

In this study, we used thylacine, an extinction icon, as a case study to investigate ancestral gene losses through a comparative genomic approach. Our genome-wide analysis identified the ancestral loss of *SAMD9L*, *HSD17B13*, *CUZD1*, *VWA7* genes and several olfactory receptors in thylacine. Short-read sequencing confirmed all gene-inactivating mutations, validating gene loss. Our investigation into the timing of gene loss events revealed a staggered pattern, occurring over a span of approximately 13-1 Ma. This temporal distribution indicates that the gene losses were not confined to a single evolutionary event but occurred progressively after the MMCT. Interestingly, the loss of *SAMD9* among mammals is negatively correlated with a carnivorous diet. The genes lost in the thylacine are also lost in about 10% of screened marsupials and show limited evidence of relaxed selection. Relaxation signals detected by RELAX and/or codeml, along with lower dN/dS values on functional branches compared to mixed branches, suggest these genes evolved under gradually relaxed constraints (see electronic supplementary material, table S6-S8) [49,58]. This indicates that their loss was not widespread in marsupials and likely reflects lineage-specific pressures or stochastic events in the thylacine. The lost genes are expressed in similar tissues in the Tasmanian devil and dunnart, implying conserved functions [59] and involved in antiviral defence, metabolism, lactation signalling, pancreatitis, and tumour susceptibility, and their loss may have reduced thylacine fitness and adaptability, potentially contributing to extinction. More broadly, identifying such gene losses informs conservation genomics and de-extinction, guiding genome editing strategies to restore extinct traits. Identifying and targeting these loci can facilitate the selection of appropriate candidate species and genes for genome editing, thereby improving the feasibility and success of de-extinction efforts and genetic engineering in conservation.

### (a) Impact of ancestral gene losses on thylacine fitness and adaptation

The thylacine genome exhibits the loss of four genes– *SAMD9L*, *HSD17B13*, *CUZD1*, and *VWA7* –each with important biological functions and implications for species fitness. The *SAMD9L* (Sterile Alpha Motif Domain Containing 9 Like) is a paralog of *SAMD9* [60]. The *SAMD9*/*9L* genes are critical antiviral factors and tumour suppressors, playing crucial roles in the innate immune defence [61,62]. The *SAMD9* gene is lost (deleted) in the mouse due to evolutionary genomic rearrangements [60], and mouse *SAMD9L* is not a functional paralog of human *SAMD9* [63]. Notably, species-specific differences in *SAMD9L* among rodents act as a natural barrier to cross-species orthopoxvirus infections, underscoring the adaptive significance of these genes in shaping immune defences across different lineages [64]. Furthermore, *SAMD9L* is the only gene that experienced convergent amino acid substitutions and an accelerated evolutionary shift among bats and birds, which are natural hosts of several zoonotic pathogens, highlighting its crucial role in conferring resistance to viruses [65]. Our study found that *SAMD9* and *SAMD9L* are highly conserved across marsupials, except for thylacine. However, *SAMD9* is lost among mammals like carnivores and pangolins while intact in closely related groups such as odd-toed ungulates, indicating a possible association between gene loss and dietary diversification. The phylogenetic correlation between *SAMD9* gene loss and the fraction of endothermic vertebrates in the diet suggests that a shift towards carnivory might have favoured the loss of *SAMD9/9L* [20]. Hence, the loss of *SAMD9L* in thylacine may have conferred a benefit following a shift to hypercarnivory in the past. At the same time, in the modern environment, it may have increased vulnerability to zoonotic viral infections and compromised immune responses. Another lost gene, *HSD17B13* (Hydroxysteroid 17-beta-dehydrogenase 13), encodes a hepatic lipid droplet-associated enzyme [66]. Loss-of-function variants in *HSD17B13* are associated with protection against the progression of steatosis to more severe liver conditions [67], suggesting that *HSD17B13* gene loss could have functional adaptive significance. The CUB and zona pellucida-like domain-containing protein 1 (*CUZD1*) is also lost in the thylacine. *CUZD1* interacts with *JAK1*/*JAK2* and *STAT5*, which are mediators of prolactin signalling critical for mammary gland development. Its absence abolishes *STAT5* phosphorylation during amelogenesis, leading to defective lactation [68]. Furthermore, *CUZD1* is abundantly expressed in pancreas acinar cells of human and mouse [69,70]. We also found that *CUZD1* is expressed in the Tasmanian devil’s pancreas. *CUZD1* deficient mice demonstrated increased severity of experimentally induced acute pancreatitis, suggesting *CUZD1* is a strong candidate as a pancreatitis susceptibility gene [70,71]. Finally, *VWA7* (Von Willebrand Factor A Domain Containing 7), also known as *G7c*, is located in the mouse and human MHC class III region [72]. The *VWA7* gene has been implicated in controlling lung tumour susceptibility and host immune response to viral infections, suggesting its potential role in disease susceptibility [73,74].

Interestingly, the timing of gene losses for *SAMD9L*, *CUZD1*, and *VWA7* is estimated to be approximately 13–6 Ma, a period that follows the middle Miocene climatic transition (MMCT; ∼15–13 Ma). This period marked significant ecological and climatic changes, during which *Thylacinidae*, previously small-bodied, unspecialised faunivores, began to undergo notable shifts. After MMCT, they increased in size and developed adaptations to a hypercarnivorous diet, likely driven by the aridification of the Australian environment [20]. Besides this, the thylacine and grey wolf exemplify remarkable convergent evolution, displaying strikingly similar craniofacial morphologies despite ∼160 million years of evolutionary divergence [16,17]. Notably, these lost genes are located near a candidate craniofacial cis-regulatory element, indicating a possible connection with phenotypic convergence (see supplementary table of [79]). This pattern underscores the complexity of gene loss events and their potential role in shaping convergent phenotypic traits, which remains to be verified experimentally [80]. In addition, our analysis revealed a drastic loss of olfactory receptors in the thylacine, paired with relaxed selection, aligning with earlier findings of its reduced olfactory lobes [20,77]. Unlike the Tasmanian Devil, a scavenger, the taller thylacine likely depended on vision and sound for hunting, aided by its ability to see over vegetation. This shift in sensory reliance is further evidenced by the pronounced ridges on its neocortex, suggesting heightened cognitive abilities to support its hunting strategies [78]. Notably, gene loss has also been observed during major evolutionary transitions, such as from unicellular to multicellular life and from land to water [75,76].

Whatever the cause for gene loss, the loss of *SAMD9L*, *HSD17B13*, *CUZD1*, and *VWA7* genes in the thylacine possibly had negative pleiotropic effects, potentially compromising its health by affecting antiviral defence, metabolic processes, lactation, pancreatitis and tumour susceptibility. Although model-based population viability analyses suggest that the disease played only a minor role in thylacine extinction [81], other studies suggest that a ‘canine-distemper-like’ disease played a role in exacerbating its extinction [82–84]. Therefore, adaptive ancestral gene loss may have provided specific advantages to the thylacine in response to environmental pressures during the MMCT, while being deleterious in the recent past, similar to the example of *PON1* gene loss in marine mammals [9].

### (b) Studying gene loss in extinct species: Opportunities, constraints, and conservation implications

Comparative studies of extinct species offer valuable insights into host-specific immune responses and phenotypic evolution [5,12]. Such studies can play a pivotal role in refining de-extinction strategies by assisting in selecting candidate species and genetic loci for genome editing, thereby informing decisions about diet and habitat preferences [85–87]. Advances in comparative genomics and ancient DNA sequencing enable researchers to link gene loss with phenotypic diversity and evolutionary mechanisms, though such analyses depend heavily on genome assembly quality [6,7,9–11,14]. For extinct species, reliance on Illumina sequencing—despite its limitations in GC-rich regions—can lead to false positives in gene loss detection [16,56]. The fragmentary nature of ancient DNA in such species further restricts the use of long-read sequencing technologies [88]. This lack of long-read sequencing, DNA damage and the absence of transcriptomic data make it even more challenging to differentiate between assembly artefacts and true gene losses [14,16].

Specifically for thylacine, the genome assembly (GCA_007646695.3) was assembled using Illumina HiSeq and NextSeq platforms. In line with previous research, our study finds that genes with high GC content tend to have lower sequencing coverage [56] and, therefore, are classified as missing, partially intact, or partially lost by TOGA. As a result, this might lead to incomplete or less reliable assemblies in GC-rich genomic regions, affecting the accuracy of downstream analyses and may lead to false positives for gene loss [56]. Although miRNA-Seq data is available [43], it does not provide insights into protein-coding gene expression. The gene losses we identified involve gene-inactivating mutations (e.g., splice-site changes, stop codons, insertions, and deletions) supported by forward and reverse short reads, indicating they are genuine rather than artefacts of ancient DNA damage. Assessment of gene loss in large multi-gene families with complex orthology relationships remains challenging and must be interpreted cautiously [6], especially in reference-guided assemblies [89]. Future improvements to the thylacine genome assembly will strengthen the assertion of gene losses. Although habitat loss, hunting, introduced species, inbreeding depression, and loss of genetic diversity are widely recognised as key contributors to species extinction [2,90], these factors have primarily dominated conservation genetics studies focused on mitigating risks to threatened species [3,91–93]. In contrast, the role of ancestral gene loss in species extinction remains largely unexplored and raises the question whether genetically altering a gene or gene combination will help the species survive in the modern world, and how the genetic background (the allelic states of genes, prevalence of regulatory elements, gene content, etc, in the rest of the genome) will affect the fitness [9–11]. Insights into ancestral lineage-specific gene losses can deepen our understanding of extinct species evolution and refine de-extinction approaches. Our study also offers a framework to test hypotheses on why specialist phylogenetic lineages struggle to persist and diversify [7,94] and provides a novel perspective on the interplay between species diversity and genetic diversity.

## 5) Conclusion

Our study offers an immediate framework that includes practical approaches in comparative genomics and insights into de-extinction strategies. Our findings from this case study of an extinction icon-the thylacine─establishes the ancestral gene loss of *SAMD9L*, *HSD17B13*, *CUZD1*, and *VWA7,* contributing unique insights into the genetic factors influencing extinction. Additionally, we observe gene loss in olfactory receptors.

In line with temporal estimations of gene losses, our study provides an empirical case where we observe the coincidence of gene loss timing (approximately 13–1 Ma, following the middle Miocene climatic transition ∼15–13 Ma). This period marked significant ecological and climatic changes, during which *Thylacinidae*, previously small-bodied, unspecialised faunivores, underwent notable shifts such as an increase in size and developing adaptations to a hypercarnivorous diet, likely driven by the aridification of the Australian environment [20]. We find evidence that *SAMD9* gene loss is associated with a switch to hypercarnivory. As the thylacine was the only surviving member of the genus *Thylacinus* and the family *Thylacinidae* until modern times, our study also offers a hypothesis on why specialist phylogenetic lineages struggle to persist and diversify over macroevolutionary timescales [7,19,94]. Furthermore, our analysis reveals a pervasive loss of olfactory receptors in the thylacine, reflecting relaxed selection and correlating with its reduced olfactory lobes. This shift in sensory reliance, possibly driven by changes in traits such as height and vision, contrasts with that of the Tasmanian devil.

In addition, our analysis provides critical information for identifying which species are better candidates for cloning and which genetic loci are suitable for genome editing, contributing to future de-extinction efforts for thylacine and guiding decisions about diet and habitat preferences [85–87]. Our study provides a framework for identifying ancestral gene losses that may improve the probability of success of de-extinction programs.

## Ethics

This work did not require ethical approval from a human subject or animal welfare committee.

## Data accessibility

This study’s data and relevant code are available on Github: https://github.com/CEGLAB-Buddhabhushan/Thylacine_Genome-wide_GENELOSS.git. The reviewer URL: https://github.com/CEGLAB-Buddhabhushan/Thylacine_Genome-wide_GENELOSS.git

## Declaration of AI use

We have not used AI-assisted technologies to create this article.

## Authors’ contributions

B.G.S.: conceptualisation, data curation, formal analysis, investigation, methodology, validation, visualisation, writing—original draft, writing—review and editing; N.V.: conceptualisation, funding acquisition, supervision, investigation, methodology, validation, writing—original draft, writing — review, and editing.

Both authors gave final approval for publication and agreed to be held accountable for the work performed therein.

## Acknowledgements

We thank the Council of Scientific & Industrial Research for a fellowship to B.G.S. We used BioRender (https://biorender.com) to arrange figures and images of species. We thank Sandhya Sharma for the animal paintings in Figure 1.

