## Supplementary Data for "Illuminating the mystery of thylacine extinction: a role for relaxed selection and gene loss"

### Detailed methods, settings and code used

| Sr. No. | Tool | Version | Purpose |
|---|---|---|---|
| 1 | LASTZ (called from make_lastz_chains pipeline) | v1.04.15 | Whole-genome alignment |
| 2 | TOGA | v1.1.7 | Gene loss inference |
| 3 | SAMtools | v1.15.1 | BAM/SAM manipulation |
| 4 | BLASTn | v2.13.0 | Sequence similarity search |
| 5 | IGV-report | v1.12.0 | Visual variant inspection |
| 6 | BWA-MEM | v0.7.17 | Mapping of whole genome sequencing reads |
| 7 | minimap2 | v2.26 | |
| 8 | bowtie2 | v2.5.4 | |
| 9 | bam-readcount | v1.0.1 | Quantifying read support for events |
| 10 | efetch/ esearch | v12 | Extract accession-specific details/data from NCBI |
| 11 | Seqkit | v2.2.0 | Toolkit for FASTA/Q file manipulation |
| 12 | pilon | v1.24 | Genome polishing using raw read data |
| 13 | HybPiper | v2.3.1 | Recover gene sequence from WGS and infers in-frame stop codons |
| 14 | Patchwork | v0.5.2 | Recover gene sequence from WGS |
| 15 | seqtk | v1.2 | Toolkit for FASTA/Q file manipulation |
| 16 | NCBI datasets | 13.43.2 | A command-line tool that is used to query and download biological sequence data across all domains of life from NCBI databases |
| 17 | BEDTools | v2.27.1 | Genomic interval operations |
| 18 | klumpy | v1.0.11 | A tool to evaluate the integrity of long-read genome assemblies and illusive sequence motifs |
| 19 | Kallisto | v0.51.0 | Program for quantifying abundances of transcripts from RNA-Seq data |
| 20 | STAR | v2.7.0d | Program for splice-aware mapping of RNA-seq reads to the genome |
| 21 | PRANK | v.170427 | Multiple sequence alignment tool |

| | | | |
|---|---|---|---|
| 22 | GUIDANCE2 | v2.01 | Suit for codon-aware alignment with bootstrap support |
| 23 | HYPHY | 2.5.48(MP) and 2.5.62(MP) | Selection analysis tools |
| 24 | PAML (codeml) | v4.9f | dN/dS estimation across branches |
| 25 | MapNH | v1.3.0 | GC biased gene conversion |
| 26 | phastBias | v1.6 | GC biased gene conversion |
| 27 | IQ-TREE | v2.3.6 | Phylogenetic analysis |

**The step-by-step commands, parameters used, input files and output files are provided below. The shell scripts are light green, the Python code is light yellow, and the R code is sky blue.**

1. **Genome-wide screening of thylacine-specific gene loss**
   a. **Genome alignment**
   To identify gene loss candidates in the thylacine (*Thylacinus cynocephalus*; as query), we performed pairwise whole-genome alignment with the Tasmanian devil (*Sarcophilus harrisii*; GCF_902635505.1; as reference) using the make_lastz_chains v2.0.8 pipeline (https://github.com/hillerlab/make_lastz_chains). Alignments were generated with LASTZ v1.04.15 [1] using parameters K=2400, L=3000, Y=9400, and H=2000, with the default scoring matrix. Alignments were post-processed using axtChain [2], chainCleaner [3], and RepeatFiller [4] (all with default settings). Output from the make chains pipeline produced pairwise chain-format genome alignments. The code used to generate the chain file is given below.

```
chr=$1
cd $chr
for i in "$chr"_chr.fa
do
sed -i 's/ .*//;s/\./_/' $i
target_name=`echo Tasmanian_devil_"$chr"`
query_name=Tasmanian_wolf
/media/morpheus/sagar/BUDDHA/TOGA_new/make_lastz_chains/make_chains.py
$target_name $query_name $i
../GCA_007646695.3_UniMelb_ThyCyn2.0_hybrid_assembly_genomic.fna --pd
Chain_"$target_name"_"$query_name" -f --chaining_memory 30
done
```

   b. **Screening of gene loss**
   Gene status in the thylacine was evaluated using TOGA v1.1.7 [5]. We used the Genome sequence of Tasmanian devil and thylacine in 2Bit format, along with genome alignment in chain format and Tasmanian devil RefSeq annotations (52692 transcripts) in BED12 format (see electronic supplementary material, figure S1). To obtain the gene model in BED12 format from a GTF file, we used the gtf2bed option in ea-utils (https://expressionanalysis.github.io/ea-utils/). The code used to identify gene loss candidates is given below.

```
chr=$1
cd $chr
```

```
for i in "$chr"_chr.fa
do
target_name=`echo Tasmanian_devil_"$chr"`
faToTwoBit "$chr"_chr.fa "$chr"_chr.2bit
/media/morpheus/sagar/BUDDHA/TOGA/toga.py
./Chain_"$target_name"_Tasmanian_wolf/"$target_name".Tasmanian_wolf.final.chain.gz
GCF_902635505.1_mSarHar1.11_genomic."$chr"_chr.bed12 "$chr"_chr.2bit
../Tasmanian_wolf.2bit --kt --pn TOGA_"$target_name"_Tasmanian_wolf --nc
/media/morpheus/sagar/BUDDHA/TOGA/nextflow_config_files/ --cb 3,5,15,25,50 --cjn
500 --nb
/media/morpheus/sagar/BUDDHA/TOGA/nextflow_config_files/cesar_bigmem_config.nf --
ces --chain_jobs_num 60
done
cd ..
```

We obtained gene loss candidates classified as L (clearly lost) in the output file of TOGA v1.1.7 [5], *loss_summ_data.tsv*. The event of gene loss was extracted from *inact_mut_data.txt*.

```
# collect_toga_inact_mut_data
for i in `seq 1 6` X; do cat
../$i/TOGA_Tasmanian_devil_"$i"_Tasmanian_wolf_o2o/inact_mut_data.txt >>
TOGA.inact_mut_data.txt; echo $i; done
sed -i 's/_/\./2' TOGA.inact_mut_data.txt
# collect_loss_summ_data
for i in `seq 1 6` X; do grep "TRANSCRIPT"
../$i/TOGA_Tasmanian_devil_"$i"_Tasmanian_wolf_o2o/loss_summ_data.tsv|cut -
f2,3|sed 's/_/\./2' >> TOGA.loss_summ_data.tsv; echo $i; done
```

### 2. Confirmation of gene loss event

#### a. Converting event coordinate to transcript coordinate

We converted the event coordinates for coding DNA sequences (CDS) for easy visualisation of gene loss events by TOGA v1.1.7 [5] and confirmation. The following script was used to perform this task.

```
import pandas as pd
import sys

# Check if transcript_id is provided as an argument
if len(sys.argv) != 2:
    print("Usage: python codon_pos_bed6.py <transcript_id>")
    sys.exit(1)

# Get the transcript_id from the command-line argument
transcript_id = sys.argv[1]

# Load the GTF file
gtf_file =
'/media/morpheus/sagar/BUDDHA/Tasmanian_wolf/Chr_wise/Chromosomes/Final_verificati
on/GCF_902635505.1_mSarHar1.11_genomic.gtf.cds.gtf'
gtf_df = pd.read_csv(gtf_file, sep='\t', comment='#', header=None)

# Assign column names
gtf_df.columns = ['seqname', 'source', 'feature', 'start', 'end', 'score',
'strand', 'frame', 'attribute']

# Extract transcript_id from the attribute column
gtf_df['transcript_id'] = gtf_df['attribute'].str.extract('transcript_id
"([^"]+)"')
```

```python
# Filter the DataFrame for the specific transcript_id
transcript_df = gtf_df[(gtf_df['transcript_id'] == transcript_id) &
(gtf_df['feature'] == 'CDS')]

# Initialize a list to store BED6 format codon positions
bed6_entries = []

# Initialize a variable to track the codon phase shift
current_phase_shift = 0

# Initialize codon counter
codon_counter = 1

# Iterate over each CDS segment and calculate codon positions considering the
phase
for _, row in transcript_df.iterrows():
    seqname = row['seqname']
    start = row['start']
    end = row['end']
    strand = row['strand']
    phase = int(row['frame'])

    # Adjust start position based on the current phase shift
    if strand == '+':
        start += current_phase_shift
        codon_start_positions = range(start, end + 1, 3)
    else:  # For negative strand
        end -= current_phase_shift
        codon_start_positions = range(end, start - 1, -3)

    # Update phase shift for the next CDS
    current_phase_shift = (3 - (end - start + 1) % 3) % 3

    # Create BED6 entries
    for codon_start in codon_start_positions:
        if strand == '+':
            chromStart = codon_start - 1  # BED is 0-based
            chromEnd = codon_start + 3    # codon is 3 bases long
        else:
            chromStart = codon_start - 3
            chromEnd = codon_start

        # Label the codon sequentially
        codon_name = f"codon{codon_counter}"
        codon_counter += 1

        bed6_entries.append([seqname, chromStart, chromEnd, codon_name, 0,
strand])

# Convert to DataFrame for easier handling
bed6_df = pd.DataFrame(bed6_entries, columns=['chrom', 'chromStart', 'chromEnd',
'name', 'score', 'strand'])

# Save the codon positions in BED6 format to a file
output_file = f'{transcript_id}.codon_positions.bed'
bed6_df.to_csv(output_file, sep='\t', header=False, index=False)

print(f"Output saved to {output_file}")
```

```bash
#!/bin/bash
transcriptID=$1
# Input FASTA file
```

```
fasta_file="$transcriptID".fa

# Extract sequence name and length using samtools faidx
sequence_name=$(grep "^>" $fasta_file | sed 's/>//')
sequence_length=$(faidx $fasta_file -i chromsizes | awk '{print $2}')

# Define the output BED6 file
output_file="${sequence_name}.cds_codons.bed"

# Initialize start position
start=0

# Open the output file
echo -n "" > $output_file

# Generate the BED6 file
for (( i=1; i<=$sequence_length; i+=3 ))
do
    end=$((start + 3))
    codon_name="codon$(( (i + 3) / 3 ))"
    echo -e "${sequence_name}\t${start}\t${end}\t${codon_name}\t0\t+" >>
$output_file
    start=$((start + 3))
done

echo "BED6 file generated: $output_file"
```

**b.** BLASTn-based: We corroborated TOGA-inferred gene-inactivating mutations using BLASTn v2.13.0 [6] on ~223 Gb (187.7 G bases) of thylacine short-read data (SRR5055303–SRR5055306) [7]. Hits were manually inspected (in IGV-reports) and confirmed (see electronic supplementary material, figure S7). We used the following code (with detailed BLASTn settings: -task blastn -evalue 0.01 -max_target_seqs 5000, save in output format '17 SQ'). We converted alignments (in SAM format to BAM format with SAMtools VIEW v1.15.1 and sorted BAM alignments using SAMtools SORT [8].

```
blastn -task blastn -evalue 0.01 -max_target_seqs 5000 -db
/media/morpheus/disk1/BUDDHA_merged_data/ThyCyn_db/SRR5055303-6.blastDB.fa -out
"$transcriptID".blastn.DNAseqDB.sam -num_threads 32 -outfmt '17 SQ'  -query
"$transcriptID".fa
sed -i "s/Query_1/$transcriptID/g" "$transcriptID".blastn.DNAseqDB.sam
samtools view -bhS "$transcriptID".blastn.DNAseqDB.sam >
"$transcriptID".blastn.DNAseqDB.bam
samtools sort "$transcriptID".blastn.DNAseqDB.bam -o
"$transcriptID".blastn.DNAseqDB.sorted.bam
samtools index "$transcriptID".blastn.DNAseqDB.sorted.bam
```

The BAM file and the events bed for CDS used to generate the IGV-report v1.12.0 [9].

```
# create IGV report
create_report "$transcriptID".codon_positions.events.bed --standalone --fasta
"$transcriptID".fa --tracks "$transcriptID".blastn.DNAseqDB.sorted.bam
"$transcriptID".codon_positions.events.bed --output
"$transcriptID".BLASTn_IGV.html --info-columns Chromosome Start_position
End_position Event score strand thickStart thickEnd itemRgb --translate-sequence-
track
```

**c. Mapping based**

We performed cross-species mapping of thylacine short read data to Tasmanian devil. We used three read mappers- BWA-MEM v0.7.17 [10], minimap2 [11] and bowtie2 v2.5.4 [12] (bwa mem, minimap2 -ax sr, and bowtie2 with very-sensitive-local flag; see electronic supplementary material, figure S8). Out of these three, BWA-MEM performed best read mapping. See the table below of the percentage of read mapped.

| SRA ID | Percent reads mapped | | |
|---|---|---|---|
| | Bowtie2 | Minimap2 | Bwa mem |
| SRR5055303 | 67.55 | 44.51 | 84.37 |
| SRR5055304 | 68.88 | 45.05 | 83.12 |
| SRR5055305 | 74.89 | 55.87 | 90.35 |
| SRR5055306 | 74.98 | 55.98 | 89.62 |

We saved alignments in BAM format and sorted BAM alignments using SAMtools SORT. SAMtools flagstat is used to get the mapping percentage [8].

```
#mapping with BWA-MEM
bwa index GCF_902635505.1_mSarHar1.11_genomic.fna
for i in *_1.fastq.gz; do j=`echo $i|sed 's/_1/_2/g'`; k=`echo $i|sed
's/_1.fastq.gz//g'`
bwa mem -t 32 GCF_902635505.1_mSarHar1.11_genomic.fna $i $j |samtools sort -@ 32 -
O BAM -o "$k".bwa.bam
samtools index -c "$k".bwa.bam
samtools flagstat -@ 16 "$k".bwa.bam > "$k".bwa.flagstat.out
done
samtools merge -@ 32 SRR5055303-6.bwa.merged.bam *.bwa.bam
samtools sort SRR5055303-6.bwa.merged.bam -@ 24 -o SRR5055303-6.bwa.sorted.bam
samtools index -c -@ 16 SRR5055303-6.bwa.sorted.bam
rm SRR5055303-6.bwa.merged.bam

# mapping with minimap2
for i in *_1.fastq.gz
do
j=`echo $i|sed 's/_1/_2/g'`
k=`echo $i|sed 's/_1.fastq.gz//g'`
minimap2 -t 32 -ax sr GCF_902635505.1_mSarHar1.11_genomic.fna $i $j | samtools
sort -@ 32 -O BAM -o "$k".minimap2.bam
samtools index -@ 16 -c "$k".minimap2.bam
samtools flagstat -@ 16 "$k".minimap2.bam > "$k".minimap2.flagstat.out
done
samtools merge -@ 16 SRR5055303-6.minimap2.merged.bam *.minimap2.bam
samtools sort SRR5055303-6.minimap2.merged.bam -@ 16 -o SRR5055303-
6.minimap2.sorted.bam
samtools index -@ 16 -c SRR5055303-6.minimap2.sorted.bam
rm SRR5055303-6.minimap2.merged.bam *.minimap2.bam

# mapping with Bowtie2
bowtie2-align-s --wrapper basic-0 --very-sensitive-local -p 16 -x
GCF_902635505.1_mSarHar1.11_genomic.fna -1 SRR5055303_1.fastq.gz -2
SRR5055303_2.fastq.gz | samtools sort -@16 -o SRR5055303.bam -
bowtie2-align-s --wrapper basic-0 --very-sensitive-local -p 16 -x
GCF_902635505.1_mSarHar1.11_genomic.fna -1 SRR5055304_1.fastq.gz -2
SRR5055304_2.fastq.gz | samtools sort -@16 -o SRR5055304.bam -
bowtie2-align-s --wrapper basic-0 --very-sensitive-local -p 36 -x
GCF_902635505.1_mSarHar1.11_genomic.fna -1 SRR5055305_1.fastq.gz -2
SRR5055305_2.fastq.gz | samtools sort -@ 36 -o SRR5055305.bam -
bowtie2-align-s --wrapper basic-0 --very-sensitive-local -p 36 -x
GCF_902635505.1_mSarHar1.11_genomic.fna -1 SRR5055306_1.fastq.gz -2
SRR5055306_2.fastq.gz | samtools sort -@ 36 -o SRR5055306.bam -
samtools merge -@ 16 ThyCyn.DNA.bowtie2.merged.bam SRR5055303.bam SRR5055304.bam
SRR5055305.bam SRR5055306.bam
samtools sort -@ 16 -o ThyCyn.DNA.bowtie2.merged.sorted.bam
ThyCyn.DNA.bowtie2.merged.bam
```

Read-level visualisation of loss events was performed using IGV-reports v1.12.0 [9,13].

```
create_report "$transcriptID"_codon_positions.events.bed --standalone --fasta
/home/buddha/work/R1_Thylacine/Thylacine_SRA_DNA/GCF_902635505.1_mSarHar1.11_genom
ic.fna --tracks /home/buddha/work/R1_Thylacine/Thylacine_SRA_DNA/SRR5055303-
6.bwa.sorted.bam /home/buddha/work/R1_Thylacine/Thylacine_SRA_DNA/SRR5055303-
6.minimap2.sorted.bam /home/buddha/work/R1_Thylacine/Thylacine_SRA_DNA/SRR5055303-
6.bowtie2.sorted.bam "$transcriptID"_codon_positions.events.bed
/home/buddha/work/R1_Thylacine/Thylacine_SRA_DNA/GCF_902635505.1_mSarHar1.11_genom
ic.gtf --output "$transcriptID".Genome_IGV.html --info-columns Chromosome
Start_position End_position Event score strand thickStart thickEnd itemRgb --
translate-sequence-track
```

Furthermore, considering more species we performed multiple sequence alignments to validate that gene models (annotation of exon boundaries, isoforms and CDS) are correct, and events are explained parsimoniously. This kind of MSA-based evaluation has been used in several studies [14–17]. The following table lists all the ambiguous alignments due to a specific type of sequence or motif. Irrespective of which of the two possible alignments is used, the event leads to a gene loss.

| **Gene name** | **Event** (see supplementary figure S6) | **Issue** |
|---|---|---|
| *SAMD9L* Thylacine | Event 1 - single base deletion of "A" | Due to the dimer "AA". |
| *SAMD9L* Thylacine | Event 3 - four base deletion of "TGGG" or "GGTG" | Due to the TGGGTG motif. |
| *VWA7* Thylacine | Event 1 - single base insertion of "C" | Within a C-homopolymer. |
| *VWA7* Thylacine | Event 2 - two base deletion of "AC" or "CA" | Due to the ACA motif. |
| *VWA7* Thylacine | Event 5 - two base insertion of "TA" | "TA" appears twice in the Thylacine. |
| *CUZD1* Thylacine | Event 1 - single base deletion of "G" | Due to the dimer "GG". |

**d. Event verification using bam-readcount**

We obtained the base level support, i.e., how many reads support the event by TOGA (query base) and base of reference (ref_base). We used *parse_brc.py* to convert the output of bam-readcount v1.0.1 [18] to TSV format.

```
# Short-read verification using bam-readcount
awk '{print $1"\t"$2"\t"$3"\t"$4; if ($4 ~ /->/) {split($4, a, ":"); split(a[4],
change, "->"); start = $2 + 1; for (i = 1; i <= length(change[1]); i++) {ref_base
= substr(change[1], i, 1); alt_base = substr(change[2], i, 1); print
$1"\t"start"\t"ref_base"\t"alt_base; start++}}}'
"$transcriptID".codon_positions.events.bed | grep -v "STOP" | awk '$3!=$4' | awk
'{if ($4 ~ /Exon/) {split($4, a, ":"); print $1"\t"$2"-"$3"\t"a[3]"\t"a[4]} else
{if ($2 == $3) {print $1"\t"$2"-"$2"\t"$3"\t"$4} else {print $1"\t"$2"-
"$2"\t"$3"\t"$4}}}'|sed 's/\t/,/g' >
"$transcriptID".codon_positions.events.brc.csv
for i in `cat "$transcriptID".codon_positions.events.brc.csv`; do

  file_name=`echo $i | sed 's/,/_/g'`
  chr=`echo $i | cut -f1 -d','`
  start_end=`echo $i | cut -f2 -d','`
  mutation_type=`echo $i | cut -f3 -d','`   # Get the 3rd column (mutation type)
  ref_base=`echo $i | cut -f4 -d','`        # Get the 4th column (ref base)
```

```
  query_base=`echo $i | cut -f5 -d','`      # Get the 5th column (query base)

  if [[ "$mutation_type" != "FS_INS" && "$mutation_type" != "FS_DEL" ]]; then
    # Normal processing if not FS_INS or FS_DEL
    bam-readcount -w0 -f "$transcriptID".fa
"$transcriptID".blastn.DNAseqDB.sorted.bam $chr:$start_end >
"$transcriptID"."$chr":"$start_end".brc.tsv
    python
/media/morpheus/sagar/BUDDHA/Tasmanian_wolf/Chr_wise/Chromosomes/Final_verificatio
n/parse_brc.py "$transcriptID"."$chr":"$start_end".brc.tsv >
"$transcriptID"."$chr":"$start_end".parse_brc.tsv
    event_status=$(awk -v chr="$chr" -v ref_base="$ref_base" -v
query_base="$query_base" 'BEGIN {found=0} $7 >= 5 && !found {print "yes";
found=1}' "$transcriptID"."$chr":"$start_end".parse_brc.tsv)
    echo -e "$i,$event_status"

  else
    # Check for + sign in 4th column if FS_INS or - sign if FS_DEL
    bam-readcount -w0 -f "$transcriptID".fa
"$transcriptID".blastn.DNAseqDB.sorted.bam $chr:$start_end >
"$transcriptID"."$chr":"$start_end".brc.tsv
    python
/media/morpheus/sagar/BUDDHA/Tasmanian_wolf/Chr_wise/Chromosomes/Final_verificatio
n/parse_brc.py "$transcriptID"."$chr":"$start_end".brc.tsv >
"$transcriptID"."$chr":"$start_end".parse_brc.tsv

    if [[ "$mutation_type" == "FS_INS" ]]; then
      # Check for "+" in 4th column of parse_brc.tsv
      if cut -f4 "$transcriptID"."$chr":"$start_end".parse_brc.tsv | grep -q "^+"
; then
        echo -e "$i,yes"
      else
        echo -e "$i,No"
      fi

    elif [[ "$mutation_type" == "FS_DEL" ]]; then
      # Check for "-" in 4th column of parse_brc.tsv
      if cut -f4 "$transcriptID"."$chr":"$start_end".parse_brc.tsv | grep -q "^-";
then
        echo -e "$i,yes"
      else
        echo -e "$i,No"
      fi
    fi
  fi
done|sed 's/,/\t/g'> "$transcriptID".codon_positions.events.TOGA_confirm.tsv
```

**All the above steps can be done using the following script.**

```
#Confirm_Loss.sh
#!/bin/bash
### provide input like ./Confirm_Loss.sh transcriptID
transcriptID=$1

# Download CDS
efetch -db nuccore -id $transcriptID -format fasta_cds_na|cut -f1,2 -d'_'|sed
's/>lcl|/>/g' > "$transcriptID".fa
# Transcript info and preliminary assesment

transcript_info_8c=`esearch -db gene -query $transcriptID | esummary | xtract -
pattern DocumentSummary -element
Id,Name,Description,Chromosome,ChrAccVer,ChrStart,ChrStop,ExonCount`
```

```
orf_status=`perl
/media/morpheus/sagar/BUDDHA/Tasmanian_wolf/Chr_wise/Chromosomes/Final_verificatio
n/ORFvalid.pl "$transcriptID".fa`
if echo "$orf_status" | grep -q "1 out of 1 sequences validated as ORFs."; then
    orf_val="yes"
else
    orf_val="no"
fi
GC_Content=`seqkit fx2tab --name --gc "$transcriptID".fa | awk '{print $2}'`
GC_Stretch=`perl
/media/morpheus/sagar/BUDDHA/Tasmanian_wolf/Chr_wise/Chromosomes/Final_verificatio
n/GC_Stretch_finder.pl "$transcriptID".fa |awk '{print $5}'`

# Convert TOGA evenets
python
/media/morpheus/sagar/BUDDHA/Tasmanian_wolf/Chr_wise/Chromosomes/Final_verificatio
n/make_codon_position_bed.py $transcriptID
grep -w "$transcriptID"
/media/morpheus/sagar/BUDDHA/Tasmanian_wolf/Chr_wise/Chromosomes/Final_verificatio
n/TOGA.inact_mut_data.txt > "$transcriptID".inact_mut_data.txt
# Check for exon missing and deleted
if grep -q "Missing exon" "$transcriptID".inact_mut_data.txt ; then
exon_missing="yes"
else
exon_missing="no"
fi

if grep -q "Deleted exon" "$transcriptID".inact_mut_data.txt ; then
exon_deleted="yes"
else
exon_deleted="no"
fi

if grep -q "SSM" "$transcriptID".inact_mut_data.txt ; then
splice_site_change="yes"
else
splice_site_change="no"
fi

if grep -q "START_MISSING" "$transcriptID".inact_mut_data.txt ; then
start_missing="yes"
else
start_missing="no"
fi

cat "$transcriptID".inact_mut_data.txt |cut -f3-6|grep -v "GENE:"|awk 'NF == 4 &&
$1!=0 {print "Exon"$1":codon"$2":"$3":"$4}' > "$transcriptID".events.bed

bash
/media/morpheus/sagar/BUDDHA/Tasmanian_wolf/Chr_wise/Chromosomes/Final_verificatio
n/generate_codon_bed_for_cds.sh $transcriptID
for i in `cat "$transcriptID".events.bed`; do pos=`echo $i|cut -f2 -d':'`; grep -w
"$pos" "$transcriptID".cds_codons.bed|sed "s/$pos/$i/g"|awk '{print
$0"\t"$2"\t"$3"\t"255","0","0}'  >> "$transcriptID".codon_positions.events.bed;
done

# BLASTn in DNAseqDB
blastn -task blastn -evalue 0.01 -max_target_seqs 5000 -db
/media/morpheus/disk1/BUDDHA_merged_data/ThyCyn_db/SRR5055303-6.blastDB.fa -out
"$transcriptID".blastn.DNAseqDB.sam -num_threads 32 -outfmt '17 SQ'  -query
"$transcriptID".fa
sed -i "s/Query_1/$transcriptID/g" "$transcriptID".blastn.DNAseqDB.sam
```

```
samtools view -bhS "$transcriptID".blastn.DNAseqDB.sam >
"$transcriptID".blastn.DNAseqDB.bam
samtools sort "$transcriptID".blastn.DNAseqDB.bam -o
"$transcriptID".blastn.DNAseqDB.sorted.bam
samtools index "$transcriptID".blastn.DNAseqDB.sorted.bam

# BLASTn in RNAseqDB
blastn -task blastn -evalue 0.01 -max_target_seqs 5000 -db
/media/morpheus/sagar/BUDDHA/Tasmanian_wolf/Chr_wise/Chromosomes/mapping/miRNA_seq
/miRNAblastDB/SRR23147611-6.miRNAblastDB.fa -out
"$transcriptID".blastn.miRNAseqDB.sam -num_threads 32 -outfmt '17 SQ'  -query
"$transcriptID".fa
sed -i "s/Query_1/$transcriptID/g" "$transcriptID".blastn.miRNAseqDB.sam
samtools view -bhS "$transcriptID".blastn.miRNAseqDB.sam >
"$transcriptID".blastn.miRNAseqDB.bam
samtools sort "$transcriptID".blastn.miRNAseqDB.bam -o
"$transcriptID".blastn.miRNAseqDB.sorted.bam
samtools index "$transcriptID".blastn.miRNAseqDB.sorted.bam

# Pilon correction
java -Xmx300g -jar
/media/morpheus/sagar/BUDDHA/Tasmanian_wolf/Chr_wise/Chromosomes/Final_verificatio
n/pilon-1.24.jar --genome "$transcriptID".fa --frags
"$transcriptID".blastn.DNAseqDB.sorted.bam --output "$transcriptID".pilon.DNAseq -
-fix all --changes

# create IGV report
create_report "$transcriptID".codon_positions.events.bed --standalone --fasta
"$transcriptID".fa --tracks "$transcriptID".blastn.DNAseqDB.sorted.bam
"$transcriptID".codon_positions.events.bed --output
"$transcriptID".BLASTn_IGV.html --info-columns Chromosome Start_position
End_position Event score strand thickStart thickEnd itemRgb --translate-sequence-
track

# Short-read verification using bam-readcount
awk '{print $1"\t"$2"\t"$3"\t"$4; if ($4 ~ /->/) {split($4, a, ":"); split(a[4],
change, "->"); start = $2 + 1; for (i = 1; i <= length(change[1]); i++) {ref_base
= substr(change[1], i, 1); alt_base = substr(change[2], i, 1); print
$1"\t"start"\t"ref_base"\t"alt_base; start++}}}'
"$transcriptID".codon_positions.events.bed | grep -v "STOP" | awk '$3!=$4' | awk
'{if ($4 ~ /Exon/) {split($4, a, ":"); print $1"\t"$2"-"$3"\t"a[3]"\t"a[4]} else
{if ($2 == $3) {print $1"\t"$2"-"$2"\t"$3"\t"$4} else {print $1"\t"$2"-
"$2"\t"$3"\t"$4}}}'|sed 's/\t/,/g' >
"$transcriptID".codon_positions.events.brc.csv
for i in `cat "$transcriptID".codon_positions.events.brc.csv`; do

  file_name=`echo $i | sed 's/,/_/g'`
  chr=`echo $i | cut -f1 -d','`
  start_end=`echo $i | cut -f2 -d','`
  mutation_type=`echo $i | cut -f3 -d','`   # Get the 3rd column (mutation type)
  ref_base=`echo $i | cut -f4 -d','`        # Get the 4th column (ref base)
  query_base=`echo $i | cut -f5 -d','`      # Get the 5th column (query base)

  if [[ "$mutation_type" != "FS_INS" && "$mutation_type" != "FS_DEL" ]]; then
    # Normal processing if not FS_INS or FS_DEL
    bam-readcount -w0 -f "$transcriptID".fa
"$transcriptID".blastn.DNAseqDB.sorted.bam $chr:$start_end >
"$transcriptID"."$chr":"$start_end".brc.tsv
    python
/media/morpheus/sagar/BUDDHA/Tasmanian_wolf/Chr_wise/Chromosomes/Final_verificatio
n/parse_brc.py "$transcriptID"."$chr":"$start_end".brc.tsv >
"$transcriptID"."$chr":"$start_end".parse_brc.tsv
```

```
    event_status=$(awk -v chr="$chr" -v ref_base="$ref_base" -v
query_base="$query_base" 'BEGIN {found=0} $7 >= 5 && !found {print "yes";
found=1}' "$transcriptID"."$chr":"$start_end".parse_brc.tsv)
    echo -e "$i,$event_status"

  else
    # Check for + sign in 4th column if FS_INS or - sign if FS_DEL
    bam-readcount -w0 -f "$transcriptID".fa
"$transcriptID".blastn.DNAseqDB.sorted.bam $chr:$start_end >
"$transcriptID"."$chr":"$start_end".brc.tsv
    python
/media/morpheus/sagar/BUDDHA/Tasmanian_wolf/Chr_wise/Chromosomes/Final_verificatio
n/parse_brc.py "$transcriptID"."$chr":"$start_end".brc.tsv >
"$transcriptID"."$chr":"$start_end".parse_brc.tsv

    if [[ "$mutation_type" == "FS_INS" ]]; then
      # Check for "+" in 4th column of parse_brc.tsv
      if cut -f4 "$transcriptID"."$chr":"$start_end".parse_brc.tsv | grep -q "^+"
; then
        echo -e "$i,yes"
      else
        echo -e "$i,No"
      fi

    elif [[ "$mutation_type" == "FS_DEL" ]]; then
      # Check for "-" in 4th column of parse_brc.tsv
      if cut -f4 "$transcriptID"."$chr":"$start_end".parse_brc.tsv | grep -q "^-";
then
        echo -e "$i,yes"
      else
        echo -e "$i,No"
      fi
    fi
  fi
done|sed 's/,/\t/g'> "$transcriptID".codon_positions.events.TOGA_confirm.tsv
sra_confirm=`awk 'BEGIN{total=0; yes_count=0} {total++; if($5 == "yes")
yes_count++} END {print yes_count "/" total, yes_count/total}'
"$transcriptID".codon_positions.events.TOGA_confirm.tsv|sed 's/ /\t/g'`
toga_status=`grep -w "$transcriptID"
/media/morpheus/sagar/BUDDHA/Tasmanian_wolf/Chr_wise/Chromosomes/Final_verificatio
n/TOGA.loss_summ_data.tsv|cut -f2`
echo
"TranscriptID,Id,Name,Description,Chromosome,ChrAccVer,ChrStart,ChrStop,ExonCount,
ORF status,GC Content,GC Stretch,Missing exon,Deleted exon,Splice site
change,START missing,TOGA status,Events supported in SRA,CL ratio"|sed 's/,/\t/g'
> "$transcriptID".info.tsv
echo -e
"$transcriptID\t$transcript_info_8c\t$orf_val\t$GC_Content\t$GC_Stretch\t$exon_mis
sing\t$exon_deleted\t$splice_site_change\t$start_missing\t$toga_status\t$sra_confi
rm" >> "$transcriptID".info.tsv

mkdir $transcriptID
mv "$transcriptID".*  $transcriptID
```

### e. HybPiper and Patchwork

We used HybPiper v2.3.1 [19] and Patchwork v0.5.2 [20] to recover gene sequence from short read data of thylacine. We also confirmed the presence of an in-frame STOP codon in the assembled sequence using HybPiper v2.3.1. The following script (with detailed parameters) was used to run Hybpiper v2.3.1 and Patchwork v0.5.2.

```
for i in *.fa
do
gene=`echo $i|sed 's/\.fa//g'`
#identify reads
blastn -task blastn  -evalue 0.05 -db
/media/morpheus/sagar/BUDDHA/Tasmanian_wolf/Hybpiper/SRA/SRR5055303-6.blastDB.fa -
out blastn."$gene".1out -num_threads 64 -outfmt 1 -query $i -max_target_seqs 10000
#get read ids list
grep "^SRR" blastn."$gene".1out|grep "^SRR5055303"|grep "/1"|cut -f1,2 -d' '  >
SRR5055303_1.lst
grep "^SRR" blastn."$gene".1out|grep "^SRR5055303"|grep "/2"|cut -f1,2 -d' '  >
SRR5055303_2.lst
grep "^SRR" blastn."$gene".1out|grep "^SRR5055304"|grep "/1"|cut -f1,2 -d' '  >
SRR5055304_1.lst
grep "^SRR" blastn."$gene".1out|grep "^SRR5055304"|grep "/2"|cut -f1,2 -d' '  >
SRR5055304_2.lst
grep "^SRR" blastn."$gene".1out|grep "^SRR5055305"|cut -f1 -d' '  > SRR5055305.lst
grep "^SRR" blastn."$gene".1out|grep "^SRR5055306"|cut -f1 -d' '  > SRR5055306.lst

#extract reads
seqtk subseq
/media/morpheus/sagar/BUDDHA/Tasmanian_wolf/Hybpiper/SRA/SRR5055303_1.fastq.gz
SRR5055303_1.lst > "$gene".SRR5055303_1.fq
seqtk subseq
/media/morpheus/sagar/BUDDHA/Tasmanian_wolf/Hybpiper/SRA/SRR5055303_2.fastq.gz
SRR5055303_2.lst > "$gene".SRR5055303_2.fq
seqtk subseq
/media/morpheus/sagar/BUDDHA/Tasmanian_wolf/Hybpiper/SRA/SRR5055304_1.fastq.gz
SRR5055304_1.lst > "$gene".SRR5055304_1.fq
seqtk subseq
/media/morpheus/sagar/BUDDHA/Tasmanian_wolf/Hybpiper/SRA/SRR5055304_2.fastq.gz
SRR5055304_2.lst > "$gene".SRR5055304_2.fq
seqtk subseq
/media/morpheus/sagar/BUDDHA/Tasmanian_wolf/Hybpiper/SRA/SRR5055305_1.fastq.gz
SRR5055305.lst > "$gene".SRR5055305_1.fq
seqtk subseq
/media/morpheus/sagar/BUDDHA/Tasmanian_wolf/Hybpiper/SRA/SRR5055305_2.fastq.gz
SRR5055305.lst > "$gene".SRR5055305_2.fq
seqtk subseq
/media/morpheus/sagar/BUDDHA/Tasmanian_wolf/Hybpiper/SRA/SRR5055306_1.fastq.gz
SRR5055306.lst > "$gene".SRR5055306_1.fq
seqtk subseq
/media/morpheus/sagar/BUDDHA/Tasmanian_wolf/Hybpiper/SRA/SRR5055306_2.fastq.gz
SRR5055306.lst > "$gene".SRR5055306_2.fq

#hybpiper

hybpiper assemble -r "$gene".SRR5055303_1.fq "$gene".SRR5055303_2.fq -t_dna $i --
prefix "$gene".SRR5055303_blast --evalue 0.05 --cpu 64
hybpiper assemble -r "$gene".SRR5055304_1.fq "$gene".SRR5055304_2.fq -t_dna $i --
prefix "$gene".SRR5055304_blast --evalue 0.05 --cpu 64
hybpiper assemble -r "$gene".SRR5055305_1.fq "$gene".SRR5055305_2.fq -t_dna $i --
prefix "$gene".SRR5055305_blast --evalue 0.05 --cpu 64
hybpiper assemble -r "$gene".SRR5055306_1.fq "$gene".SRR5055306_2.fq -t_dna $i --
prefix "$gene".SRR5055306_blast --evalue 0.05 --cpu 64

# retrive sequence

hybpiper retrieve_sequences  --targetfile_dna $i --single_sample_name
"$gene".SRR5055303_blast --fasta_dir SRR5055303.extracted_seq dna
hybpiper retrieve_sequences  --targetfile_dna $i --single_sample_name
"$gene".SRR5055304_blast --fasta_dir SRR5055304.extracted_seq dna
```

```
hybpiper retrieve_sequences  --targetfile_dna $i --single_sample_name
"$gene".SRR5055305_blast --fasta_dir SRR5055305.extracted_seq dna
hybpiper retrieve_sequences  --targetfile_dna $i --single_sample_name
"$gene".SRR5055306_blast --fasta_dir SRR5055306.extracted_seq dna

#patchwork

faTrans -stop $i "$gene".faTrans.fa
sed -i 's/>Sarcophilus_harrisii-/>Sarcophilus_harrisii@/g' "$gene".faTrans.fa
grep "^SRR" blastn."$gene".1out | cut -f1 -d' ' > "$gene".patchwork.lst
seqtk subseq /media/morpheus/sagar/BUDDHA/Tasmanian_wolf/Hybpiper/SRA/SRR5055303-
6.blastDB.fa "$gene".patchwork.lst > "$gene".patchwork.fasta
julia --
project=/media/morpheus/sagar/BUDDHA/Tasmanian_wolf/Hybpiper/tools/patchwork
/media/morpheus/sagar/BUDDHA/Tasmanian_wolf/Hybpiper/tools/patchwork/src/Patchwork
.jl --contigs "$gene".patchwork.fasta --reference "$gene".faTrans.fa --output-dir
"$gene"_patchwork

echo "HYBPIPER and PATCHWORK completed for $gene"
done
cd ..
```

Furthermore, NCBI RefSeq annotation is available for one more species from the *Dasyuridae* family, the yellow-footed antechinus (*Antechinus flavipes*). We cross-validated our gene loss results using the chromosome-level genome and available CDS annotations of *Antechinus flavipes* (yellow-footed antechinus). The antechinus genome reconfirmed the loss of the genes reported in our study. We also checked the gene order of the flaking gene in *Homo sapiens*, *Monodelphis domestica*, *Antechinus flavipes*, *Sarcophilus harrisii* and *Thylacinus cynocephalus* to confirm the 1-to-1 gene orthology.

### 3. Reconstruction of gene loss history

**a.** Retrieval of marsupial genomes: We retrieved 21 marsupial genomes from NCBI, including chromosomal-level and scaffold-level assemblies (see Supplementary Table S3). We used the NCBI datasets [21] command for this.

```
# 1_download_data.sh
curl -o datasets 'https://ftp.ncbi.nlm.nih.gov/pub/datasets/command-line/v2/linux-
amd64/datasets'
chmod 777 datasets
for i in `cat chr_marsupial.lst`
do
acc=`echo $i|cut -f1 -d'-'`
sp=`echo $i|cut -f2 -d'-'`
./datasets download genome accession $acc --filename "$sp"_dataset.zip --include
genome --api-key c181c44f2dca87803a9c0f9a24a32f67a608
unzip "$sp"_dataset.zip
genome=`ls -1 ncbi_dataset/data/$acc/*.fna |cut -f4 -d'/'`
mv ncbi_dataset/data/$acc/*.fna "$sp"-"$genome"
rm -r md5sum.txt "$sp"_dataset.zip ncbi_dataset README.md
done
```

**b. Making BLASTn database**

The following script is used to make a blast database.

```
# 2_make_database.sh
ls -1 *.fna > list_of_genomes
for genome in `cat list_of_genomes`
do
makeblastdb -in $genome -out $genome -dbtype nucl
faidx $genome -i chromsizes > "$genome".sizes.genome
done
```

c. **Extraction of syntenic gene locus sequence, genome alignment and inference of gene status**

Due to the substantial computational resources required for whole-genome alignment, we opted not to perform a full-genome comparison between the Tasmanian devil and the other 19 marsupial species. Instead, we carried out local genome alignments focused on each focal gene and its 30 kb flanking regions on both sides, ensuring that the syntenic context was retained. To identify orthologous genomic coordinates in each focal species, we conducted a BLASTn (with the parameters -evalue 0.05 -outfmt 7) search against each focal species genome assembly using the Tasmanian devil gene loss candidate gene sequence as a query. The resulting exonic hits were merged using BEDTools v2.27.1 [22] with the merge function with -d 10000. To ensure inclusion of syntenic context, we expanded the merged regions by 30 kb on both upstream and downstream ends using the slop function. Finally, we extracted the corresponding nucleotide sequences from the focal species' genome using the getfasta function. The duplicated sequences were removed with *seqkit rmdup*.

This local alignment strategy provided a computationally efficient yet synteny-aware framework for comparing gene intact/loss across marsupial genomes.

The region-specific sequences extracted for each focal gene (including ±30 kb flanking regions) were used to generate genome alignments in chain format compatible with TOGA. For this, we employed the make_lastz_chains pipeline (v2.0.8) available at https://github.com/hillerlab/make_lastz_chains, which utilises LASTZ v1.04.15 to perform pairwise alignments and processes the results using axtChain, chainCleaner, and RepeatFiller. These alignment chains and the Tasmanian devil genome annotation (used as the reference) were input into TOGA v1.1.7 to infer gene intact/loss across marsupial species. We validated the gene loss event for thylacine as discussed in the previous section. To validate TOGA-predicted gene loss events in other marsupials, we conducted a short-read search using publicly available whole-genome sequencing (WGS) data. Additionally, a species tree for all 21 marsupial species was obtained from the TimeTree database (https://timetree.org) and was used to place gene loss events in a phylogenetic context. See the detailed setting and script to perform the above task given below.

```
#genome_chain_toga.sh
genome=$1
i=$2
chr_id=$3
species_name=`echo $genome|cut -f1 -d'-'`
blastn -task blastn -evalue 0.05 -db $genome -out
$i/blastn_"$i"_cds_"$species_name".bls -num_threads 8 -outfmt 7 -query $i/"$i".fa
/media/morpheus/sagar/BUDDHA/Tools/blast2bed/blast2bed
$i/blastn_"$i"_cds_"$species_name".bls
sort-bed  $i/blastn_"$i"_cds_"$species_name".bed >
$i/blastn_"$i"_cds_"$species_name".sort.bed
mv $i/blastn_"$i"_cds_"$species_name".sort.bed
$i/blastn_"$i"_cds_"$species_name".bed
sed -i "/#.*/d;s/$i/$species_name/g" $i/blastn_"$i"_cds_"$species_name".bed
bedtools merge -s -header -d 10000 -i  $i/blastn_"$i"_cds_"$species_name".bed >
$i/blastn_"$i"_cds_"$species_name".merge.bed
#faidx $genome -i chromsizes > "$genome".sizes.genome
bedtools slop -i $i/blastn_"$i"_cds_"$species_name".merge.bed -g
"$genome".sizes.genome -b 30000 -header >
$i/blastn_"$i"_cds_"$species_name".30kbslop.bed
bedtools getfasta -fi $genome -bed $i/blastn_"$i"_cds_"$species_name".30kbslop.bed
> $i/"$species_name".30kbslop.fasta
```

```
seqkit rmdup -s  $i/"$species_name".30kbslop.fasta >
$i/"$species_name".30kbslop.fas
mv $i/"$species_name".30kbslop.fas $i/"$species_name".30kbslop.fasta
echo $species_name $genome

cd $i

sed -i 's/ .*//;s/\./_/;s/:/_/g;s/-//g;s/[()]//g' "$species_name".30kbslop.fasta
/media/morpheus/sagar/BUDDHA/TOGA_new/make_lastz_chains/make_chains.py
Sarcophilus_harrisii $species_name Sarcophilus_harrisii-"$chr_id".fa
"$species_name".30kbslop.fasta --chaining_memory 40 --project_dir
chain_Sarcophilus_harrisii_"$species_name" --kt
cd chain_Sarcophilus_harrisii_"$species_name"
mv query.2bit ../"$species_name".2bit
mv target.2bit ../Sarcophilus_harrisii.2bit
mv Sarcophilus_harrisii."$species_name".final.chain.gz ../
cd ..
rm -r chain_Sarcophilus_harrisii_"$species_name"

for chain in Sarcophilus_harrisii."$species_name".final.chain.gz
do
bed_input="$i"-GCF_902635505.1_mSarHar1.11_genomic.bed12
id=`cat $bed_input|cut -f4`
/media/morpheus/sagar/BUDDHA/TOGA/toga.py $chain $bed_input
Sarcophilus_harrisii.2bit "$species_name".2bit --pn
TOGA_Sarcophilus_harrisii_"$species_name" --nc
/media/morpheus/sagar/BUDDHA/TOGA/nextflow_config_files/ --cesar_bigmem_config
/media/morpheus/sagar/BUDDHA/TOGA/nextflow_config_files/cesar_bigmem_config.nf --
cesar_jobs_num 500 --cesar_buckets 3,5,25,50 --ces --kt --chain_jobs_num 60
/media/morpheus/sagar/BUDDHA/TOGA/supply/plot_mutations.py --publication_mode_heni
$bed_input TOGA_Sarcophilus_harrisii_"$species_name"/inact_mut_data.txt $id
sorted/mutation_plot/"$species_name".svg
done

for toga in TOGA_Sarcophilus_harrisii_"$species_name"
do
cat $toga/inact_mut_data.txt| sed "s/$id/$species_name/g" >>
sorted/"$i".inact_mut_data.txt
orthology_classification=`cat $toga/orthology_classification.tsv|tail -1|cut -f5`;
loss_summ=`cat $toga/loss_summ_data.tsv|tail -1|cut -f3`
echo -e "$species_name\t$orthology_classification\t$loss_summ" >>
sorted/"$i".orthology.loss_summ.tsv
cat $toga/codon.fasta |tail -2 |sed 's/ //g;s/X/N/g'|sed "s/>.*/>$species_name/g"
>> sorted/"$i".codon.fasta
cat $toga/nucleotide.fasta |tail -2 |sed 's/ //g'|sed "s/>.*/>$species_name/g" >>
sorted/"$i".nucleotide.fasta
done
cd ..
```

```
#3_21_species_parallel_extract_seq_chain_toga.sh
#!/bin/bash

##create transcript.lst file with all gene transcripts needed to be checked
# Initialize variables and environment
i=$1
ulimit -n 16384
export JAVA_HOME=/usr/lib/jvm/java-11-openjdk-amd64
export PATH=$JAVA_HOME/bin:$PATH
export NXF_VER=22.10.0

# Ensure directories exist and retrieve necessary data
```

```
[ ! -d "$i" ] && mkdir -p  $i/sorted/mutation_plot
efetch -db nuccore -id $i -format fasta_cds_na | cut -f1,2 -d'_' | sed
's/>lcl|/>/g' > $i/"$i".fa
length=$(faidx  $i/"$i".fa -i chromsizes | cut -f2)
GC=$(seqkit fx2tab --name --gc  $i/"$i".fa | cut -f2)
echo -e "$i\t$length\t$GC" > $i/"$i".tsv
chr_id=$(grep -w "$i" GCF_902635505.1_mSarHar1.11_genomic.bed12 | cut -f1 | sed
's/_/\./2')
grep -w "$i" GCF_902635505.1_mSarHar1.11_genomic.bed12 > $i/"$i"-
GCF_902635505.1_mSarHar1.11_genomic.bed12
echo $chr_id > $i/chr.lst
seqtk subseq Sarcophilus_harrisii-GCF_902635505.1_mSarHar1.11_genomic.fna
$i/chr.lst -l 60 > $i/Sarcophilus_harrisii-"$chr_id".fa
sed -i 's/ .*//;s/\./_/' $i/Sarcophilus_harrisii-"$chr_id".fa

# Run parallel jobs for multiple species
./genome_chain_toga.sh Antechinus_flavipes-GCA_016432865.2_AdamAnt_v2_genomic.fna
$i $chr_id &
./genome_chain_toga.sh Dasyurus_viverrinus-
GCA_020854095.1_UniMelb_DasViv_v1.0_genomic.fna $i $chr_id &
./genome_chain_toga.sh Dromiciops_gliroides-
GCA_019393635.1_mDroGli1.pri_genomic.fna $i $chr_id &
./genome_chain_toga.sh Gracilinanus_agilis-GCA_016433145.1_AgileGrace_genomic.fna
$i $chr_id &
./genome_chain_toga.sh Lagorchestes_hirsutus-
GCA_028533205.1_Lagorchestes_hirsutus_HiC_genomic.fna $i $chr_id &
./genome_chain_toga.sh Macropus_fuliginosus-GCA_028583105.1_mf-2k_genomic.fna $i
$chr_id &
./genome_chain_toga.sh Macropus_giganteus-GCA_028627215.1_mg-2k_genomic.fna $i
$chr_id &
./genome_chain_toga.sh Macrotis_lagotis-
GCA_037893015.1_bilby.v1.9.chrom.fasta_genomic.fna $i $chr_id &
./genome_chain_toga.sh Monodelphis_domestica-
GCA_027887165.1_mMonDom1.pri_genomic.fna $i $chr_id &
wait
./genome_chain_toga.sh Notamacropus_eugenii-
GCA_028372415.1_mMacEug1.pri_genomic.fna $i $chr_id &
./genome_chain_toga.sh Phalanger_gymnotis-GCA_028646595.1_pg-2k_genomic.fna $i
$chr_id &
./genome_chain_toga.sh Potorous_gilbertii-
GCA_028658325.1_Potorous_gilbertii_HiC_genomic.fna $i $chr_id &
./genome_chain_toga.sh Pseudocheirus_occidentalis-
GCA_028646575.1_Pseudocheirus_occidentalis_HiC_genomic.fna $i $chr_id &
./genome_chain_toga.sh Pseudochirops_corinnae-
GCA_028646515.1_Pseudochirops_corinnae_HiC_genomic.fna $i $chr_id &
./genome_chain_toga.sh Pseudochirops_cupreus-
GCA_028627135.1_Pseudochirops_cupreus_HiC_genomic.fna $i $chr_id &
./genome_chain_toga.sh Thylacinus_cynocephalus-
GCA_007646695.3_UniMelb_ThyCyn2.0_hybrid_assembly_genomic.fna $i $chr_id &
./genome_chain_toga.sh Trichosurus_vulpecula-
GCA_011100635.1_mTriVul1.pri_genomic.fna $i $chr_id &
./genome_chain_toga.sh Vombatus_ursinus-GCA_028626985.1_vu-2k_genomic.fna $i
$chr_id &
./genome_chain_toga.sh Myrmecobius_fasciatus-
GCA_023553655.1_mMyrfas1.20211206_genomic.fna $i $chr_id &
./genome_chain_toga.sh Sminthopsis_crassicaudata-Dunnart_asm_12-2021_sm.fa $i
$chr_id &
# Wait for all parallel jobs to finish
wait

# Continue with the next steps
echo "All jobs for $i completed."
```

Furthermore, we assessed the integrity of the genome assembly of grey short-tailed opossum (*Monodelphis domestica*) using the klumpy tool v1.0.11 [23]. For this, input the BAM file created using BWA-MEM v0.7.17 [10] with -x pacbio flag. This ensured that the absence of the gene was not due to assembly gaps or annotation errors (see electronic supplementary material, figure S15). We used the following code to confirm aseelbly.

```
#mapping with bwa
bwa index GCF_027887165.1_mMonDom1.pri_genomic.fna
for i in m54306Ue_211111_163748.hifi_reads.fastq.gz
m54306Ue_211113_015853.hifi_reads.fastq.gz
m64055e_211117_204835.hifi_reads.fastq.gz
m64330e_211103_063050.hifi_reads.fastq.gz
m64334e_211117_205854.hifi_reads.fastq.gz
do
k=`echo $i|cut -f1 -d'.'`
bwa mem -t 64 -x pacbio GCF_027887165.1_mMonDom1.pri_genomic.fna $i |samtools sort
-@ 64 -O BAM -o "$k".bwa.pacbio.bam
samtools index -c "$k".bwa.pacbio.bam
done
samtools merge -@ 32 MonDom1.merged.bam *.bwa.pacbio.bam
rm *.bwa.pacbio.bam
samtools sort MonDom1.merged.bam  -@ 48 -o Monodelphis_domestica.merge.sorted.bam
samtools index -c -@ 48 Monodelphis_domestica.merge.sorted.bam
#Klumpy
klumpy scan_alignments --alignment_map Monodelphis_domestica.merge.sorted.bam --
threads 16
klumpy alignment_plot --alignment_map Monodelphis_domestica.merge.sorted.bam --
reference NC_077232.1 --candidates
Monodelphis_domestica.merge.sorted_Candidate_Regions.tsv --min_len 3000 --
window_size 10000 --window_step 5000 --color red --vertical_line_gaps --
vertical_line_klumps --format svg --leftbound 266519535 --rightbound 266794692 --
annotation GCF_027887165.1_mMonDom1.pri_genomic.gtf --gap_file
Monodelphis_domestica.HSD17B13.NC_077232.1_gaps.tsv
#Found the following genes: IBSP NUDT9 LOC103092505 LOC100016979 KLHL8
```

### 4. Transcriptional status of lost genes

#### a. BLASTn search in miRNA-seq data

We screened the expression of the lost gene in the thylacine miRNA-seq short-read database (SRR23147611, SRR23147612, SRR23147613, SRR23147614, SRR23147615 and SRR23147616) [24] using BLASTn v2.13.0 [6,25]. To validate the presence of protein-coding gene expression in this dataset, we also queried (as a positive control) the sequences of LOC100913894 (*ACTA*) and LOC100925998 (*MYH7*), based on a previous report [24].

```
# BLASTn in RNAseqDB
blastn -task blastn -evalue 0.01 -max_target_seqs 5000 -db
/media/morpheus/sagar/BUDDHA/Tasmanian_wolf/Chr_wise/Chromosomes/mapping/miRNA_seq
/miRNAblastDB/SRR23147611-6.miRNAblastDB.fa -out
"$transcriptID".blastn.miRNAseqDB.sam -num_threads 32 -outfmt '17 SQ'  -query
"$transcriptID".fa
sed -i "s/Query_1/$transcriptID/g" "$transcriptID".blastn.miRNAseqDB.sam
samtools view -bhS "$transcriptID".blastn.miRNAseqDB.sam >
"$transcriptID".blastn.miRNAseqDB.bam
samtools sort "$transcriptID".blastn.miRNAseqDB.bam -o
"$transcriptID".blastn.miRNAseqDB.sorted.bam
samtools index "$transcriptID".blastn.miRNAseqDB.sorted.bam
```

#### b. Transcriptional status of lost genes in dunnart and Tasmanian devil

We used the RNA-Seq data from 27 Tasmanian devil and dunnart tissues (PRJEB34650 [26], PRJNA1028148 [27]; see Supplementary Table S4) for expression analysis. Using the following script, we obtained the TPM values via pseudo-alignment using Kallisto v0.51.0.

```
## download cds fasta
wget
https://ftp.ncbi.nlm.nih.gov/genomes/all/GCF/902/635/505/GCF_902635505.1_mSarHar1.11/GCF_902635505.1_mSarHar1.11_cds_from_genomic.fna.gz
gunzip GCF_902635505.1_mSarHar1.11_cds_from_genomic.fna.gz
#change the header of CDS file
sed 's/^.*gene=/>/' GCF_902635505.1_mSarHar1.11_cds_from_genomic.fna | cut -f1,4 -d']'|sed 's/\[protein_id=//g;s/\]//g;s/ /-/g' >
Reformated.GCF_902635505.1_mSarHar1.11_cds_from_genomic.fna
for i in `cat protein.lst`;do id=`esearch -db protein -query $i|  efetch -format gp|grep "DBSOURCE"|sed 's/DBSOURCE    REFSEQ: accession //g'`; sed -i "s/$i/$i-$id/g" Reformated.GCF_902635505.1_mSarHar1.11_cds_from_genomic.fna; echo $i-$id;done
#Kallisto
kallisto index --make-unique -i transcripts.idx
Reformated.GCF_902635505.1_mSarHar1.11_cds_from_genomic.fna
for i in `ls -1 *_1.fastq.gz`
do
j=`echo $i|sed 's/_1/_2/g'`
k=`echo $i|sed 's/_1.fastq.gz//g'`
kallisto quant -t 32 -i transcripts.idx -o "$k".kallisto_out -b 100 $i $j
done
```

c. **Assessment of gene loss and relaxed selection signatures at the transcriptional level**

We confirmed the *VWA7* gene lacks expression in numbats' tongue, lung, and liver tissues [28]. We used STAR-read aligner [29] to map RNA-seq reads to the genome (see electronic supplementary material, figure S18).

```
#map_RNA.sh
STAR --runThreadN 16 --runMode genomeGenerate --genomeDir . --genomeFastaFiles
Myrmecobius_fasciatus.VWA7.JAJPUD010000745.1.fa --genomeSAindexNbases 8
## for mapping
for i in *_1.fastq.gz
do
j=`echo $i|sed 's/_1/_2/g'`
k=`echo $i|sed 's/_1.fastq.gz//g'`
STAR --runThreadN 16 --outSAMtype BAM SortedByCoordinate --genomeDir . --readFilesIn $i $j --readFilesCommand zcat --outFileNamePrefix "$k"_ --limitBAMsortRAM 1031735638
samtools index "$k"_*.bam
done
```

Similarly, *SAMD9* and *SAMD9L* expression status in *Macropus eugenii* assed by mapping RNA seq read to its genome (GCF_028372415.1) and visusualised with IGV v1.12.0 [9] (see electronic supplementary material, figure S12).

### 5. Evaluating the strength of selection

#### a. Dataset preparation

To test the relaxation of selection in gene loss candidates, we obtained pairwise codon alignment sequences between the Tasmanian devil and other marsupials, including the thylacine from TOGA v1.1.7 output. Codons affected by frameshifting insertions or deletions and premature stop codons were replaced by "NNN" to maintain a reading frame. A multiple sequence alignment (MSA) was generated using PRANK v.170427 [30] in GUIDANCE2 v2.01 [31]. The time-calibrated phylogenetic tree was obtained from the Timetree website (www.timetree.org).

```
clade=$1
cd $clade
# to remove internode labels from the TimeTree nwk file
for i in "$clade".nwk
do
sed -e "s/'[^()]*'//g" $i > temp.nwk
mv temp.nwk $i
echo 'library(ape)' > tree_script.r
echo "a<-read.tree(\"$i\")" >> tree_script.r
echo 'b<-unroot(a)' >> tree_script.r
echo "write.tree(b,\"$i.tree\")" >> tree_script.r
Rscript tree_script.r
mv $i.tree $i
done

# This script checks whether the species names in the fasta and nwk files are
identical.
for i in "$clade".fa
do
grep ">" $i|sed 's/>//g' > $i.txt
j="$clade".nwk
sed 's/(/\n/g' $j|sed 's/)/\n/g' |sed 's/;/\n/g' |sed 's/:/\n/g' |sed 's/,/\n/g'
|grep "^[A-Z]" > $j.txt
echo $j
cat $i.txt $j.txt |sort|uniq -c |awk '$1<2 {print $2}'
rm *.txt
done
# MSA using PRANK in Guidance2
guidance=/home/ceglab358/BUDDHA/Tools/guidance.v2.02/www/Guidance/guidance.pl
for i in `ls "$clade".fa`
do
j=`echo $i|sed 's/.fa//g'`
perl $guidance --program GUIDANCE --seqFile "$i" --msaProgram PRANK --seqType
codon --outDir "$i".100_PRANK --genCode 1 --bootstraps 100 --proc_num 16
cp "$i".100_PRANK/MSA.PRANK.aln.With_Names "$j".aln
rm -r "$i".100_PRANK
done
cd ..
```

**b. Selection analysis using HYPHY**

We used HyPhy's [32] RELAX [33], aBSREL [34], BUSTED [35], FEL [36], and MEME [37] models to detect relaxed and positive selection. Multiple testing was adjusted using the FDR method. Relaxation or positive selection test is done for each species by putting it as the foreground species in each model of the HYPHY program.

```
#HYPHY RELAX
hyphy relax --alignment $i --tree "$j"_treeLabled.txt --test fg --output
"$j"_treeoutput_relax
#HYPHY aBSREL
HYPHYMP aBSREL --alignment $i --tree  "$j"_treeLabled.txt --branches fg >
"$j"_treeoutput_aBSREL
#HYPHYMP BUSTED
HYPHYMP busted --alignment $i --tree  "$j"_treeLabled.txt --branches fg  >
"$j"_treeoutput_BUSTED
#HYPHY MEME
HYPHYMP meme --alignment $i --tree  "$j"_treeLabled.txt --branches fg
>"$j"_treeoutput_meme
#HYPHY FEL
```

```
HYPHYMP fel --alignment $i --tree  "$j"_treeLabled.txt --branches fg
>"$j"_treeoutput_fel
```

**c. Selection analysis using codeML**

To estimate signatures of selection across branches, we used the codeml program from the PAML package [38]. Analyses were conducted using unrooted gene trees, and we estimated branch-specific dN/dS (ω) ratios under multiple models. Specifically, we employed the M0, bfree, and bneutral branch models to test for variable selection pressures across branches. Codon frequencies were modelled using both the F3×4 and F1×4 schemes to account for differences in nucleotide composition across codon positions. For more details, please see Shinde, Sagar Sharad et al. [39].

**d. GC-biased gene conversion (gBGC)**

We estimated the magnitude of gBGC using MapNH v1.3.0 [40] and phastBias v1.6 [41]. For more details, please see Shinde, Sagar Sharad et al. [35].

```
# phastBias
phastBias --bgc 3 --output-tracts $i.gff $i $i.mod $k > $k.wig
# mapnh
mapnh map.type=GC output.counts.tree.prefix=all_gc.prefix2 input.sequence.file=$i
input.tree.file=$tree test.branch=1 test.global=1 model=K80
```

### 6. Limitations with GC-content and paralogues

One of the major findings from our study is the need for several precautions while identifying gene loss events in genome assemblies currently available for most vertebrates. Claims of gene loss that fail to consider these factors are prone to high false positive rates [42–45]. The effect of GC content on the rapid evolutionary divergence of genes and the challenge of picking up such genes using short-read sequencing approaches is demonstrated in rodents and birds [46–48]. A recent review highlights the evolutionary significance of GC content and intragenomic mutational heterogeneity [49].

To ensure that we have explored these aspects in greater depth, we have shown the statistical significance of the effect of GC content and sequence identity between paralogs. The pairwise Wilcoxson tests demonstrate that genes classified as intact by TOGA have a significantly lower GC content than those identified as lost, missing, partially intact, partially missing and unclear loss. The missing genes have a mean GC content of 60.35%; at least some of these are likely to be recovered using long-read sequencing. We avoid false positive claims of gene loss by excluding these missing genes from our analysis. The motivation for keeping GC content vs the average length of GC stretches while highlighting the gene loss we have demonstrated is to clarify that these events are not due to high GC. In addition, this explains the TOGA classification of missing genes in the thylacine genome assembly based solely on short-read data. (figure 3a, see electronic supplementary material, figure S2).

We used a similar approach to evaluate the effect of paralog percent identity on the TOGA classification of gene status. However, we did not find any clear pattern of percent identity of the paralog determining the gene status (see electronic supplementary material, figure S2). We also compared GC content vs percent identity of the paralog along with the evolutionary age of the paralog (i.e., paralog last common ancestor with Tasmanian Devil). We found that genes classified as intact or partially intact by TOGA tend to have genes with high sequence identity with the paralog. However, genes classified as lost, missing and unclear loss have a paucity of genes with high sequence identity with the paralog (see electronic supplementary material, figure S3). The observed pattern suggests that paralog identity has less effect on TOGA classification than GC content. Only some copies of the paralogs may be assembled in the genome, leading to this observation. Hence, some gene loss events identified using the genome assembly may be false positives.

The above analysis does not provide specific examples of how the paralogs affect identifying individual gene loss events. To overcome this, we performed another analysis comparing the GC content of the focal gene with the GC content of its paralogs and the percent identity with the paralog. In this analysis, we ranked the genes (x-axis) and their corresponding paralogs (y-axis) by GC content (see electronic supplementary material, figure S4). Although TOGA classifies genes as UL (Unclear Loss), PI (Partially Intact) due to limitations in data, not all genes that are classified as clearly Lost (L) are true gene loss events. For instance, incorrectly assembling closely related paralogs can lead to anomalous results. For example, the *PAX7* gene and its paralog affect the validation of the frame-disrupting mutation (see electronic supplementary material, figure S5).

**a. Data retrieval from NCBI, calculation of GC-content and GC stretches**

We used the Tasmanian devil coding DNA sequences to calculate GC-content and GC-streaches with the following script.

```
#!/bin/bash
# Create header for the output TSV file
echo -e "Gene_name\tTranscriptID\tChromosome\tTranscript_length\tGC
Content\tGC_Stretch\tMissing_exon\tDeleted_exon\tTOGA_status" >
S.harrisii.info.tsv

# Iterate over filtered transcript IDs
for i in $(grep -vE "unassigned_transcript|XR|NR" loss_summ_data.reformated.tsv |
cut -f1); do
    TranscriptID="$i"

    # Get the gene name
    Gene_name=$(awk -v var1="$TranscriptID" '$3 == "transcript" && $0 ~
"transcript_id \"" var1 "\"" { match($0, /gene_id "([^"]+)"/, arr); print arr[1]
}' GCF_902635505.1_mSarHar1.11_genomic.gtf)

    # Get chromosome information
    Chromosome=$(grep "$TranscriptID" GCF_902635505.1_mSarHar1.11_genomic.gtf |
awk '$3 == "transcript"' | cut -f1)

    # Extract the transcript sequence
    grep -A1 ">ref:$TranscriptID" nucleotide.reformated.fasta | sed 's/:/_/g' >
"$TranscriptID.fa"

    # Get the length and GC content
    length=$(faidx "$TranscriptID.fa" -i chromsizes | cut -f2)
    GC=$(seqkit fx2tab --name --gc "$TranscriptID.fa" | cut -f2)

    # Calculate GC stretch
    GC_Stretch=$(perl
/media/morpheus/sagar/BUDDHA/Tasmanian_wolf/Chr_wise/Chromosomes/Final_verificatio
n/GC_Stretch_finder.pl "$TranscriptID.fa" | awk '{print $5}')

    # Get inactive mutation data
    grep "$TranscriptID" inact_mut_data.txt > "$TranscriptID.inact_mut_data.txt"

    # Check for missing and deleted exons
    exon_missing="No"
    if grep -q "Missing exon" "$TranscriptID.inact_mut_data.txt"; then
        exon_missing="Yes"
    fi
```

```
    exon_deleted="No"
    if grep -q "Deleted exon" "$TranscriptID.inact_mut_data.txt"; then
        exon_deleted="Yes"
    fi

    # Get TOGA status
    toga_status=$(grep -w "$TranscriptID" loss_summ_data.reformated.tsv | cut -f2)

    # Append results to the output file
    echo -e
"$Gene_name\t$TranscriptID\t$Chromosome\t$length\t$GC\t$GC_Stretch\t$exon_missing\
t$exon_deleted\t$toga_status" >> S.harrisii.info.tsv

    # Print TranscriptID for tracking
    echo "$TranscriptID"

    # Clean up temporary files
    rm "$TranscriptID.fa" "$TranscriptID.inact_mut_data.txt"
"$TranscriptID.fa.fai"
done

#awk 'BEGIN{FS="\t"} NF != 9 {print "Row", NR, "has", NF, "columns."}'
S.harrisii.info.tsv
## Gel longest isoform
head -n1 S.harrisii.info.tsv > S.harrisii.longest_isoform.info.tsv
tail -n +2 S.harrisii.info.tsv | \
cut -f1 | \
sort -u | \
while read GENE; do
    LONGEST=$(grep -w "^$GENE" S.harrisii.info.tsv | \
              sort -t$'\t' -k4,4nr | \
              head -n1)
    printf "$LONGEST\n" >> longest.list
done
cat longest.list | while read LONGEST_ISOFOMR; do
    grep -F "$LONGEST_ISOFOMR" S.harrisii.info.tsv >> tmp.tsv
done
sort -k1,1 tmp.tsv >> S.harrisii.longest_isoform.info.tsv
rm longest.list tmp.tsv
```

**b. Retrieval of paralog information from ENSEMBL**

We used ENSEMBL [50] biomart to retrieve paralog information for Tasmanian devils genes (https://training.ensembl.org/exercise/BioMart_paralogues). We added the information of gene status in thylacine (inferred using TOGA) to this info.

```
# Tasmanian_devil_paralogue_mart_export.txt obtained from ENSEMBL BIOMART
head -1 Tasmanian_devil_paralogue_mart_export.txt | awk '{print $0
"\tTOGA_status"}' > TOGA_info.nonempty.Tasmanian_devil_paralogue_mart_export.tsv

# Loop to annotate and filter
for i in $(cut -f4 Tasmanian_devil_paralogue_mart_export.txt | sort -u); do
    TOGA_status=$(awk -v gene="$i" '$1 == gene {print $9; exit}'
S.harrisii.info.plot.tsv)
    if [[ -n "$TOGA_status" ]]; then
        awk -v gene="$i" -v status="$TOGA_status" '$4 == gene {print $0 "\t"
status}' Tasmanian_devil_paralogue_mart_export.txt
    fi
done >> TOGA_info.nonempty.Tasmanian_devil_paralogue_mart_export.tsv
```

**c. Plotting**

The following code is used to visualise:

**i. Effect of paralog sequence identity on TOGA classification**

```
########## rain cloud plot ########

library(devtools)
library(ggside)
library(tidyverse)
library(tidyquant)
library(dplyr)
library(ggplot2)
library(ggrepel)
library(gghalves)
data <-
read.table("./TOGA_info.nonempty.Tasmanian_devil_paralogue_mart_export.tsv",
header = T, sep = "\t")
data<-na.omit(data)
data[1,]
mean_df <- data %>%
  group_by(TOGA_status) %>%
  summarise(mean_iden =
mean(`Paralogue_.id._target_Tasmanian_devil_gene_identical_to_query_gene`, na.rm =
TRUE))
pairwise.wilcox.test(data$Paralogue_.id._target_Tasmanian_devil_gene_identical_to_
query_gene, data$TOGA_status, p.adjust.method = "BH")
count_df <- data %>%
  group_by(TOGA_status) %>%
  summarise(n = n(),
            mean_iden =
mean(`Paralogue_.id._target_Tasmanian_devil_gene_identical_to_query_gene`, na.rm =
TRUE))

# Define pairwise combinations of interest
comparisons <- list(
  c("L", "I"),
  c("M", "I"),
  c("PI", "I"),
  c("PM", "I"),
  c("UL", "I"),
  c("M", "L"),
  c("PI", "L"),
  c("PM", "L"),
  c("UL", "L"),
  c("PI", "M"),
  c("PM", "M"),
  c("UL", "M"),
  c("PM", "PI"),
  c("UL", "PI"),
  c("UL", "PM")
)



# Basic raincloud plot
paralog_iden<-ggplot(data, aes(x = TOGA_status, y =
Paralogue_.id._target_Tasmanian_devil_gene_identical_to_query_gene, fill =
TOGA_status)) +

  # Half-violin plot (distribution)
  geom_half_violin(
    aes(fill = TOGA_status),
    side = "l",
    alpha = 0.6,
```

```
    color = NA,
    trim = FALSE
  ) +
  geom_text(data = count_df, aes(x = TOGA_status, y = 95, label = paste0("n=",
n)),
            size = 3.5, color = "black")+

  # Boxplot in the middle
  geom_boxplot(
    width = 0.15,
    outlier.shape = NA,
    alpha = 0.7
  ) +

  # Jittered individual data points
  geom_half_point(
    side = "r",
    shape = 21,
    size = 1,
    alpha = 0.4,
    position = position_jitter(width = 0.07)
  ) +
  stat_compare_means(method = "wilcox.test", label = "p.signif", comparisons =
comparisons) +
  stat_summary(fun = mean, geom = "point", shape = 23, size = 3,
               fill = "white", color = "black", stroke = 1.2) +
  geom_text(data = mean_df, aes(x = TOGA_status, y = mean_iden,
                                label = sprintf("%.2f", mean_iden)),
            vjust = 1,hjust =1.5,  size = 4, color = "black") +
  labs(x = "TOGA Status",
       y = "% Identity (Target Tasmanian devil gene vs Paralog)"
  ) +
  theme_minimal() +
  theme(legend.position = "none")


data <- read.table("S.harrisii.info.plot.tsv", header = T, sep = "\t")
count_df <- data %>%
  group_by(TOGA_status) %>%
  summarise(n = n(),
            mean_iden = mean(GC_Content, na.rm = TRUE))
mean_df <- data %>%
  group_by(TOGA_status) %>%
  summarise(mean_gc = mean(GC_Content, na.rm = TRUE))
pairwise.wilcox.test(data$GC_Content, data$TOGA_status, p.adjust.method = "BH")
# Define pairwise combinations of interest
comparisons <- list(
  c("L", "I"),
  c("M", "I"),
  c("PI", "I"),
  c("PM", "I"),
  c("UL", "I"),
  c("M", "L"),
  c("PI", "L"),
  c("PM", "L"),
  c("UL", "L"),
  c("PI", "M"),
  c("PM", "M"),
  c("UL", "M"),
  c("PM", "PI"),
  c("UL", "PI"),
  c("UL", "PM")
)
```

```
# Basic raincloud plot
GC_content<-ggplot(data, aes(x = TOGA_status, y = GC_Content, fill = TOGA_status))
+

  # Half-violin plot (distribution)
  geom_half_violin(
    aes(fill = TOGA_status),
    side = "l",
    alpha = 0.6,
    color = NA,
    trim = FALSE
  ) +
  geom_text(data = count_df, aes(x = TOGA_status, y = 95, label = paste0("n=",
n)),
            size = 3.5, color = "black")+
  # Boxplot in the middle
  geom_boxplot(
    width = 0.15,
    outlier.shape = NA,
    alpha = 0.7
  ) +

  # Jittered individual data points
  geom_half_point(
    side = "r",
    shape = 21,
    size = 1,
    alpha = 0.4,
    position = position_jitter(width = 0.07)
  ) +
  stat_compare_means(method = "wilcox.test", label = "p.signif", comparisons =
comparisons) +
  stat_summary(fun = mean, geom = "point", shape = 23, size = 3,
               fill = "white", color = "black", stroke = 1.2) +
  geom_text(data = mean_df, aes(x = TOGA_status, y = mean_gc,
                                label = sprintf("%.2f", mean_gc)),
            vjust = 1,hjust =1.5,  size = 4, color = "black") +
  labs(x = "TOGA Status",
       y = "GC Content (%)"
  ) +
  theme_minimal() +
  theme(legend.position = "none")


p1 <- GC_content +
  rremove("x.text") +
  rremove("xlab") +
  theme_classic() +
  theme(
    legend.position = "bottom",
    legend.box = "horizontal" )+
      guides(fill = guide_legend(nrow = 1))

p2 <- paralog_iden +
  theme_classic() +
  theme(
    legend.position = "bottom",
    legend.box = "horizontal"
  ) +
  guides(fill = guide_legend(nrow = 1))
```

```
# Arrange with common one-row legend
png("GC_content_Vs_per.identity.raincloud_plot_with_pairwise_wilcox.png",
units="in", width=16, height=9, res=900)
ggarrange(
  p1,
  p2,
  align = "v",
  ncol = 1,
  nrow = 2,
  common.legend = TRUE,
  legend = "bottom"
)
dev.off()
```

#### ii. Effect of GC-content and sequence identity between Tasmanian devil gene and paralog (%)

```
library(tidyverse)
library(dplyr)
library(ggplot2)

df <- read.table("TOGA_info.nonempty.Tasmanian_devil_paralogue_mart_export.tsv",
header = TRUE, sep = '\t')
df<-na.omit(df)


facet_GC<-ggplot(df, aes(
  x = Paralogue_.id._target_Tasmanian_devil_gene_identical_to_query_gene,
  y = Gene_._GC_content,
  color = Paralogue_last_common_ancestor_with_Tasmanian_devil
)) +
  geom_point(alpha = 0.6, size = 1.5) +
  facet_wrap(~ TOGA_status) +
  theme_classic2() +
  labs(
    x = "% Identity (Target Tasmanian devil gene vs Paralog)",
    y = "GC Content (%)",
  )+
  theme(legend.position = "bottom")

png("GC_content_Vs_per.identity.png", units="in", width=16, height=9, res=900)
facet_GC
dev.off()
```

#### iii.Effect of GC-content, sequence identity between paralog on TOGA classification (genes ranked by GC-content)

```
############# Gene rank plot #################
getwd()
library(ggplot2)
library(dplyr)
library(tidyverse)
library(forcats)
library(ggpubr)


data <-
read.table("./TOGA_info.nonempty.Tasmanian_devil_paralogue_mart_export.tsv",
header = T, sep = "\t")
data<-na.omit(data)
df<- data[,c(1,4,6,8,10,15,16,17,19)]
```

```
head(df)
df_unique <- df[!duplicated(df), ]
head(df_unique)
###############
gene_gc_ranked <- df_unique %>%
  select(Gene_stable_ID, gene_gc = Gene_._GC_content) %>%
  distinct() %>%
  arrange(gene_gc) %>%
  mutate(Gene_stable_ID_rank = row_number())
length(unique(gene_gc_ranked$Gene_stable_ID_rank))
head(gene_gc_ranked)
# Step 2: Rank unique paralogue GC content
paralog_gc_ranked <- df_unique %>%
  select(Tasmanian_devil_paralogue_gene_stable_ID, gene_gc = Gene_._GC_content)
%>%
  distinct() %>%
  arrange(gene_gc) %>%
  mutate(Tasmanian_devil_paralogue_gene_stable_ID_rank = row_number())
length(unique(paralog_gc_ranked$Tasmanian_devil_paralogue_gene_stable_ID_rank))
head(paralog_gc_ranked)



df_plot <- df_unique %>%
  left_join(gene_gc_ranked, by = "Gene_stable_ID") %>%
  left_join(paralog_gc_ranked, by = "Tasmanian_devil_paralogue_gene_stable_ID")

head(df_plot)
View(df_plot)
###############
df_plot <- df_plot %>%
  mutate(
    identity_bin_raw = cut(
      Paralogue_.id._target_Tasmanian_devil_gene_identical_to_query_gene,
      breaks = c(0, 25, 50, 75, 100),
      include.lowest = TRUE,
      right = FALSE  # [0-25), [25-50), etc.
    )
  )

# Step 2: Create readable labels
df_plot <- df_plot %>%
  mutate(
    identity_bin_labeled = recode_factor(identity_bin_raw,
                                         "[0,25)" = "0-25%",
                                         "[25,50)" = "25-50%",
                                         "[50,75)" = "50-75%",
                                         "[75,100]" = "75-100%"
    )
  )

# Step 3: Reorder levels (for legend ordering)
df_plot <- df_plot %>%
  mutate(
    identity_bin = fct_relevel(identity_bin_labeled, "0-25%", "25-50%", "50-75%",
"75-100%")
  )

View(df_plot)
head(df_plot)
ggplot(df_plot, aes(x = Gene_stable_ID_rank, y =
Tasmanian_devil_paralogue_gene_stable_ID_rank)) +
  geom_point(aes(
```

```
    color = TOGA_status,
    size = identity_bin
  ), alpha = 0.3) +
  scale_color_manual(values = c("skyblue", "blue", "green", "purple", "orange",
"pink", "yellow")) +  # Manual color scale
  labs(
    x = "Gene stable ID (rank by GC content)",
    y = "Tasmanian devil paralogue transcript ID (rank by GC content)",
    color = "TOGA status",
    size = "% Identity (Target Tasmanian devil gene vs Paralog)"
  ) +
  theme_bw()

gene_rank<-ggplot(df_plot, aes(x = Gene_stable_ID_rank, y =
Tasmanian_devil_paralogue_gene_stable_ID_rank)) +
  geom_point(aes(
    color = TOGA_status,
    size = identity_bin,
  ), alpha = 0.3) +
  labs(
    x = "Gene stable ID (rank by GC content)",
    y = "Tasmanian devil paralogue transcript ID (rank by GC content)",
    color = "TOGA status",
    size = "% Identity (Target Tasmanian devil gene vs Paralog)"
  ) +
  facet_wrap(~ TOGA_status) +
  theme_classic2()+
  theme(legend.position = "bottom")


##### for all####
all_label_df <- df_plot %>%
  group_by(Gene_name) %>%
  slice_max(order_by =
Paralogue_.id._target_Tasmanian_devil_gene_identical_to_query_gene, n = 1) %>%
  ungroup()
library(ggplot2)
library(ggpubr)

All_with_label<-ggplot(df_plot, aes(x = Gene_stable_ID_rank, y =
Tasmanian_devil_paralogue_gene_stable_ID_rank)) +
  geom_point(aes(
    color = TOGA_status,
    shape = identity_bin
  ), alpha = 0.3) +
  geom_text(
    data = all_label_df,
    aes(label = Gene_name),
    size = 2.5,hjust=1.5,
    check_overlap = TRUE  # avoids overlapping labels
  ) +
  labs(
    x = "Gene stable ID (rank by GC content)",
    y = "Tasmanian devil paralogue transcript ID (rank by GC content)",
    color = "TOGA status",
    size = "% Identity (Target Tasmanian devil gene vs Paralog)"
  ) +
  theme_classic2() +
  facet_wrap(~ TOGA_status) +
  theme(legend.position = "bottom")

png("All_class.gene_rank.plot.png", units="in", width=16, height=9, res=900)
All_with_label
```

```
dev.off()



### with shapes
All_with_label <- ggplot(df_plot, aes(x = Gene_stable_ID_rank, y =
Tasmanian_devil_paralogue_gene_stable_ID_rank)) +
  geom_point(aes(
    shape = identity_bin
  ), alpha = 0.6, size = 2.5) +
  geom_text(
    data = all_label_df,
    aes(label = Gene_name),
    size = 2.5,
    hjust = 1.5,
    check_overlap = TRUE
  ) +
  labs(
    x = "Gene stable ID (rank by GC content)",
    y = "Tasmanian devil paralogue transcript ID (rank by GC content)",
    shape = "% Identity Bin"
  ) +
  scale_shape_manual(values = c(4, 2, 8, 20)) +  # adjust as per number of
identity_bin levels
  theme_classic2() +
  facet_wrap(~ TOGA_status) +
  theme(legend.position = "bottom")
All_with_label <- ggplot(df_plot, aes(x = Gene_stable_ID_rank, y =
Tasmanian_devil_paralogue_gene_stable_ID_rank)) +
  geom_point(aes(
    shape = identity_bin,
    fill = identity_bin  # use fill instead of color
  ), color = "black", alpha = 0.6, size = 2.5) +
  geom_text(
    data = all_label_df,
    aes(label = Gene_name),
    size = 2.5,
    hjust = 1.5,
    check_overlap = TRUE
  ) +
  labs(
    x = "Gene stable ID (rank by GC content)",
    y = "Tasmanian devil paralogue gene stable ID (rank by GC content)",
    shape = "% Identity Bin",
    fill = "% Identity Bin"
  ) +
  scale_shape_manual(values = c(4, 1, 21, 24)) +  # fillable shapes
  theme_classic2() +
  facet_wrap(~ TOGA_status) +
  theme(legend.position = "bottom")
All_with_label

library(ggplot2)
library(dplyr)

# Get unique TOGA_status values
toga_levels <- unique(df_plot$TOGA_status)

# Loop through each TOGA_status and save individual facet plots
for (status in toga_levels) {
  df_subset <- df_plot %>% filter(TOGA_status == status)
  label_subset <- all_label_df %>% filter(TOGA_status == status)
```

```
  p <- ggplot(df_subset, aes(x = Gene_stable_ID_rank, y =
Tasmanian_devil_paralogue_gene_stable_ID_rank)) +
    geom_point(aes(
      shape = identity_bin,
      fill = identity_bin
    ), color = "black", alpha = 0.6, size = 2.5) +
    geom_text(
      data = label_subset,
      aes(label = Gene_name),
      size = 2.5,
      hjust = 1.5,
      check_overlap = TRUE
    ) +
    labs(
      x = "Gene stable ID (rank by GC content)",
      y = "Tasmanian devil paralogue gene stable ID (rank by GC content)",
      shape = "% Identity Bin",
      fill = "% Identity Bin",
      title = paste("TOGA Status:", status)
    ) +
    scale_shape_manual(values = c(4, 1, 21, 24)) +
    theme_classic2() +
    theme(legend.position = "bottom")

  # Save the plot
  ggsave(filename = paste0("facet_", gsub("[^A-Za-z0-9]", "_", status), ".png"),
         plot = p, width = 16, height = 9, units = "in", dpi = 300)
}

write.table(df_plot, file = "df_plot.tsv", sep = "\t", row.names = FALSE, quote =
FALSE)
```

### 7. *SAMD9* and *SAMD9L* gene tree

#### a. Sequence Retrieval and Multiple Sequence Alignment (MSA)

Fasta sequences for *SAMD9* and *SAMD9L* genes were retrieved from TOGA output and merged into a single multi-fasta file. The header for each sequence contained the gene and species name to maintain traceability. This multi-fasta file was used to construct multiple sequence alignments using PRANK v.170427 [30] implemented through GUIDANCE2 v2.01 [51], with the alignment mode set to codons.

#### b. Construction of Gene Trees

The maximum likelihood gene trees were constructed using IQ-TREE2 v2.3.6 [52,53] with the following parameters: -m MFP --alrt 1000 -B 1000 --boot-trees, enabling model selection, SH-aLRT branch support, and ultrafast bootstrap approximation (see electronic supplementary material, figure S9).

#### c. Sliding Window Tree Analysis (1 kb windows)

To investigate potential local sequence misassemblies between *SAMD9* and *SAMD9L*, the aligned sequences were divided into non-overlapping 1 kb windows using seqkit sliding (-s 1000 -W 1000). Gene trees were generated for each window and visualised using FigTree, with node support values annotated for assessment of local phylogenetic signal (see electronic supplementary material, figure S9-S10).

#### d. Concordance Factor Estimation

Gene trees generated from the full MSA and individual 1 kb windows were used to estimate gene concordance factors (gCFs). The gCF values provide a measure of how consistently each branch in the species tree is supported by the underlying gene trees, allowing evaluation of potential discordance due to misalignment or paralogy.

```
#GENE TREE USING IQ-TREE
```

```
mkdir Gene_Tree_SAMD9-9L
cd Gene_Tree_SAMD9-9L
sed 's/>/>SAMD9_/g' ../SAMD9/SAMD9.fa > SAMD9-SAMD9L.fa
sed 's/>/>SAMD9L_/g' ../SAMD9L/SAMD9L.fa >> SAMD9-SAMD9L.fa
sed -i 's/-//g' SAMD9-SAMD9L.fa
guidance=/home/morpheus/gprc6a/guidance.v2.02/www/Guidance/guidance.pl
perl $guidance --program GUIDANCE --seqFile SAMD9-SAMD9L.fa --msaProgram PRANK --
seqType nuc --outDir SAMD9-SAMD9L.100_PRANK --genCode 1 --bootstraps 100 --
proc_num 48
seqkit sort SAMD9-SAMD9L.100_PRANK/MSA.PRANK.aln.With_Names >SAMD9-
SAMD9L.PRANK.aln
rm -r SAMD9-SAMD9L.100_PRANK

mkdir iqtree2_MF
cp SAMD9-SAMD9L.PRANK.aln iqtree2_MF
cd iqtree2_MF
#One would be more confident if a clade has its SH-aLRT >= 80% and UFboot >= 95%.
/home/morpheus/anaconda3/bin/iqtree2 -T AUTO -s SAMD9-SAMD9L.PRANK.aln -m MFP --
alrt 1000 -B 1000 --boot-trees


##phylogenetic tree for 1kb region and Gene concordance factor (gCF)
mkdir splitMSA_1kb
cp SAMD9-SAMD9L.PRANK.aln splitMSA_1kb
cd splitMSA_1kb
seqkit sliding -s 1000 -W 1000 -o output_chunks.fasta SAMD9-SAMD9L.PRANK.aln
/home/morpheus/anaconda3/bin/iqtree2 -T AUTO -s SAMD9-SAMD9L.PRANK.aln -m MFP --
alrt 1000 -B 1000 --boot-trees
grep "SAMD9L_Dasyurus_viverrinus_sliding" output_chunks.fasta|cut -f2 -d':' >
window.lst
seqtk seq output_chunks.fasta > SAMD9-SAMD9L.1kb_window.aln
for i in `cat window.lst`
do
grep "$i" SAMD9-SAMD9L.1kb_window.aln |sed 's/>//g' > "$i".lst
seqtk subseq SAMD9-SAMD9L.1kb_window.aln "$i".lst  > SAMD9-
SAMD9L.1kb_window."$i".aln
sed -i 's/_sliding:.*//' SAMD9-SAMD9L.1kb_window."$i".aln
/home/morpheus/anaconda3/bin/iqtree2 -T AUTO -s SAMD9-SAMD9L.1kb_window."$i".aln -
m MFP --alrt 1000 -B 1000 --boot-trees
#Gene concordance factor (gCF)
/home/morpheus/anaconda3/bin/iqtree2 -t SAMD9-SAMD9L.PRANK.aln.treefile --gcf
SAMD9-SAMD9L.1kb_window."$i".aln.treefile --prefix gCF."$i".concord
done
```

### 8. Correlation between *SAMD9* gene loss and dietary changes

#### a. Inference of the gene *SAMD9/9L* gene status

*SAMD9/9L* gene status (intact or loss) was determined using TOGA [5]. Briefly, we used *SAMD9*, *SAMD9L* and *CDK6* gene sequences to extract scaffolds/chromosomes containing these genes from the genome assemblies of mammalian species. The genome assemblies were downloaded using NCBI datasets. The scaffold/chromosome fasta sequence used as query to create a genomic alignment in chain format between human SAMD9/9L chromosome 7 (NC_000007.14) as reference in *make_chains.py* of the make_lastz_chains pipeline. Next, we used chain and *SAMD9* gene annotation of human (NM_017654.4) in BED12 format to asses gene status in other species using TOGA.

```
#chain.sh

#!/bin/bash

counter=0
```

```
for j in *.CDK6_SAMD9_SAMD9L.fa; do
    (
        # Clean up FASTA headers and filename
        sed -i 's/ .*//;s/\./_/' "$j"
        query_name=$(echo "$j" | awk -F'.' '{print $1}')

        # Run the chain generation script
        /mnt/disk4/BUDDHA/tools/make_lastz_chains/make_chains.py Homo_sapiens
"$query_name" Homo_sapiens_chr7.fa "$j" --executor_queuesize 16 --project_dir
Chain-"$query_name"

        # Clean up unnecessary temp files
        cd Chain-"$query_name"
        rm -fr TEMP_run.lastz/ TEMP_run.cat/ TEMP_run.fillChain/ TEMP_pslParts/
TEMP_axtChain/ TEMP_psl/ cleanUp.csh DEF make_chains.log
make_chains_py_params.json master_script.sh
        cd ..
    ) &

    ((counter++))
    if (( counter % 15 == 0 )); then
        wait  # Wait for 15 jobs to finish
    fi
done

wait  # Wait for any remaining jobs

#TOGA
mkdir -p /mnt/disk4/BUDDHA/SAMD9-9L/Diet_relation/sorted/mutation_plot/
for i in `ls -1 -d Chain-*`
do
cd $i
query_name=$(echo "$i" | awk -F'-' '{print $2}')
/mnt/disk4/BUDDHA/TOGA-1.1.14/toga.py
Homo_sapiens."$query_name".allfilled.chain.gz /mnt/disk4/BUDDHA/SAMD9-
9L/Homo_sapiens_SAMD9.bed12 Homo_sapiens.2bit "$query_name".2bit --pn TOGA-
"$query_name" --nc /mnt/disk4/BUDDHA/TOGA-1.1.14/nextflow_config_files/ --
cesar_bigmem_config /mnt/disk4/BUDDHA/TOGA-
1.1.14/nextflow_config_files/cesar_bigmem_config.nf --cesar_jobs_num 500 --
cesar_buckets 3,5,25,50 --ces --kt --chain_jobs_num 60
/mnt/disk4/BUDDHA/TOGA-1.1.14/supply/plot_mutations.py --publication_mode_heni
/mnt/disk4/BUDDHA/SAMD9-9L/Homo_sapiens_SAMD9.bed12 TOGA-
"$query_name"/inact_mut_data.txt NM_017654.4 /mnt/disk4/BUDDHA/SAMD9-
9L/Diet_relation/sorted/mutation_plot/"$query_name".svg

cat TOGA-"$query_name"/inact_mut_data.txt| sed "s/NM_017654.4/$query_name/g" >>
/mnt/disk4/BUDDHA/SAMD9-9L/Diet_relation/sorted/SAMD9.inact_mut_data.txt
orthology_classification=`cat TOGA-"$query_name"/orthology_classification.tsv|tail
-1|cut -f5`
loss_summ=`cat TOGA-"$query_name"/loss_summ_data.tsv|tail -1|cut -f3`
echo -e "$query_name\t$orthology_classification\t$loss_summ" >>
/mnt/disk4/BUDDHA/SAMD9-9L/Diet_relation/sorted/SAMD9.orthology.loss_summ.tsv
cat TOGA-"$query_name"/codon.fasta |tail -2 |sed 's/ //g;s/X/N/g'|sed
"s/>.*/>$query_name/g" >> /mnt/disk4/BUDDHA/SAMD9-
9L/Diet_relation/sorted/"$clade".codon.fasta
cat TOGA-"$query_name"/nucleotide.fasta |tail -2 |sed 's/ //g'|sed
"s/>.*/>$query_name/g" >> /mnt/disk4/BUDDHA/SAMD9-
9L/Diet_relation/sorted/"$clade".nucleotide.fasta
cd /mnt/disk4/BUDDHA/SAMD9-9L/Diet_relation/
done
```

```
mv sorted SAMD9_sorted
cd SAMD9_sorted
echo -e "species_name,SAMD9_gene_status" > SAMD9.TOGA_inferred_gene_status.csv
awk '$3=="I" || $3=="L"' SAMD9.orthology.loss_summ.tsv |cut -f1,3|sed 's/\t/,/g'
>> SAMD9.TOGA_inferred_gene_status.csv
cut -f1 -d',' SAMD9.TOGA_inferred_gene_status.csv > Species.lst

head -n 1 Trehalase_data.csv | awk '{print "Species_name", "SAMD9_gene_status",
$0}'|sed 's/ /,/g' > final_output.csv

#EXTRACT TOGA INFO
for i in `tail -n+2 Species.lst`
do
sp=`grep -w "$i" SAMD9.TOGA_inferred_gene_status.csv|cut -f1 -d','`
SAMD9_status=`grep -w "$i" SAMD9.TOGA_inferred_gene_status.csv|cut -f2 -d','`
info=`grep -w "$i" Trehalase_data.csv`
echo -e "$sp,$SAMD9_status,$info" >> final_output.csv
echo $sp
done
```

**b. Gathering of diet info for mammals**

The Low-quality annotation of *SAMD9* in 7 out of 7 carnivore species indicates the possible link between *SAMD9* gene loss and carnivores' diet. Therefore, we compiled the dietary information. We were able to retrieve the proportion of endothermic vertebrates for a total of 74 mammal species whose *SAMD9* gene status is also available from Jiao, Hengwu et al. [16] and Kapsetaki, Stefania E et al. [54]. To examine whether *SAMD9* gene loss is associated with the percentage of endothermic vertebrates in the diet, we performed phylogenetic logistic regression analyses using the R package "phylolm" [55,56]. Specifically, we tested whether the proportion of endothermic vertebrates in the diet of each species could predict the retention (coded as 1) or loss (coded as 0) of *SAMD9*. The phylogenetic logistic regression analyses used logistic_MPLE and logistic_IG10 [55,56].

To assess the robustness of the phylogenetic logistic regression results, we repeated the analysis five times using different values for set.seed(). Although the exact p-values and slopes varied slightly across runs, the overall pattern was consistent: the logistic_IG10 method consistently produced statistically significant p-values (< 0.01), while the logistic_MPLE method yielded non-significant results in all runs. The results presented in figure 2b correspond to the run with set.seed(35842). Figures from the other runs are available on our GitHub repository. We used 2000 bootstraps to determine the 95% confidence intervals.
