## Supplementary Figures for "Illuminating the mystery of thylacine extinction: a role for relaxed selection and gene loss"

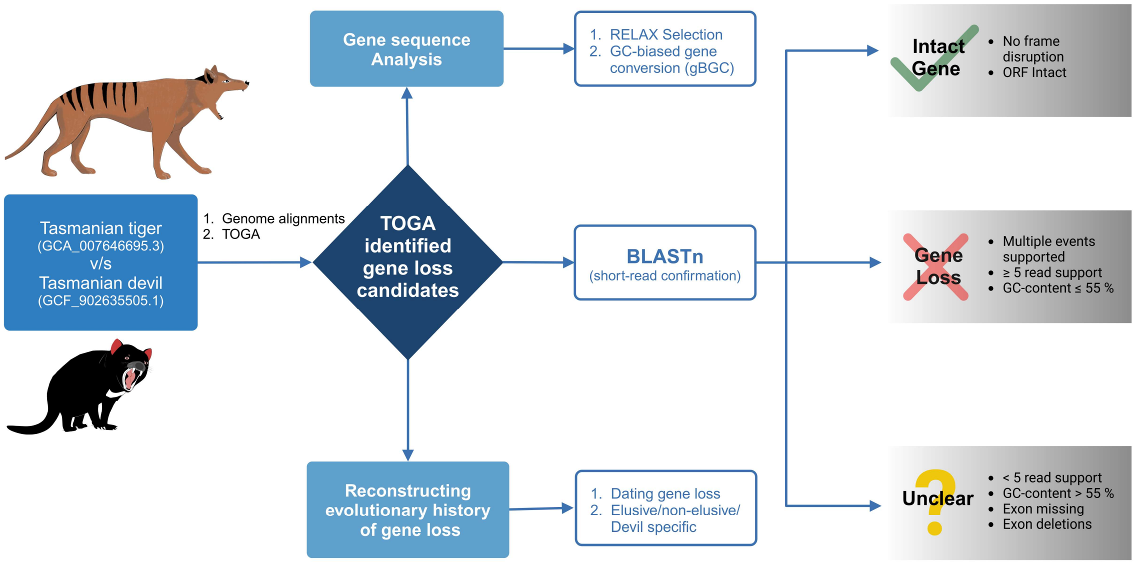


**figure S1.** Workflow for identifying gene loss candidates using TOGA and confirming them using BLASTn of short-read data. The criteria for classifying TOGA-identified "clearly lost" gene loss candidates include GC content, exon missing, and/or deletion. Molecular evolutionary analysis (gBGC and Relax) was conducted along with dating gene loss.

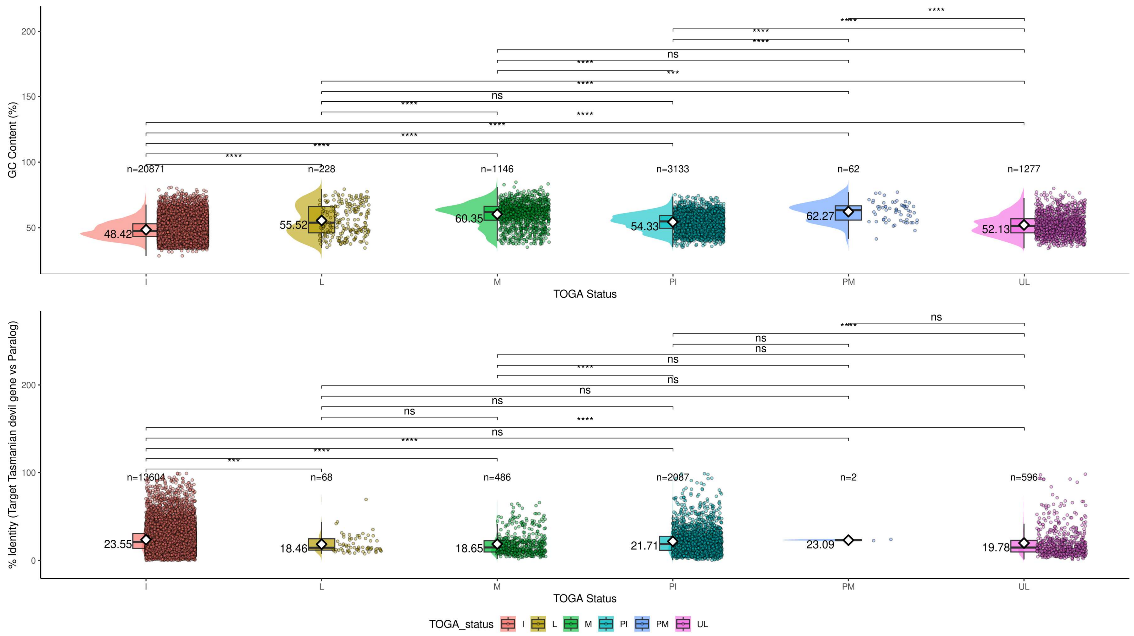

**figure S2.** Effect of GC-content and paralogs on TOGA classification. The top panel shows the pairwise Wilcoxson test, which shows that genes classified as intact (I) by TOGA have a significantly lower GC content (on y-axis) than those identified as lost (L), missing (M), partially intact (PI), partially missing (PM) and unclear loss (UL). The bottom panel shows the effect of sequence identity between Tasmanian devil and paralog (in %) on TOGA classification. The rain cloud plot shows the distribution of data points (along with density) and a boxplot. The white diamond represents the mean GC-content or % identity between paralogs. The sample size is given above each TOGA classification. The ggplot2 [8] was used to generate the plot.

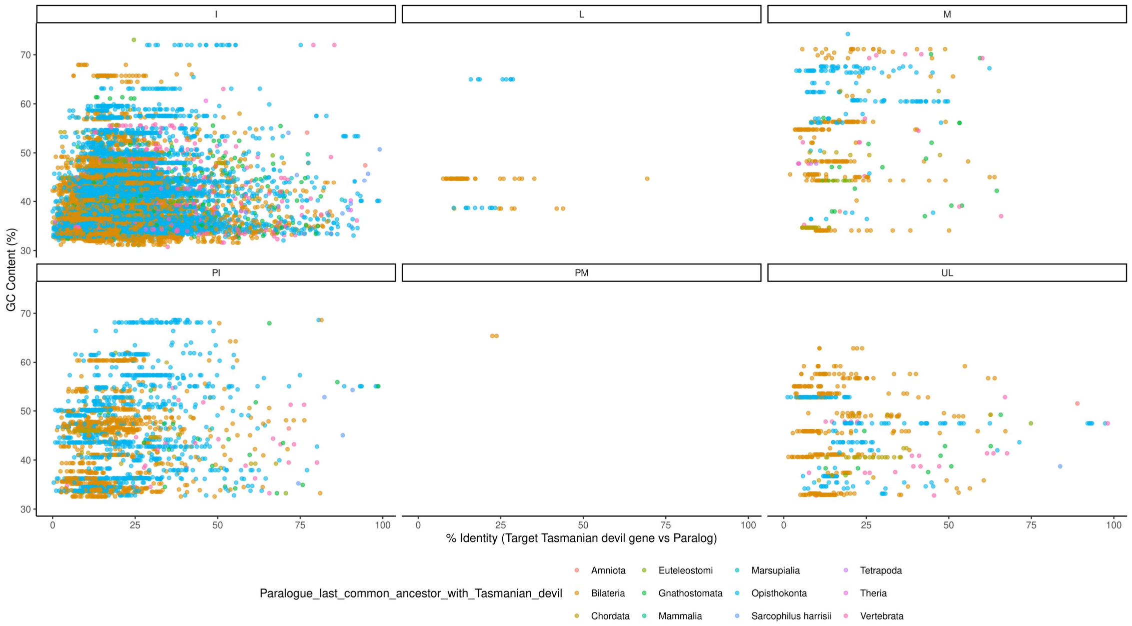


**figure S3.** Effect of GC content and paralogy on gene classification by TOGA. Scatterplot showing the relationship between GC content (y-axis) and sequence identity (%) between Tasmanian devil gene stable IDs and their corresponding paralogs (x-axis). Each point represents a gene–paralog pair, coloured by the last common ancestor of the paralog. The plot is faceted by TOGA classification: intact (I), lost (L), missing (M), partially intact (PI), partially missing (PM), and unclear loss (UL). Genes classified as PI and PM exhibit high GC content and high sequence identity to their paralogs, suggesting that elevated GC content and sequence similarity may contribute to ambiguous gene classification.

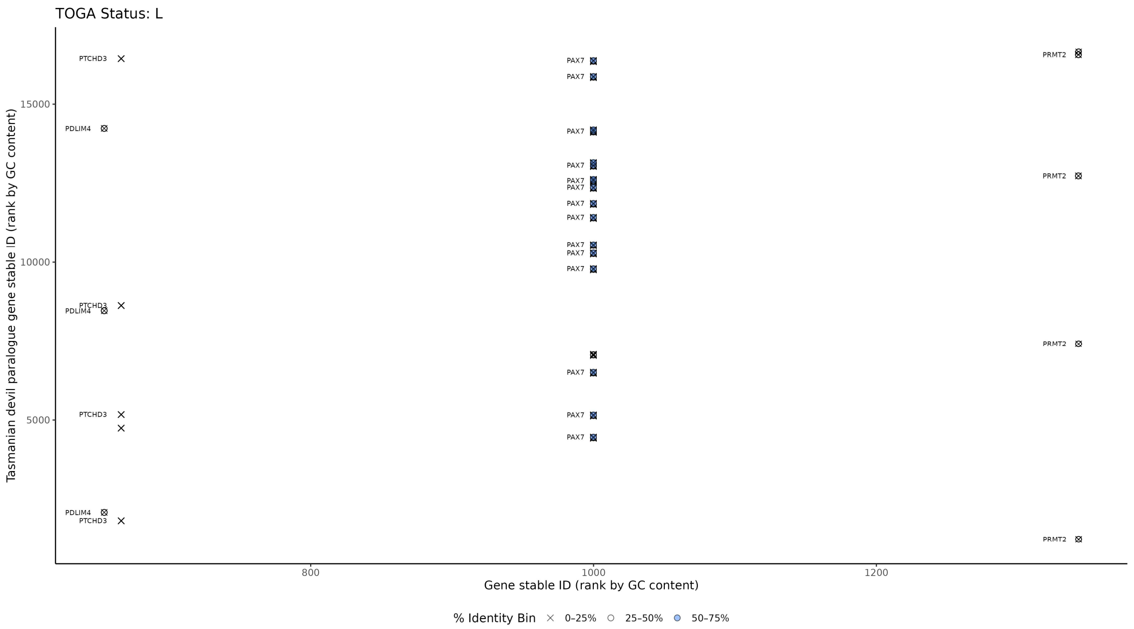


**figure S4.** Effect of GC content and paralogy on false or unclear gene loss classification. Genes classified as clearly lost (L) by TOGA—*PDLIM4*, *PTCHD3*, *PAX7*, and *PRMT2*—are visualised based on their GC content and paralogy. The genes (x-axis) and their corresponding paralogs (y-axis) are ranked by GC content. The shapes indicate the percentage identity between each gene and its paralog. Notably, *PRMT2* exhibits high GC content (66.14 %), while *PAX7* has multiple paralogs with sequence identity exceeding 50%. These patterns suggest that high GC content and the presence of highly similar paralogs may contribute to false or ambiguous gene loss events.

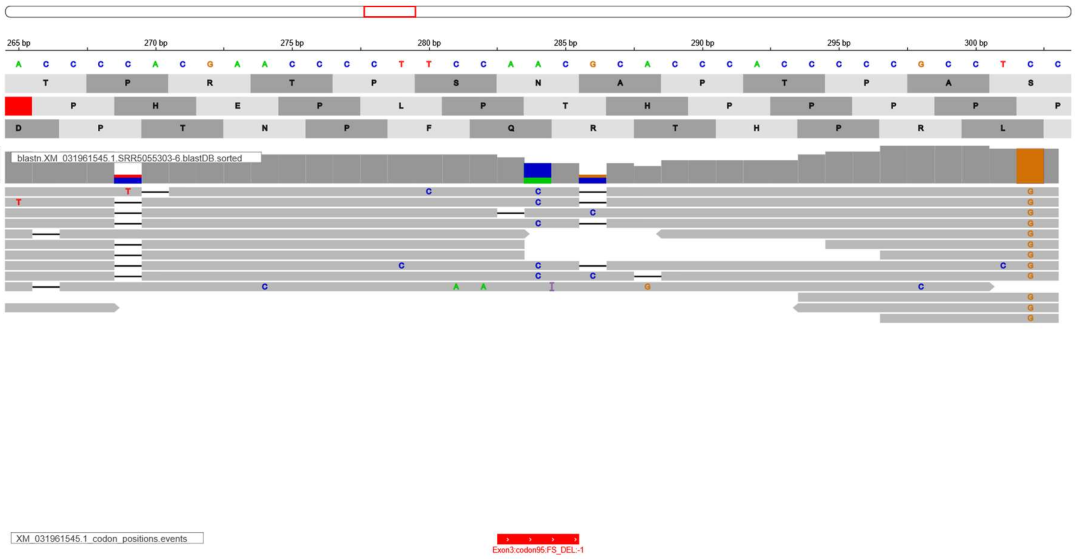

**figure S5.** Ambiguity in single-base deletion detection due to paralogous sequences. TOGA identified a single-base deletion event in the *PAX7* gene; however, a short-read-based BLASTn search revealed multiple reads aligning to paralogous sequences. These overlapping hits from paralogs introduce ambiguity in interpreting the deletion event, suggesting that the observed signal may result from misalignment or mapping artefacts rather than a true deletion.

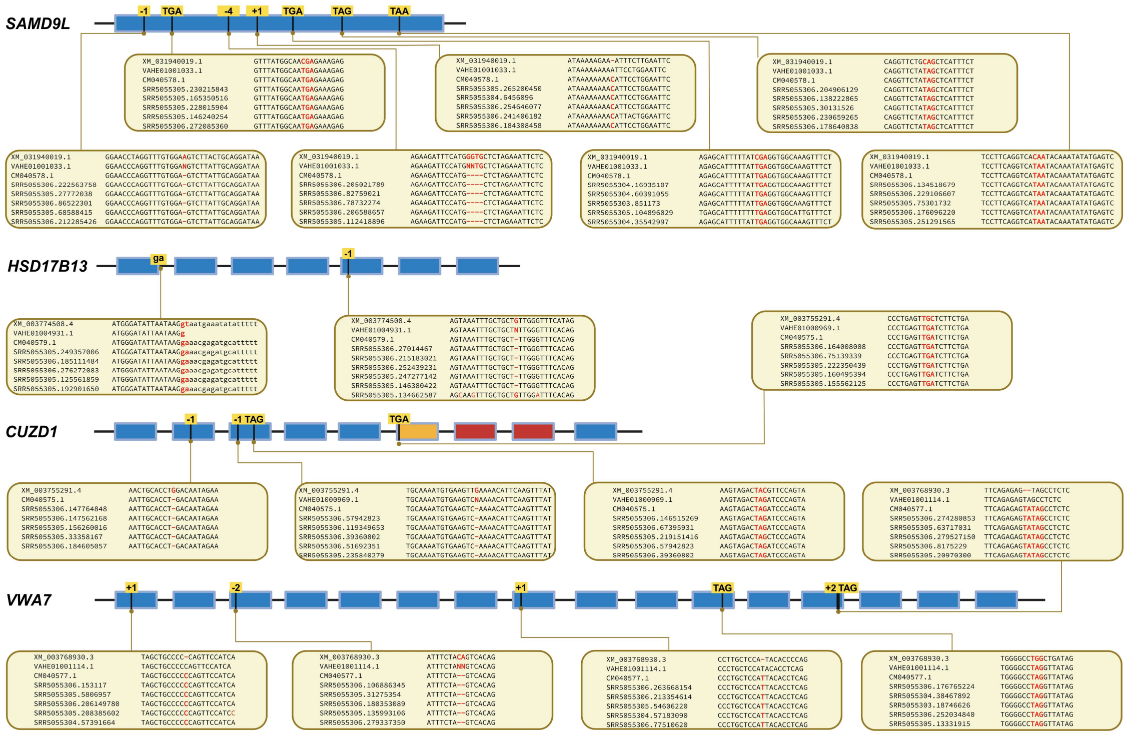


**figure S6.** Frame-disrupting events leading to gene inactivation. Short-read support for frame-disrupting changes and/or in-frame stop codons, resulting in the inactivation of *SAMD9L*, *HSD17B13*, *CUZD1*, and *VWA7* genes. The blue boxes represent intact exons, while exons affected by frame-disrupting changes are highlighted in yellow. Exons with no blastn hits are shown in red, and those with partial blastn hits are depicted in orange. The inset at each event, shown in the comment box, depicts raw read support for that event visualised using MView [1]. The first row represents the transcript ID (starting with XM) of the focal gene in the Tasmanian devil. The second row shows blastn hits in the thylacine scaffold-level assembly (GCA_007646695.1; starting with VAHE). The third row shows blastn hits in the thylacine chromosomal-level genome (GCA_007646695.3; starting with CM). The bottom rows display short-read support from thylacine SRA data (starting with SRR).

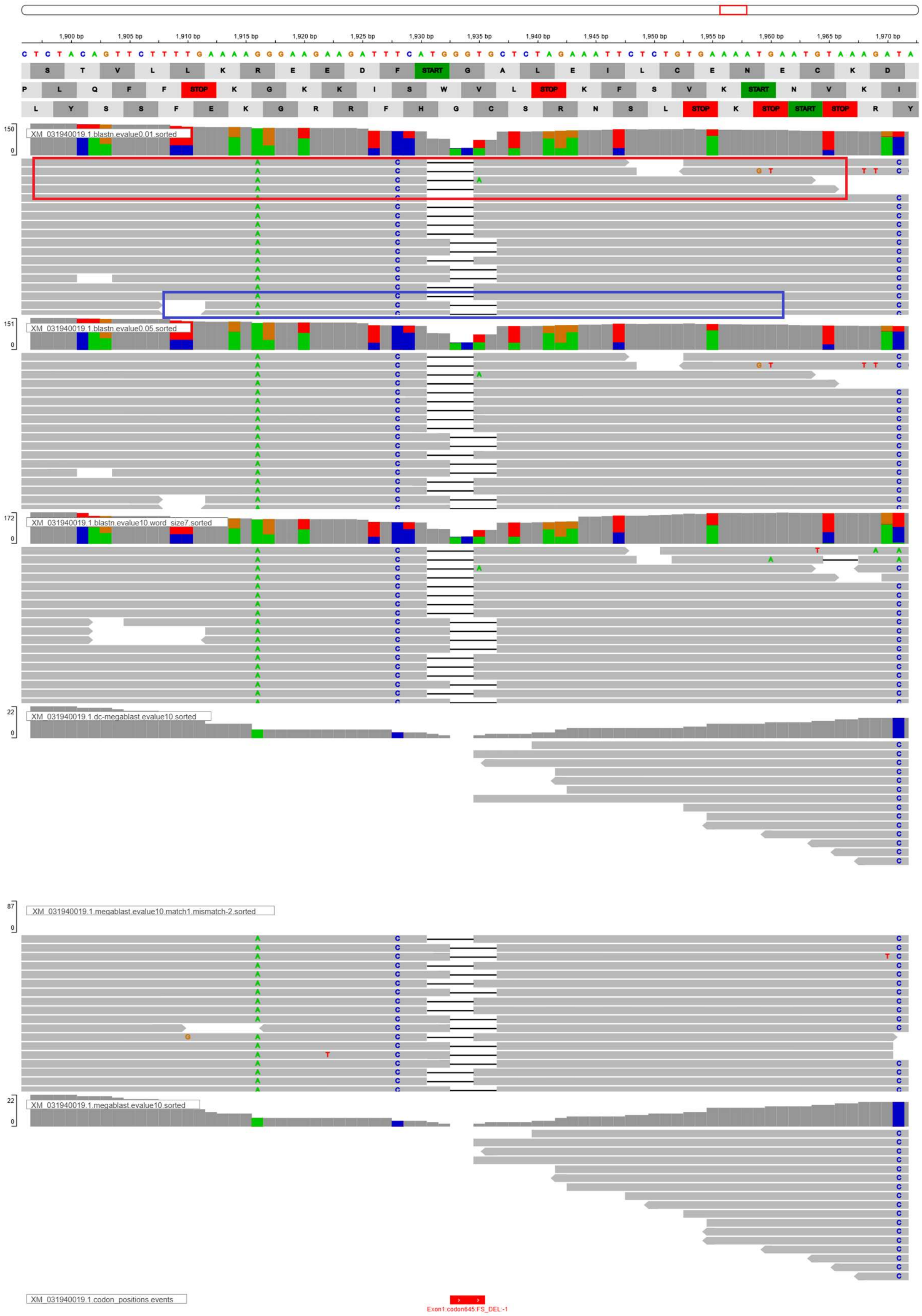

**figure S7.** Evidence for four-base deletion by BLASTn-based approach (TGGG/GGTG). We performed BLASTn search against the short read database of thylacine using Tasmanian devil *SAMD9L* gene sequence as query with different BLASTn parameters [2] (i.e. -task blastn/megablast/dc-megablast -evalue 0.01/0.05/10 -word_size 7/11 -match 1 and mismatch

-2, output saved in output format – '17 SQ') used to create an IGV report [3,4]. The red square box indicating reads aligned more toward the left shows the deletion of TGGG, while reads aligned toward the right side result in the deletion of GGTG (shown with a blue box).

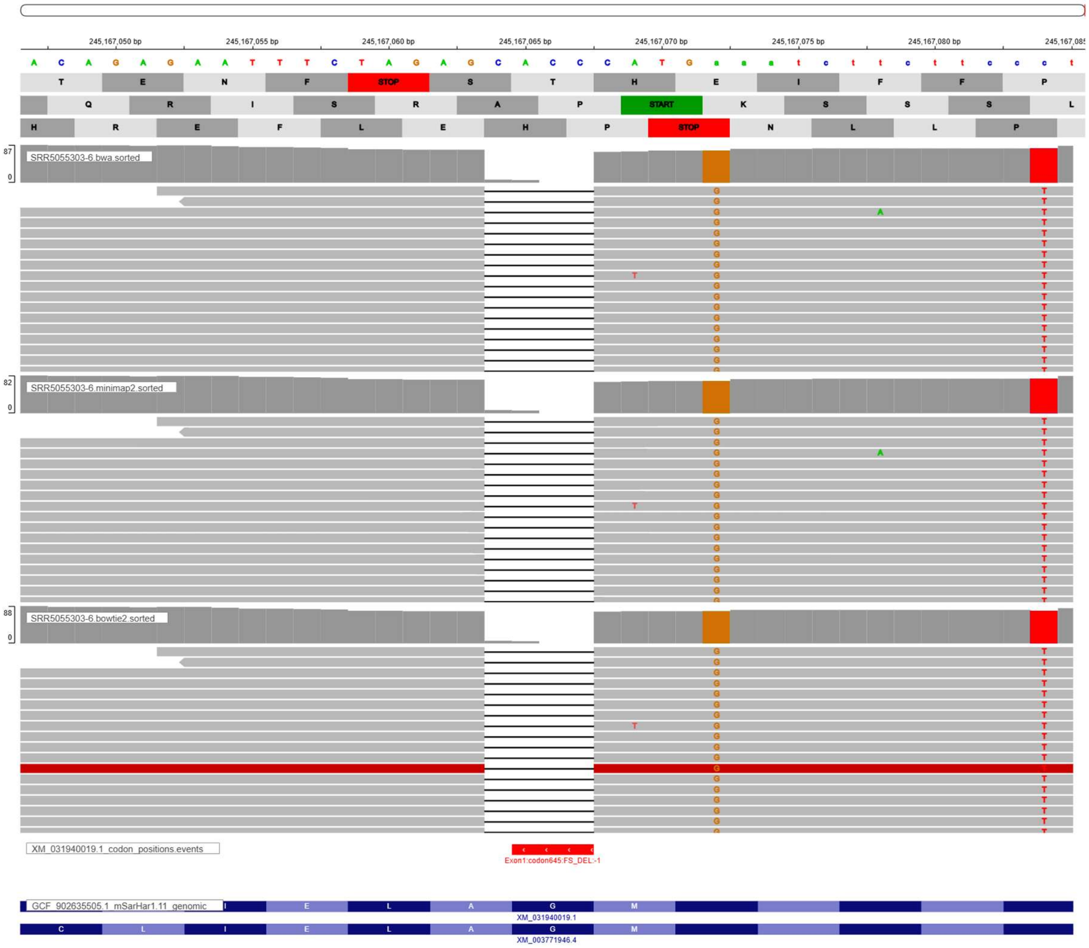


**figure S8.** Evidence for four-base deletion (GGTG) in *SAMD9L* of thylacine using a mapping-based approach. We performed cross-species mapping of thylacine short-read data SRR5055303, SRR5055304, SRR5055305 and SRR5055306 to Tasmanian devil using BWA-MEM v 0.7.17 [5], minimap2 v2.17 [6]and bowtie2 v2.5.3 [7]. We found consistent deletion of GGTG with all three read mappers. The IGV reports were used to generate the plot [4].

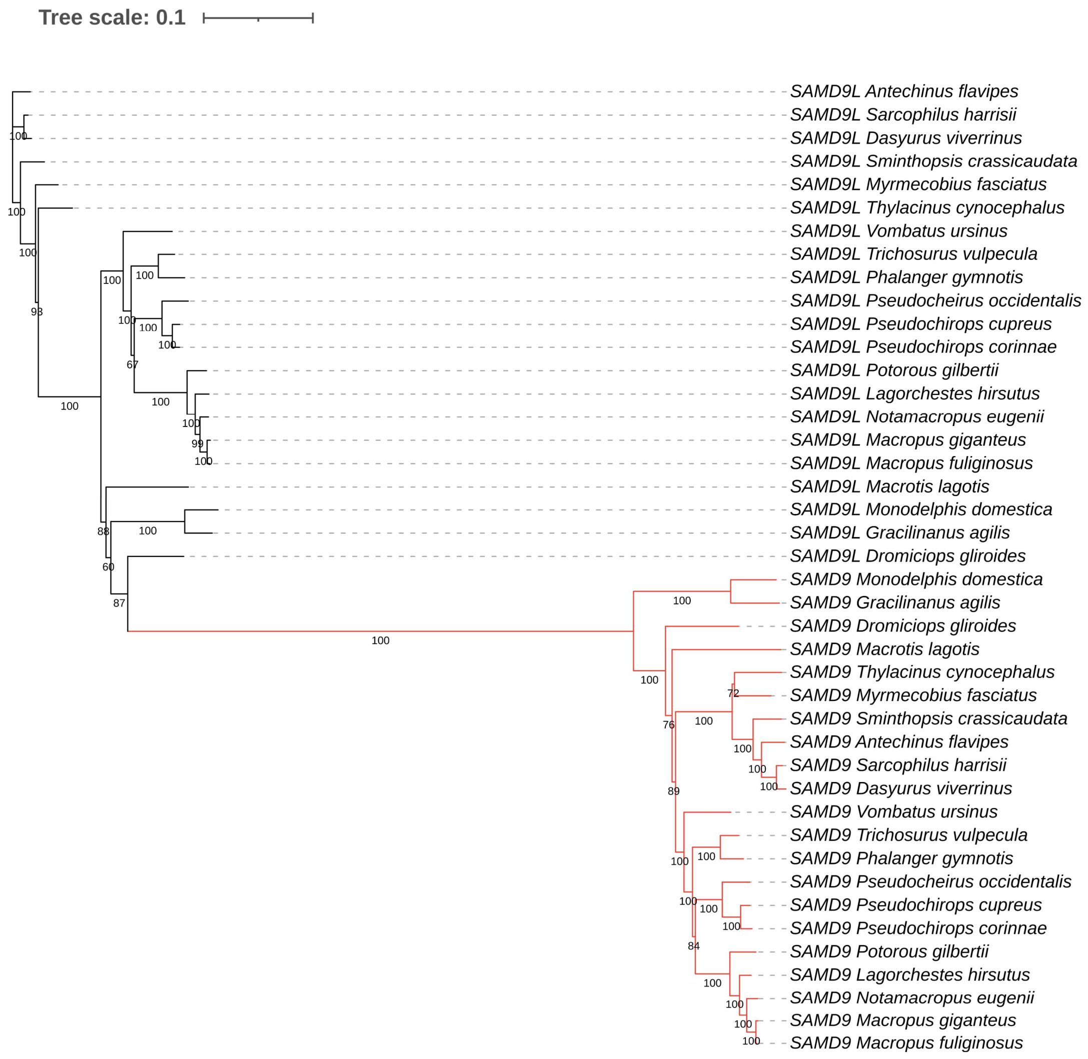


**figure S9.** Gene tree of *SAMD9*/*SAMD9L*. Gene tree of *SAMD9*/*SAMD9L* constructed using IQ-TREE. Red-coloured branches represent orthologs of *SAMD9* in marsupial species. Bootstrap values are provided at each node.

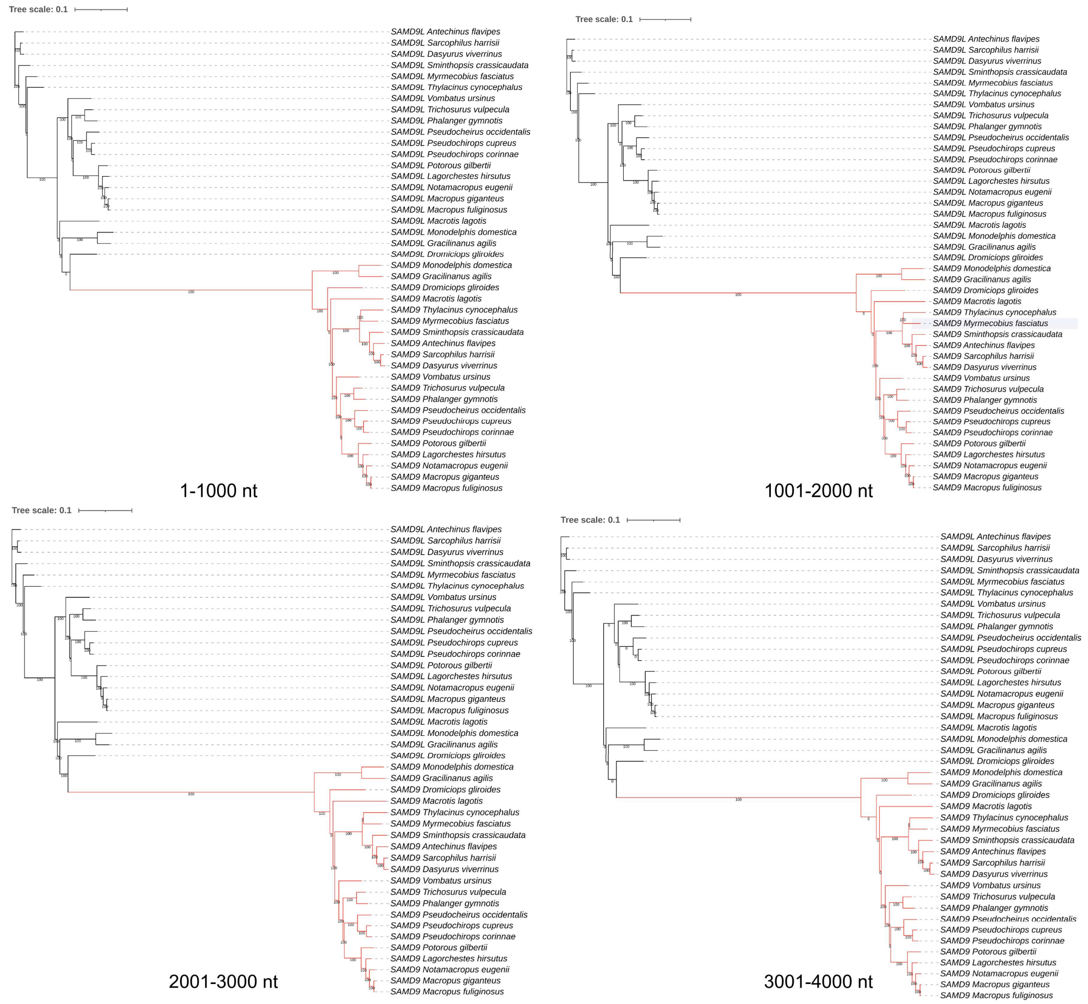


**figure S10.** Gene tree based on 1 kb non-overlapping windows along the multiple sequence alignment of the coding sequence of *SAMD9*/*SAMD9L*. Values at each node indicate the concordance factor with the gene tree derived from the full-length sequence with a gene tree of 1kb sequence. Red-coloured branches represent orthologs of *SAMD9* across marsupial species.

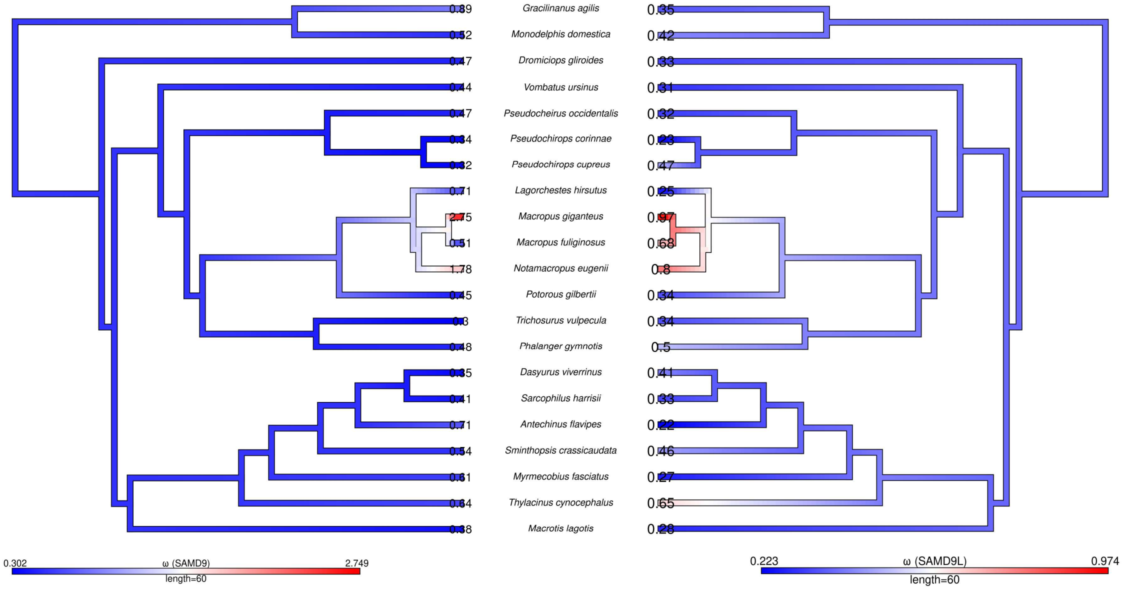


**figure S11.** *SAMD9/9L* gene branch-specific selection evaluation was done using the ω (ω = dN/dS; dN, nonsynonymous substitution rate; dS, synonymous substitution rate), calculated using the codeML program of PAML under Model 1 (branch model) using 1x4 codon frequency model. (tree on the left) ω across the marsupial species of *SAMD9* gene. (tree on the right) ω across the marsupial species of *SAMD9L* orthologs. We used phytools to generate a plot [9].

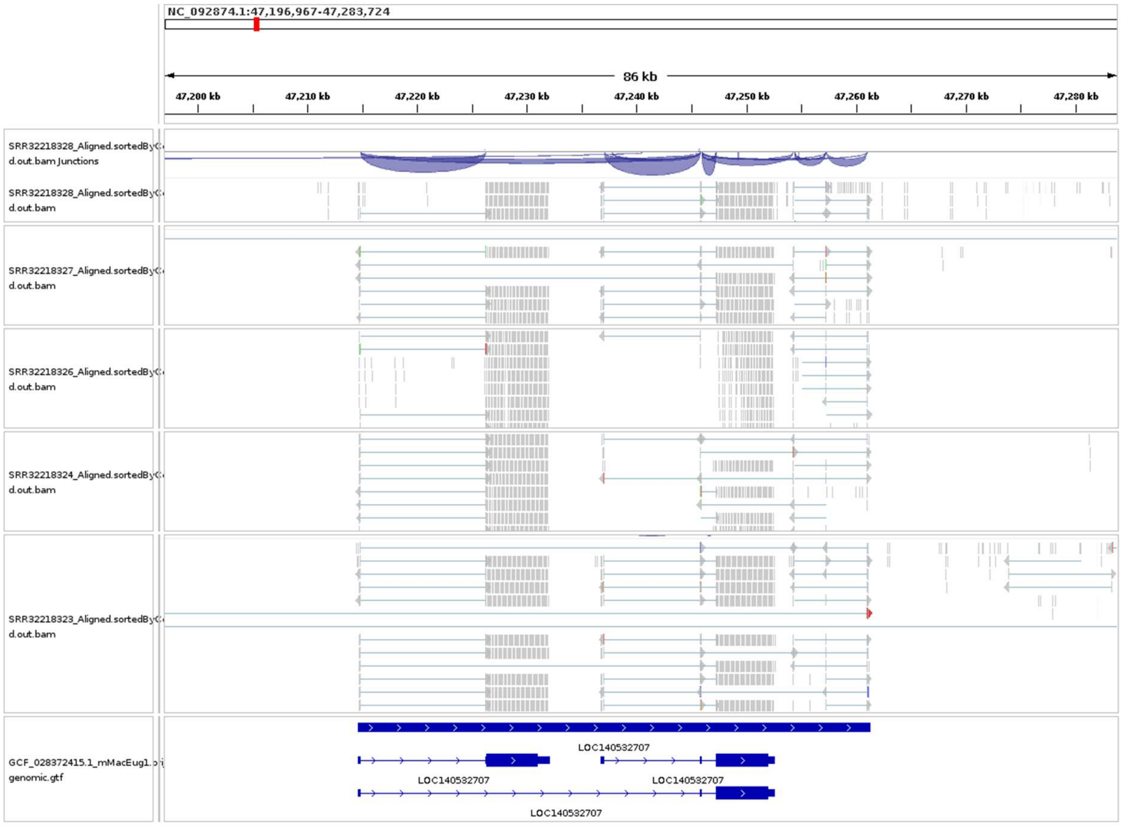


**figure S12.** Expression of *SAMD9* and *SAMD9L* genes in the tammar wallaby (*Notamacropus eugenii*). Short-read RNA-seq data (SRR32218323, SRR32218324, SRR32218325, SRR32218326, SRR32218327, SRR32218328) of the tammar wallaby were mapped to the reference genome (*Notamacropus eugenii*, GCF_028372415.1) using STAR aligner v2.7.1a [10]. The resulting BAM files were visualised in Integrative Genomics Viewer (IGV) [3] along with the corresponding GTF annotation file to assess the transcriptional status of the *SAMD9* and *SAMD9L* genes.

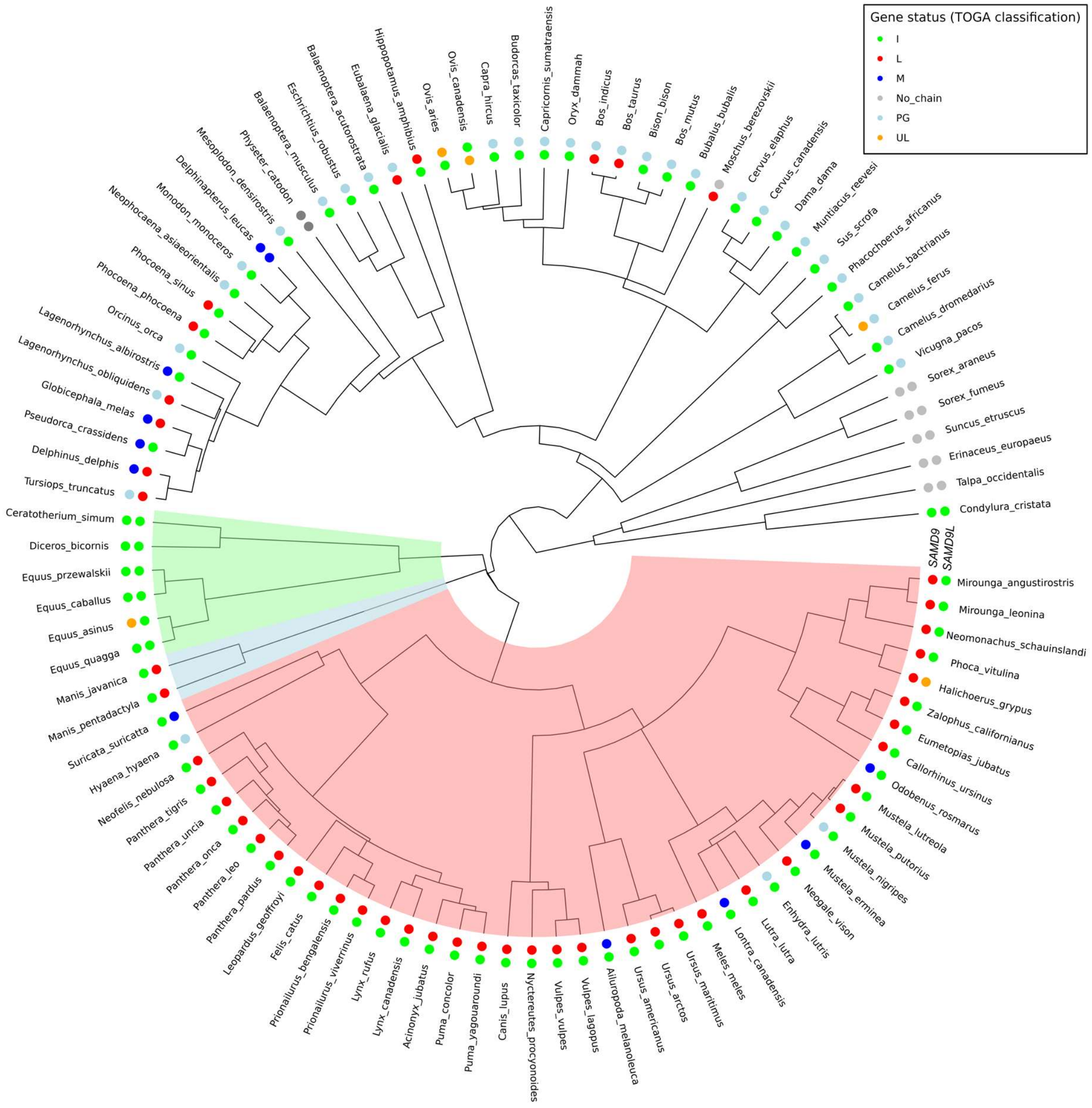


**figure S13.** Phylogenetic tree of mammalian species depicting the gene status of the *SAMD9* and *SAMD9L*. The circular phylogenetic tree illustrates relationships among species, with colours highlighting specific orders (light shades of red, blue, and green colours for Carnivora, Pholidota, and Perissodactyla, respectively). Inner dots represent the *SAMD9* gene status, while outer dots represent the *SAMD9L* gene status. The gene status is marked by coloured dots next to each species name: Green - I (intact); Red - L (clearly lost); Blue - M (missing); Light blue - PG (no orthologous chains identified); Orange - UL (uncertain loss); Grey - No_chain (found no classifiable chains). The gene statuses were derived using TOGA gene classification methods. We used ggtree to generate the plot [11]. The phylogenetic tree was obtained from the Timetree website (https://timetree.org/).

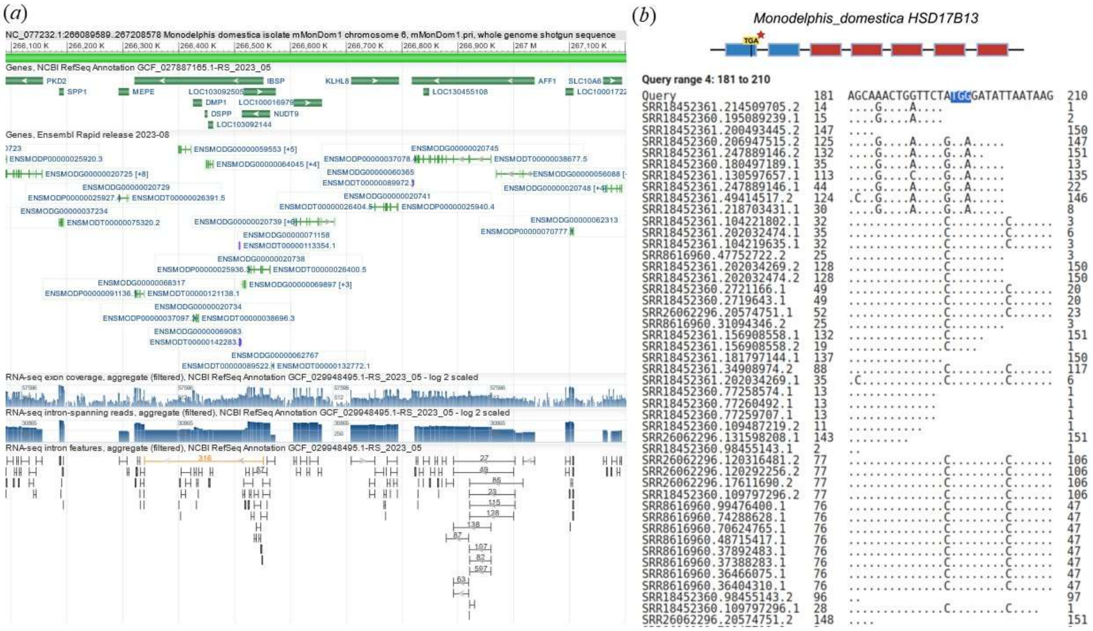

**figure S14.** *HSD17B13* gene loss in grey short-tailed opossum (*Monodelphis domestica*). (a) The NCBI genome data viewer lacks the annotation for *HSD17B13*, while *HSD17B11* is annotated as LOC100016979. (b) Short-read support for the in-frame stop codon, resulting in the inactivation of *HSD17B13* genes. The blue boxes represent intact exons, while exons affected by frame-disrupting changes are highlighted in yellow. Exons with no blastn hits are shown in red. The red star above the in-frame stop codon, i.e., TGA, depicts this event as polymorphic.

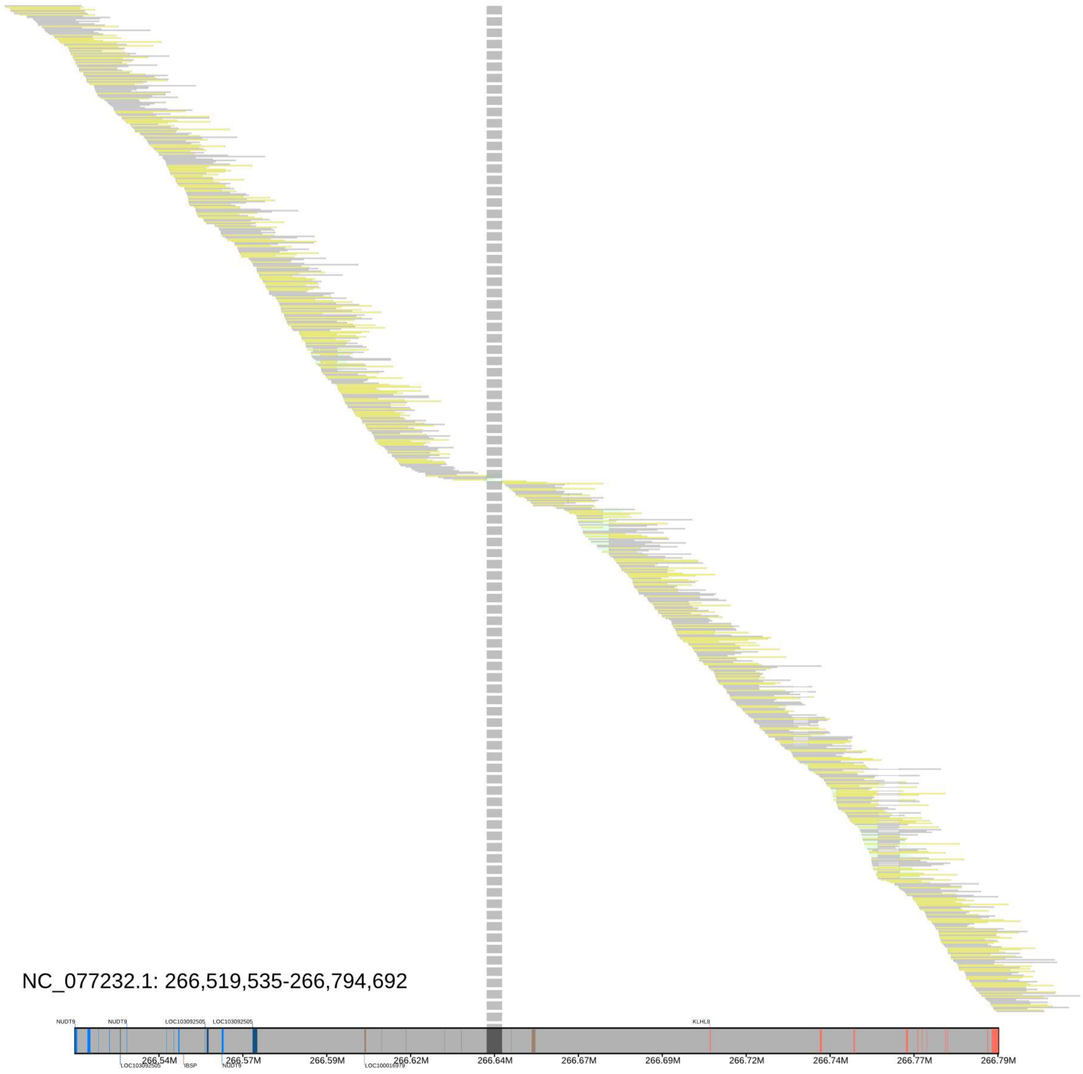


**figure S15.** Assembly gap at *HSD17B13* locus in grey short-tailed opossum (*Monodelphis domestica*). Alignment plot of the *HSD17B13* locus for grey short-tailed opossum (*Monodelphis domestica*). Yellow bars illustrate alignments to the reverse strand, and grey bars above the reference genome represent alignments to the forward strand. Blue and red boxes on the reference genome bar portray Exons of syntenic genes. Vertical dashed lines depict gaps in the assembly. Klumpy was used to generate an alignment plot [12].

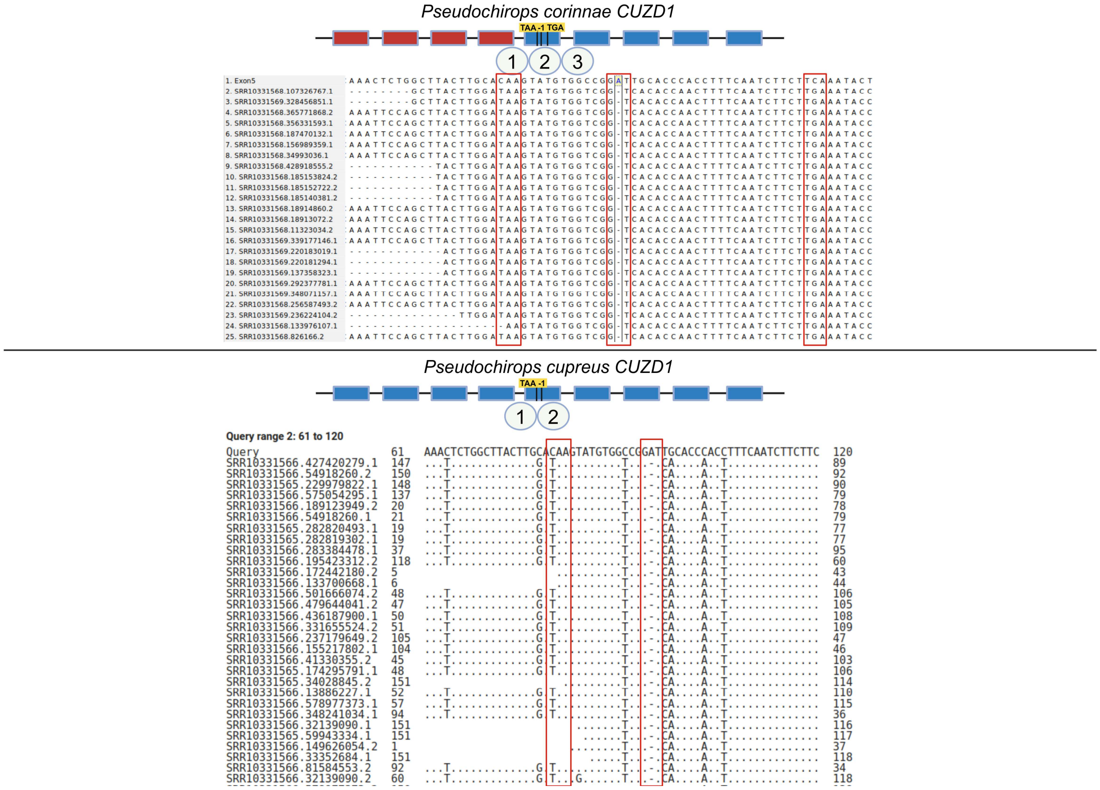


**figure S16.** Frame-disrupting events leading to *CUZD1* gene loss in *Pseudochirops corinnae* and *Pseudochirops cupreus*. The blue boxes represent intact exons, while exons affected by frame-disrupting changes are highlighted in yellow. Exons with no blastn hits are shown in red. Short-read support for frame-disrupting changes and/or in-frame stop codons in Exon5, resulting in the inactivation of *CUZD1* genes from SRR10331568-SRR10331569 and SRR10331565-SRR10331566 of *Pseudochirops corinnae* (top panel) and *Pseudochirops cupreus* (bottom panel), respectively.

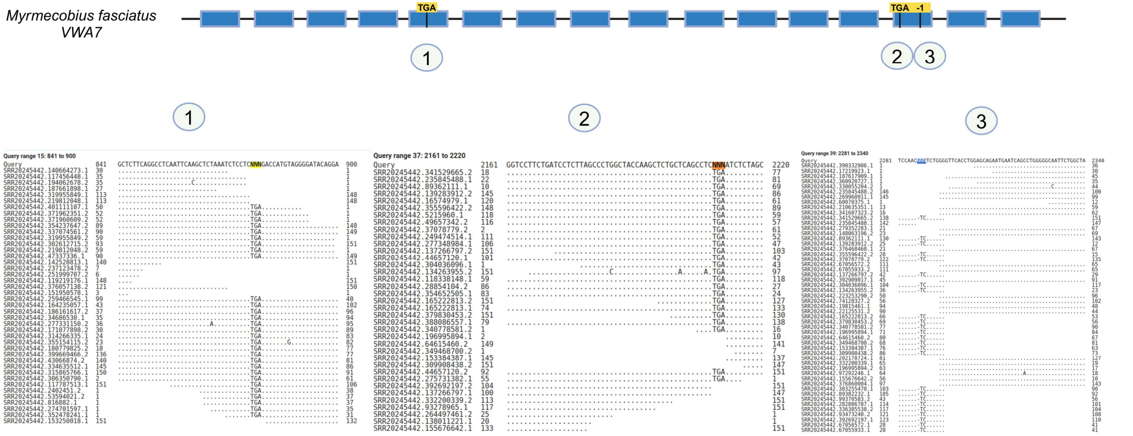


**figure S17.** Frame-disrupting events leading to *VWA7* gene loss in numbat (*Myrmecobius fasciatus*). The blue boxes represent intact exons, while exons affected by frame-disrupting changes are highlighted in yellow. The short-read support is shown for a particular event (numbered accordingly) from SRR20245442 of the whole genome sequencing data of numbat.

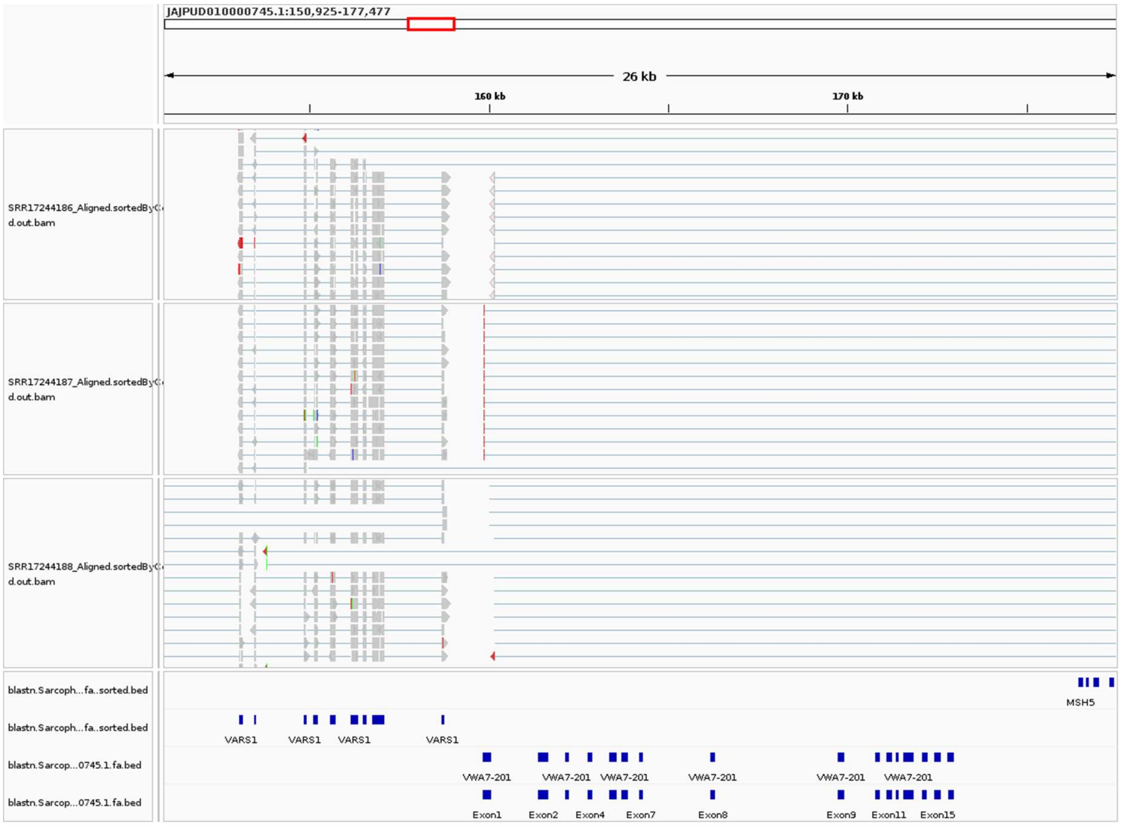


**figure S18.** Transcriptional status at the *VWA7* locus in the numbat (*Myrmecobius fasciatus*). The absence of mapped reads at the inactivated *VWA7* gene locus, contrasted with their presence at the *VARS1* gene (used as a control), supports the inactivation of *VWA7* in the numbat. The bottom blue boxes represent exonic BLASTn hits, with SRR IDs on the left indicating transcriptomic data from the tongue (SRR17244186), lung (SRR17244187), and liver (SRR17244188) of a female numbat.
